# Long-term low-intensity vibration attenuates age-associated adipogenic bias and enhances osteogenic potential in mesenchymal stem cells

**DOI:** 10.64898/2026.11.13.766500

**Authors:** Omar Faruq, Calvin L Jones, Nilson R.S Silva, Scott Birks, Sean Howard, Gunes Uzer

## Abstract

Aging impairs mesenchymal stem cells (MSCs) function and regenerative potential, contributing to a shift in MSCs differentiation away from osteogenesis and toward adipogenesis. This study investigated the effects of in vivo low-intensity vibration (LIV) on bone characteristics, exercise performance, and the functional, mechanical, and molecular phenotypes of bone marrow-derived MSCs from 24-month-old mice. LIV (90 Hz, 0.7 g) was initiated at 5 months and administered twice daily until 24 months. Outcomes were compared with age-matched, non-vibrated controls. LIV was associated with improved voluntary wheel-running performance, including increased running distance and velocity, alongside a trend towards increased trabecular bone volume fraction and trabecular number. MSCs isolated from LIV-treated mice (+LIV) exhibited reduced senescence marker p16 and increased Ki67 proliferation index. +LIV MSCs had increased osteogenic differentiation and mineralization along with upregulated Runx2 and Col1a1 expression. +LIV MSCs exhibited decreased adipogenic differentiation and lipid accumulation, showing downregulated Fasn and Ap2 expression. +LIV MSCs exhibited increased F-actin fiber number, altered vinculin-associated focal adhesions, and increased apparent whole-cell stiffness. Nuclear characterization revealed increased lamin A/C levels and a shift in the storage–loss crossover frequency from 68 to 94 Hz, indicating that elastic-dominant behavior persisted to a higher frequency. These mechanical changes coincided with increased global H3K4me3 and decreased H3K9me3 and H3K27me3 in +LIV MSCs. Collectively, these findings suggest that in vivo LIV is associated with enhanced osteogenic potential in aged MSCs, accompanied by coordinated changes in cytoskeletal organization, nuclear mechanics, and histone modifications, although causal relationships among these changes remain to be established.

## 1. Introduction

Aging is associated with numerous pathophysiological conditions and a gradual decline in the normal function of various organ systems. The progressive loss of bone mass and deterioration of the musculoskeletal system are closely associated with advancing age. **[1]** Mesenchymal Stem Cells (MSCs) are multipotent progenitor cells that play essential roles in tissue homeostasis through their self-renewal capacity, multilineage differentiation potential, and immunomodulatory functions in bone. With advancing age, MSCs exhibit a shift in lineage commitment, characterized by increased adipogenic and reduced osteogenic potential. **[2]** Previous studies focusing on age-related changes in MSCs function have demonstrated that aging profoundly alters the fate and functional properties of MSCs, resulting in impaired proliferative capacity, accumulation of cellular senescence, altered lineage commitment, and metabolic dysfunction. **[3,4]** These age-associated changes in MSCs function may contribute to the diminished regenerative capacity of the musculoskeletal system.

MSCs functional activities are regulated by a combination of biochemical and biophysical cues that act across multiple levels, from the extracellular matrix (ECM) to the intracellular and subcellular compartments. ECM proteins, including collagen, fibronectin, and laminin, facilitate MSCs adhesion and contribute to lineage specification through focal adhesion-mediated signaling. **[5,6]** MSCs are highly sensitive to the mechanical properties of their microenvironment, which serve as important determinants of lineage commitment both *in vitro* and *in vivo.* **[7]** Furthermore, cytoskeletal organization and elasticity, cell–matrix adhesion strength, and cell shape play critical roles in regulating MSCs fate and differentiation. **[8–10]**

Aging is accompanied by progressive alterations in the mechanical properties of both cells and their surrounding microenvironment, which may contribute to the functional decline of MSCs. **[11]** Because aging alters the mechanical phenotype of MSCs, understanding how mechanical cues regulate MSCs fate is particularly important for elucidating age-associated changes in osteogenic and adipogenic differentiation. These mechanical cues are transduced through integrin–focal adhesion complexes, cytoskeletal remodeling, and multiple mechanosensitive signaling pathways. Several studies have investigated the effects of mechanical stimulation on stem cell function and demonstrated that mechanical stimulation can promote osteogenic differentiation. **[12,13]** Interestingly, mechanical stimulation has been shown to influence MSCs lineage specification, suggesting that modulation of the mechanical signals may represent a potential strategy for promoting osteogenesis and counteracting age-associated changes in MSCs differentiation.

Mechanical stimulation, particularly through low-intensity vibration (LIV) platforms, has emerged as a strategy for delivering low-magnitude mechanical signals to the musculoskeletal system. **[14]** LIV is often considered a safe exercise surrogate that provides anabolic mechanical input to maintain musculoskeletal competence. Related approaches, including whole-body vibration (WBV), low-magnitude mechanical signals (LMMS), and low-magnitude, high-frequency vibration, have also been investigated for their potential skeletal and muscular benefits. Clinically, LIV delivered at frequencies ranging from approximately 10 to 100 Hz has been reported to improve bone quality and quantity in young women with osteoporosis, promote bone indices in childhood cancer survivors, and benefit individuals with conditions associated with impaired skeletal health. **[15–17]** Studies in animal models have further demonstrated that exogenously applied mechanical stimulation can increase trabecular bone density and volume, enhance bone stiffness and strength, and attenuate age- or disease-associated bone loss. **[18–20]** In addition, LIV has been reported to increase muscle strength, contractility, and cross-sectional area, supporting its broader anabolic effects on the musculoskeletal system. In stem cell studies, LIV and related low-magnitude mechanical stimuli have typically been applied across a range of frequencies, often approximately 10–800 Hz, and have been shown to modulate MSCs behavior, including reducing adipogenic differentiation while promoting osteogenic differentiation and cellular proliferation. **[21,22]** Collectively, these findings suggest that low-magnitude mechanical stimulation can influence MSCs fate and musculoskeletal tissue homeostasis. However, whether LIV can effectively modulate the differentiation and functional phenotype of MSCs during advanced chronological aging is not yet fully understood.

Despite growing evidence that LIV can modulate MSCs fate and promote osteogenic lineage commitment, whether aged MSCs retain the ability to respond to mechanical signals remains poorly understood. This question is particularly important in the context of chronological aging, during which MSCs exhibit metabolic dysfunction, increased cellular senescence, altered lineage commitment, and changes in cellular and cytoskeletal mechanics. Although LIV has been shown to regulate MSCs differentiation and musculoskeletal function, most previous studies have focused on young or relatively healthy cells and animals, leaving the effects of long-term LIV exposure on MSCs residing in the bone marrow of aged animals largely unexplored. In particular, it remains unclear whether long-term *in vivo* LIV can modify the lineage differentiation and mechanical phenotype of MSCs derived from aged animals. Furthermore, the extent to which LIV modifies the mechanical phenotype of aged MSCs, including cytoskeletal organization, cellular stiffness, and focal adhesion properties, as well as its possible effects on MSCs lineage commitment, remains unknown. Addressing these questions may provide important insights into how mechanical stimulation regulates MSCs function during *in vivo* chronological aging and whether LIV represents a potential strategy to mitigate age-associated declines in musculoskeletal regeneration.

In this study, we investigated whether LIV can modulate the functional and mechanical phenotype of MSCs during advanced chronological aging. We hypothesized that prolonged LIV exposure would alter the mechanical and cytoskeletal properties of aged MSCs and shift their lineage commitment toward osteogenesis while decreasing adipogenesis. To test this hypothesis, MSCs isolated from 24-month-old mice with or without a 19-mo *in vivo* LIV exposure, started at 5-mo of age were evaluated for differentiation, senescence as well as cellular structural and mechanical measures. In addition, transcriptomic and epigenetic analyses were performed to identify potential molecular programs associated with the LIV-induced changes. Together, these approaches allowed us to determine whether LIV improves MSCs function during advanced aging.

## 2. Materials and methods

### 2.1. Animal handling

C57BL/6J female mice were obtained from Jackson Laboratories (JAX) and housed in separately ventilated cages under controlled conditions. All animals were provided with appropriate access to food and water, and the study was conducted in compliance with institutional guidelines. All procedures were approved by the Boise State University Institutional Animal Care and Use Committee.

### 2.2. Low intensity vibration (LIV) treatment in mice

LIV treatment commenced at 5 months of age and continued until 24 months, for a total of 19 months. Mice were placed separately empty chambers on the vibrating platform and received vertical sinusoidal LIV at 0.7 g and 90 Hz for 20 min per treatment, twice a day, 5 days per week for 19 months. Non-vibrated mice served as controls (−LIV) were subjected to similar conditions but were not subjected to LIV.

### 2.3. Micro-CT analysis

Proximal tibiae (n = 7 mice per group) were stored frozen in PBS-moistened gauze until scanning and subsequently scanned while immersed in PBS using an X-ray microtomography system (SkyScan 1172F). The scanning parameters included an X-ray source voltage of 55 kV, a current of 181 μA, a voxel size of 10.33 µm³, and an integration time of 230 ms. For trabecular bone analysis, the volume of interest (VOI) began 10 slices distal to the proximal growth plate (physis) and extended 1 mm distally. The cortical bone VOI was defined beginning 2.15 mm distal to the physis and extending an additional 0.5 mm. Image reconstruction was performed using NRecon software (Bruker). Following image reorientation, when necessary, quantitative measurements were obtained using CTAn (Bruker), and three-dimensional renderings were generated with CTVol software.

### 2.4. Wheel access regimen

Following LIV treatment, mice (n = 6 per group) were randomly allocated to either a voluntary wheel-running group or a sedentary control group without wheel access, using a previously described protocol. Briefly, after a one-week acclimation period in the new cages, the animals underwent a six-week voluntary wheel access regimen. Mice were individually housed, and daily running activity, including total distance, elapsed time, average speed, and maximum speed, was recorded using a cyclocomputer (CatEye Velo 7). Body weight and food consumption were monitored and recorded weekly.

### 2.5. Bone marrow cell extraction, expansion, and culture

Bone marrow cells were harvested from the tibiae and femurs using a centrifugation-based method, as previously described by us and others. **[23,24]** Briefly, the bones were dissected, cleaned of surrounding soft tissue, and centrifuged at more than 12,000 × g to collect the bone marrow. The resulting pellet was resuspended in growth medium containing α-MEM supplemented with 20% fetal bovine serum (FBS), 100 U/mL penicillin, and 100 μg/mL streptomycin, and then plated for culture. After 48 h, the cultures were washed with phosphate-buffered saline (PBS) to remove non-adherent cells, and fresh, prewarmed growth medium was added. To eliminate remaining non-adherent and loosely attached cells, the culture surface was gently rinsed by repeatedly aspirating and dispensing the medium approximately five times, followed by the addition of 2 mL of fresh medium. The cells were subsequently cultured until approximately 50% confluence, detached, and replated to expand the cell population. Further expansion was performed until the cultures reached approximately 80% confluence before subsequent experiments.

### 2.6. Characterization of Bone Marrow-Derived Mesenchymal Stem Cells

Bone marrow-derived MSCs (n = 3 mice per group) were characterized by immunofluorescence staining for established MSC-associated surface markers and hematopoietic lineage markers. Briefly, cells were seeded onto tissue culture plates, fixed in 4% paraformaldehyde for 10 min at room temperature, and permeabilized with 0.3% Triton X-100 in PBS. Nonspecific antibody binding was minimized by blocking with 5% bovine serum albumin (BSA; Sigma-Aldrich, Cat. No. A3912-100G) in PBS. Cells were then incubated overnight at 4°C with primary antibodies against the MSC-associated positive markers CD73 (Invitrogen, Cat. No. PA5-29750), CD105 (Invitrogen, Cat. No. MA5-23894), as well as the hematopoietic lineage-associated negative markers CD34 (Invitrogen, Cat. No. 14-0341-82) and CD11b (Invitrogen, Cat. No. 14-0112-82). After primary antibody incubation, cells were treated with the corresponding fluorescent secondary antibodies. F-actin was visualized using Alexa Fluor™ 594-conjugated phalloidin (Cayman Chemical, Cat. No. 20549), while nuclei were counterstained with NucBlue™ Live Cell Stain (Invitrogen, Cat. No. R37605), following the manufacturers’ instructions. Further details of the immunofluorescence staining protocol are provided in the subsequent section.

The clonogenic potential of the isolated MSCs (n = 3 mice per group) was evaluated using a colony-forming unit-fibroblast (CFU-F) assay. Briefly, 500 cells were plated in each well of a 6-well plate and cultured for 14 days at 37°C in a humidified atmosphere containing 5% CO₂. Following incubation, colonies were fixed and stained with 0.5% crystal violet (Sigma-Aldrich) for 5 min at room temperature. Colonies containing more than 50 cells were imaged using a light microscope and subsequently quantified.

### 2.7. Immunofluorescence staining and quantification

For immunofluorescence analysis, bone marrow-derived MSCs from the +LIV and −LIV groups (n = 3 mice per group) were seeded in imaging plates at a density of 1 × 10⁴ cells per well and cultured in growth medium for 48 h. Cells were fixed with 4% paraformaldehyde for 10 min at room temperature, permeabilized with 0.3% Triton X-100 in PBS, and blocked with 5% bovine serum albumin (BSA; Sigma-Aldrich, Cat. No. A3912-100G) prepared in PBS.

The cells were incubated overnight at 4°C with primary antibodies against Ki67 (Novus Biologicals, Cat. No. NB110-89717), p16 (Cell Signaling Technology, Cat. No. 80772S), vinculin (Abcam, Cat. No. ab129002), and lamin A/C (Cell Signaling Technology, Cat. No. 4C11), each diluted 1:300 in blocking buffer. Following PBS washes, cells were incubated with Alexa Fluor™ 594 goat anti-rabbit IgG secondary antibody (Invitrogen, Cat. No. A11037) at a 1:300 dilution for 1 h at room temperature in the dark. The actin cytoskeleton was labeled with Phalloidin-iFluor™ 488 (Cayman Chemical, Cat. No. 20549), and nuclei were counterstained with NucBlue™ Live Cell Stain (Invitrogen, Cat. No. R37605), according to the manufacturers’ instructions.

Fluorescence images were captured using a Zeiss LSM 900 confocal microscope equipped with 10× and 20×/1.4 objectives. Ki67, p16, vinculin, and lamin A/C images were acquired using the 10× objective. Identical imaging parameters were maintained across experimental groups to ensure consistency during quantitative analysis.

For Ki67 and p16 quantification, fluorescence images from the corresponding channels were converted to 8-bit grayscale using ImageJ (version 1.52). A uniform intensity threshold was applied consistently across all images to define regions of interest (ROIs). Integrated fluorescence density was quantified and normalized to the corresponding cell number to account for differences in cell abundance. Vinculin and lamin A/C fluorescence signals were quantified using CellProfiler (version 4.2.8) with a custom-designed image analysis pipeline.**(25)** Briefly, nuclei and vinculin- or lamin A/C-associated signals were identified from their respective fluorescence channels, followed by cell segmentation and object identification. The MeasureObjectIntensity and MeasureObjectSizeShape modules were used to quantify fluorescence intensity and area measurements at the individual-object level. Integrated fluorescence intensity and object area were subsequently normalized to the corresponding cell number to account for differences in cell abundance. The resulting normalized measurements were used for subsequent statistical analyses.

### 2.8. Atomic force microscopy of cellular viscoelastic properties

Aged MSCs (n = 3 mice per group) were seeded at a density of 4,000 cells/cm^2^ to ensure isolated single cell engagement. Atomic force microscopy (AFM) imaging was conducted after 24 hours of culture in alpha-MEM supplemented with 20% FBS, 100 U/mL penicillin, and 100 μg/mL streptomycin. Cells were maintained at 37 °C using a stage-top Petri dish heater and supplied with 5% CO₂ via adaptor rings connected to a CO₂ controller (PeCon) throughout AFM acquisition. Each plate was imaged for no longer than 1 hour.

AFM was performed using a NanoWizard V Bioscience (Bruker, Billerica, MA, USA) integrated with a Zeiss LSM Airyscan (Zeiss, Oberkochen, Germany) for cell visualization. Mechanical measurements were acquired with MLCT-SPH-5UM probes (Bruker, Billerica, MA, USA). Probe deflection sensitivity was determined in non-contact mode using the JPK software, entering the manufacturer’s pre-calibrated spring constant of 0.044 N/m for the probe used throughout the study. Mechanical data were obtained using Force Spectroscopy and MicroRheology experiments in the JPK SPM software and. mechanical data were processed using JPK Data Processing software.

Force Spectroscopy force ramps were acquired with a fixed setpoint of 0.6 nN. All force curves were sampled at 2048 Hz. 6 × 6-pixel force maps were acquired over 100 × 100 µm regions centered on the nucleus. To obtain apparent Young’s modulus (E_app_), raw force curves were first smoothed using a Gaussian filter with a width of 3.00, then baseline-corrected over a standardized fit range of 50–100% of the x-axis (height) and further corrected for offset and tilt. Contact point correction was applied to define the contact point as the first crossing of the y-axis (vertical deflection). Vertical tip position (piezo height minus cantilever deflection) was used to account for cantilever bending. E_app_ was calculated by fitting force-indentation curves (derived from vertical deflection vs. vertical tip position) with the Hertz/Sneddon contact model assuming a spherical indenter geometry (equation #), where the contact radius is related implicitly to the indentation depth, with a nominal probe radius of 5 µm, assuming a Poisson’s ratio of 0.40. E_app_ values were restricted to a physiological range of 0–30 kPa to exclude spurious contacts with artifacts or the substrate, and outliers were removed using a 1.5× interquartile range (IQR) criterion.

For MicroRheology, 2 x 2 pixel force maps (10 µm × 10 µm) were acquired directly over the nuclear region of each cell. A total of five force maps were acquired per cell, with the modulation frequency logarithmically swept from 10 Hz to 220 Hz using three points per decade (10 Hz, 22 Hz, 46 Hz, 100 Hz, and 220 Hz) to calculate transition frequency (f_T_). MicroRheology force ramps were segmented into four phases: (1) an extension period to a setpoint of 0.6 nN, (2) a 500 ms constant-height hold, (3) a modulation phase consisting of 20 cycles with a 2 nm amplitude, and (4) a retraction period. The modulation segment of the MicroRheology plot (vertical deflection vs. time) was analyzed with a standardized fit range of 10–90%. Storage modulus (G’) and loss modulus (G”) were derived from the modulation segment using JPK Data Processing Software, also under the assumption of a spherical indenter geometry. For the dynamic fits, cells were again assumed to have a Poisson’s ratio of 0.40. Outliers were identified and excluded using a 1.5× interquartile range (IQR) criterion applied to the corresponding Dynamic Young’s modulus (E*), acquired concurrently with G′ and G″. f_T_ was determined by simple linear regressions fit to the loss tangent (tanδ) (tanδ = G”/G’) as a function of frequency for each cell. The x-intercept of the derived regression equation where y =1 (G′ = G″), was defined as f_T_. **[26]**

### 2.9. Adipogenic differentiation and lipid accumulation assays

Bone marrow-derived MSCs (n = 3 mice per group) from the +LIV and −LIV groups were plated at a density of 1.5 × 10⁵ cells per well in 6-well plates for Oil Red O staining. After 24 h of culture in growth medium, adipogenic differentiation was induced using α-MEM supplemented with 10% fetal bovine serum (FBS), 100 U/mL penicillin, 100 μg/mL streptomycin, 5 μg/mL insulin, 0.1 μM dexamethasone, and 50 μM indomethacin. The adipogenic induction medium was replaced every 2–3 days over a 7-day period.

Lipid accumulation was assessed by Oil Red O staining, as previously described. **[27]** Briefly, cells were fixed in 10% neutral-buffered formalin for 30 min at room temperature, rinsed with distilled water, and treated with 100% propylene glycol USP (Poly Scientific R&D Corporation, Cat. No. S264-8OZ) for 2 min. The cells were then incubated with Oil Red O solution (Poly Scientific R&D Corporation, Cat. No. S1848-8OZ) for 15 min, ensuring complete coverage of the cell monolayer. After removing excess stain, the cells were differentiated in 85% propylene glycol (Poly Scientific R&D Corporation, Cat. No. S264A-8OZ) for 1 min and thoroughly rinsed with distilled water before imaging.

### 2.10. Osteogenic differentiation and mineralization assays

For osteogenic differentiation, MSCs from the +LIV and −LIV groups (n = 3 mice per group) were plated at 2 × 10⁵ cells per well in 6-well plates for Alizarin Red S staining. Once the cells reached full confluence, the growth medium was replaced with osteogenic induction medium containing α-MEM supplemented with 20% fetal bovine serum (FBS), 100 U/mL penicillin, 100 μg/mL streptomycin, 10 mM β-glycerophosphate, and 50 μg/mL ascorbic acid. The induction medium was changed every 2–3 days over a 21-day period.

At the end of the differentiation period, calcium-rich mineral deposition was evaluated using Alizarin Red S (Sigma-Aldrich, Cat. No. A5533-25G), following a previously described protocol. **[28]** Briefly, cells were fixed in 70% ethanol for 1 h at room temperature and subsequently stained with 40 mM Alizarin Red S for 10 min. After staining, excess dye was removed, and the cells were rinsed five times with distilled water before imaging.

### 2.11. Quantitative real-time PCR

To assess the expression of adipogenic and osteogenic marker genes, bone marrow-derived MSCs (n = 3 mice per group) were plated in 6-well plates and maintained under adipogenic or osteogenic differentiation conditions. Total RNA was collected after 7 days of adipogenic induction and 21 days of osteogenic induction using TRIzol™ Reagent, followed by RNA purification with the QIAzol RNA extraction kit according to the manufacturers’ protocols.

Purified RNA was reverse-transcribed into complementary DNA (cDNA) using a Script cDNA Synthesis Kit in a thermal cycler. Quantitative real-time PCR (qRT-PCR) was then conducted using SsoAdvanced™ Universal SYBR® Green Supermix (Bio-Rad Laboratories, Cat. No. 1725270) on a CFX Real-Time PCR Detection System (Bio-Rad Laboratories, Hercules, CA, USA). Gene expression was normalized to 18S rRNA and relative expression levels were determined using the 2^−ΔΔCt method. Primer sequences for all genes analyzed are provided in Table 1.

**Table 1.**
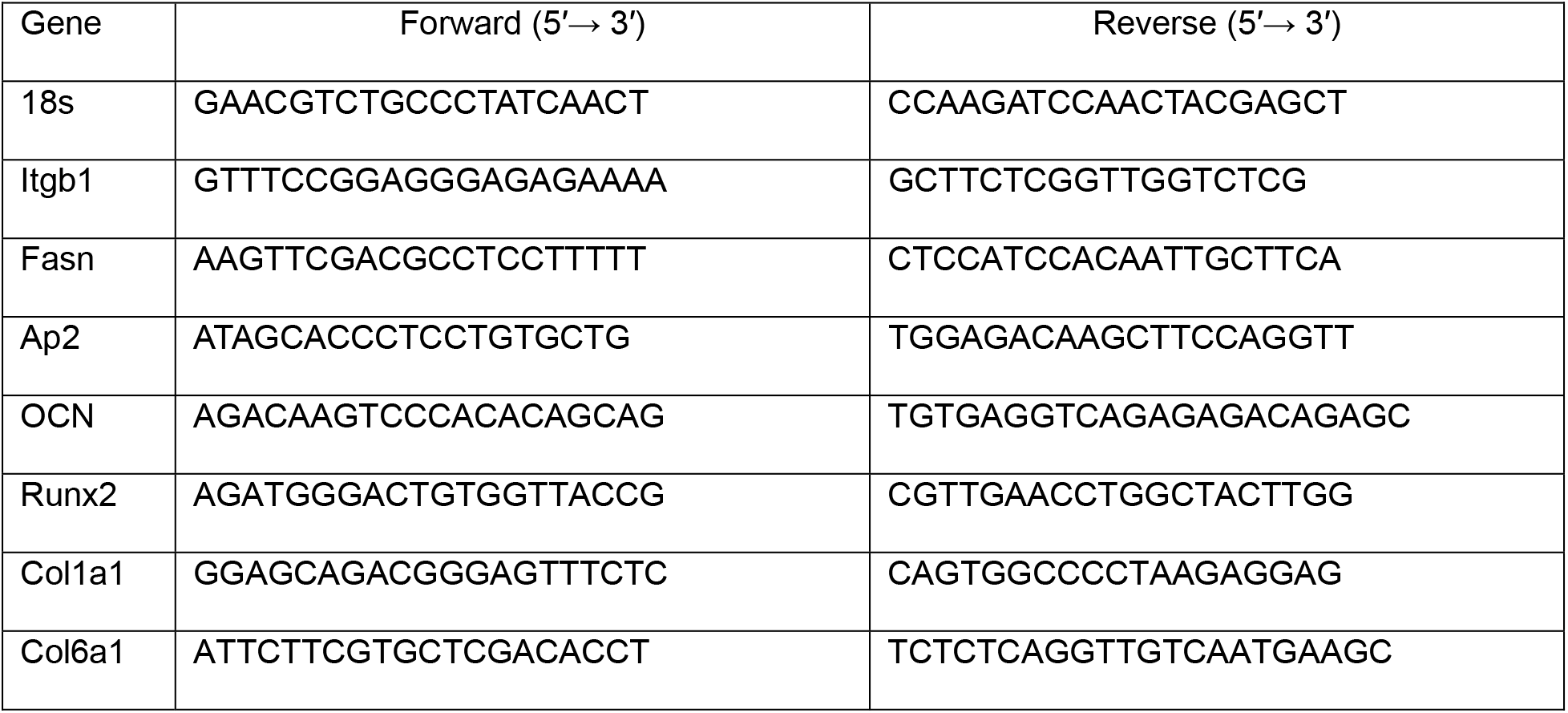
Primers used for RT-qPCR.

### 2.12. RNA sequencing and bioinformatics analysis

After adipogenesis and osteogenesis, total RNA was isolated from MSCs using the RNeasy Mini Kit (Qiagen, Hilden, Germany) following the manufacturer’s protocol. RNA sequencing was performed using three independent biological RNA samples from the +LIV group and one pooled RNA sample from the -LIV group. The pooled −LIV reference was prepared by combining equal amounts of total RNA from individual biological samples. The purified RNA samples were sent to Novogene Co., Ltd. (Sacramento, CA, USA) for library preparation, mRNA sequencing, and initial bioinformatics processing.

Briefly, a reference genome index was generated using HISAT2 (v2.0.5), and paired-end clean reads were mapped to the reference genome using the same software. Gene-level read counts were obtained with featureCounts (v1.5.0-p3), while gene expression levels were calculated as fragments per kilobase of transcript per million mapped reads (FPKM), based on transcript length and mapped read counts. Following normalization of gene expression data, expression levels in the +LIV group were compared with the pooled −LIV reference. Genes exhibiting a minimum 25% increase or decrease in expression relative to the −LIV reference (fold change ≥1.25 or ≤0.80, respectively) were selected for exploratory gene expression analysis. Functional annotation and pathway enrichment analyses were subsequently performed to identify biological processes associated with long-term LIV exposure using the Database for Annotation, Visualization and Integrated Discovery (DAVID)

### Statistical Analysis

Statistical analyses were performed using GraphPad Prism (version 10.2.0; GraphPad Software, LLC, San Diego, CA, USA). Data are presented as mean ± standard deviation (SD). Statistical significance was determined using a two-tailed unpaired Student’s *t*-test when parametric assumptions were satisfied. Normality was assessed by the Shapiro–Wilk test and homogeneity of variances was evaluated using Levene’s test. When assumptions were not met, we used the Mann–Whitney U test as a non-parametric alternative. A *p*-value of less than 0.05 was considered statistically significant.

## 3. Results

### 3.1. Isolation and identification of MSCs from experimental mice

As shown in **Figure 1a**, MSCs were isolated from mice in both the −LIV and +LIV groups at 24 months of age. The isolated cells were characterized using colony-forming unit (CFU) assays (**Figure 1b–c**) and immunofluorescence staining for MSC-associated positive and negative markers (**Figure 1d–g**). CFU assays demonstrated that MSCs from both groups retained colony-forming capacity. Immunofluorescence analysis further confirmed the expression of the positive MSCs markers CD73 and CD105, along with the absence of the negative markers CD34 and CD11b in both groups. Furthermore, MSCs from both groups demonstrated adipogenic and osteogenic differentiation potential when cultured in their respective differentiation media. Collectively, these findings support the successful isolation and characterization of bone marrow-derived MSCs from both the −LIV and +LIV groups.

**Figure 1.**
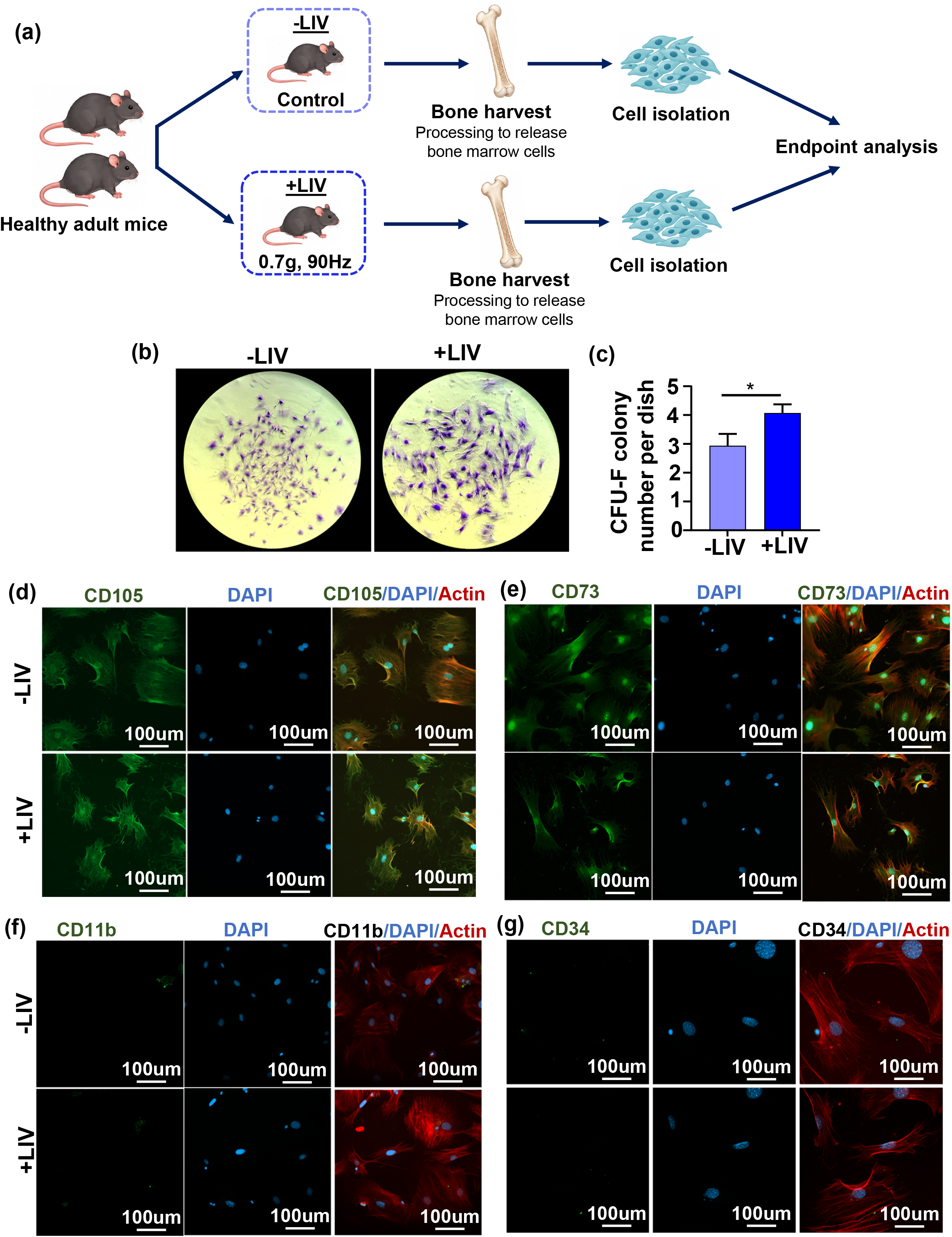
Long-term low-intensity vibration (LIV) treatment, isolation of bone marrow-derived mesenchymal stem cells (MSCs), and assessment of colony-forming capacity and immunophenotypic characteristics. (a) Schematic representation of the experimental design. Mice received long-term LIV treatment (0.7 g, 90 Hz) beginning at 5 months of age and continuing until 24 months of age. At 24 months, bone marrow-derived MSCs were isolated from LIV-treated (+LIV) and non-vibrated control (−LIV) mice. (b) Representative images of colony-forming unit-fibroblast (CFU-F) assays following 14 days of culture. (c) Quantification of CFU-F colony numbers per dish, demonstrating significantly greater colony formation in the +LIV group compared with the −LIV group (p < 0.05). (d, e) Representative immunofluorescence images showing the expression of the MSC-associated positive surface markers CD105 and CD73, respectively. (f, g) Representative immunofluorescence images showing the absence of the hematopoietic lineage-associated markers CD11b and CD34, respectively. F-actin is shown in red, nuclei are counterstained with DAPI (blue), and surface markers are shown in green. Experiments were performed using passage 2 MSCs isolated from three independent biological replicates per group (n = 3 mice per group). Representative images were acquired from multiple randomly selected fields of view for each biological replicate. Data are presented as mean ± SD. Statistical comparisons between the +LIV and −LIV groups were performed using an unpaired, two-tailed Student’s t-test. *p < 0.05, **p < 0.01, ***p < 0.001, and ****p < 0.0001.

### 3.2. LIV improves exercise performance in aged mice

**Figure 2a** showing the LIV system, including the signal generator and control unit, vibration platform, and experimental chamber used to deliver mechanical simulation. Vibration waveforms demonstrating the oscillatory motion along the three orthogonal axes (X, Y, and Z) during LIV application. The waveforms illustrate the predominantly vertical oscillatory displacement and the corresponding vibration profiles across the three axes. At 24 months of age, tibiae from one hindlimb of each mouse in the −LIV and +LIV groups were harvested for micro-computed tomography (micro-CT) analysis. Comparing the −LIV group with the +LIV group at 24 months of age showed a trend towards an increased trabecular bone volume fraction (BV/TV, +68%, p=0.090) and trabecular number (Tb.N, +73%, p=0.10) and an 8% decrease in total cross-sectional area (Tt. Ar., p=0.051) suggesting a potential beneficial effect of LIV on bone microarchitecture in aged mice (**Table 2**). Furthermore, voluntary wheel-running behavior was monitored over six weeks in mice with or without access to a running wheel (**Figure 2b**). Body mass and food intake were also measured throughout the experimental period (**Figure 2c–d**). The results showed that the average daily running distance significantly increased from 0.87 ± 0.22 km/day in the −LIV group to 1.47 ± 0.21 km/day in the +LIV group (**Figure 2f**, p < 0.01). Similarly, the average running velocity increased significantly from 0.38 ± 0.03 km/h to 0.53 ± 0.05 km/h (**Figure 2g**, p < 0.01). However, no significant difference was observed in average running time between the groups **(Figure 2e**). Collectively, these findings suggest that LIV may improve voluntary running performance in aged mice, as evidenced by increased running distance and velocity without a significant change in running time.

**Figure 2.**
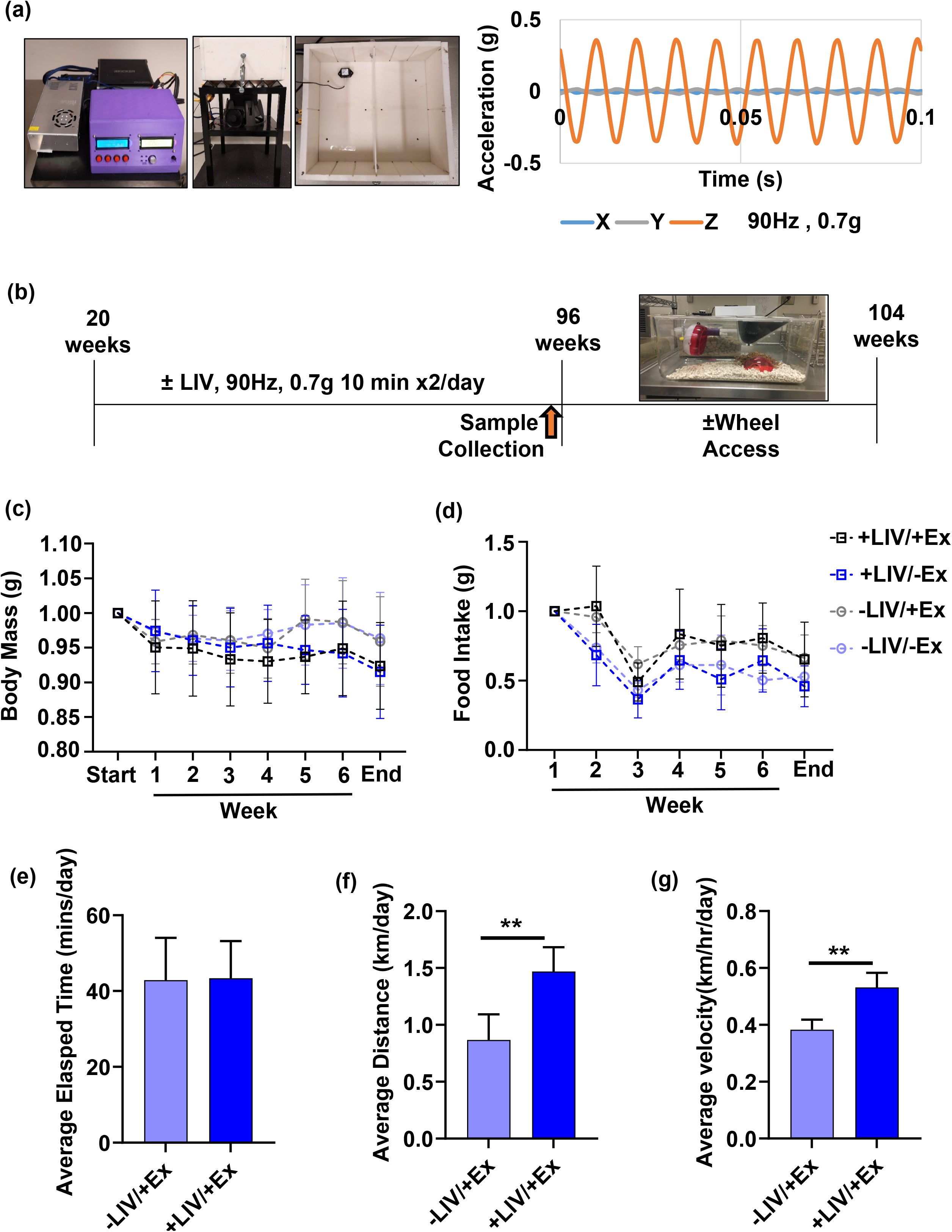
Long-term LIV treatment improves exercise performance in aged mice. (a) Experimental setup of the LIV system, and representative vibration waveforms along the X,Y, and Z axes.(b) Schematic representation of the experimental timeline. Following 96 weeks of LIV treatment, mice underwent a 1-week recovery period, followed by a 6-week assessment of exercise performance using a running wheel equipped with a cyclometer. (c) Body mass and (d) food intake were monitored throughout the 6-week observation period. (e) Average running time (elapsed time) was not significantly different between groups, whereas (f) average running distance and (g) average running velocity were significantly increased in the +LIV group compared with the −LIV group (p < 0.01). Data were obtained from 6 mice per group. Values are presented as mean ± SD. Statistical comparisons between the +LIV and −LIV groups were performed using an unpaired, two-tailed Student’s t-test. *p < 0.05, **p < 0.01, ***p < 0.001, and ****p < 0.0001.

**Table 2.** Quantitative analysis of tibial bone microarchitecture and cortical bone morphology by micro-CT.

| Parameter | Description | -LIV<br>(mean±SD) | +LIV<br>(mean±SD) | p-value<br>(two-tailed) | p-value<br>(one-tailed) |
|---|---|---|---|---|---|
| BV/TV | Bone volume fraction (%) | 3.971±2.013 | 6.650±3.561 | 0.090 | 0.045 |
| Tb.Th. | Trabecular thickness (mm) | 0.058±0.011 | 0.060±0.005 | 0.715 | 0.357 |
| Tb.Sp. | Trabecular spacing (mm) | 0.617±0.084 | 0.546±0.113 | 0.189 | 0.099 |
| Tb.N. | Trabecular number (mm<sup>-1</sup>) | 0.663±0.37 | 1.144±0.695 | 0.102 | 0.051 |
| Tt. Ar. | Total cross-section (mm<sup>2</sup>) | 1.520±0.080 | 1.386±0.157 | 0.053 | 0.026 |
| Ct. Ar. | Cortical area(mm2) | 0.603±0.041 | 0.559±0.119 | 0.343 | 0.171 |
| Ct. Ar./Tt. Ar. | Cortical area fraction (%) | 0.397±0.022 | 0.403±0.072 | 0.839 | 0.419 |
| Ct. Th. | Cortical thickness (mm) | 0.140±0.008 | 0.139±0.015 | 0.343 | 0.171 |

### 3.3. LIV enhances Ki67 expression and reduces the senescence-associated marker p16 in aged MSCs

We assessed the expression of the proliferation-associated marker Ki67 and the senescence-associated marker p16 in MSCs. As shown in **Figures 3a and 3c**, immunofluorescence staining (red fluorescence) revealed the abundance of Ki67 and p16, respectively, in MSCs isolated from the −LIV and +LIV groups. Ki67 fluorescence intensity per cell was significantly higher in the +LIV group (0.132 ± 0.11) than in the −LIV group (0.037 ± 0.006; **Figure 3b**, p < 0.01). Conversely, p16 fluorescence intensity per cell was significantly reduced following LIV treatment, decreasing from 0.742 ± 0.08 in the −LIV group to 0.34 ± 0.005 in the +LIV group (**Figure 3d**, p < 0.01).

**Figure 3.**
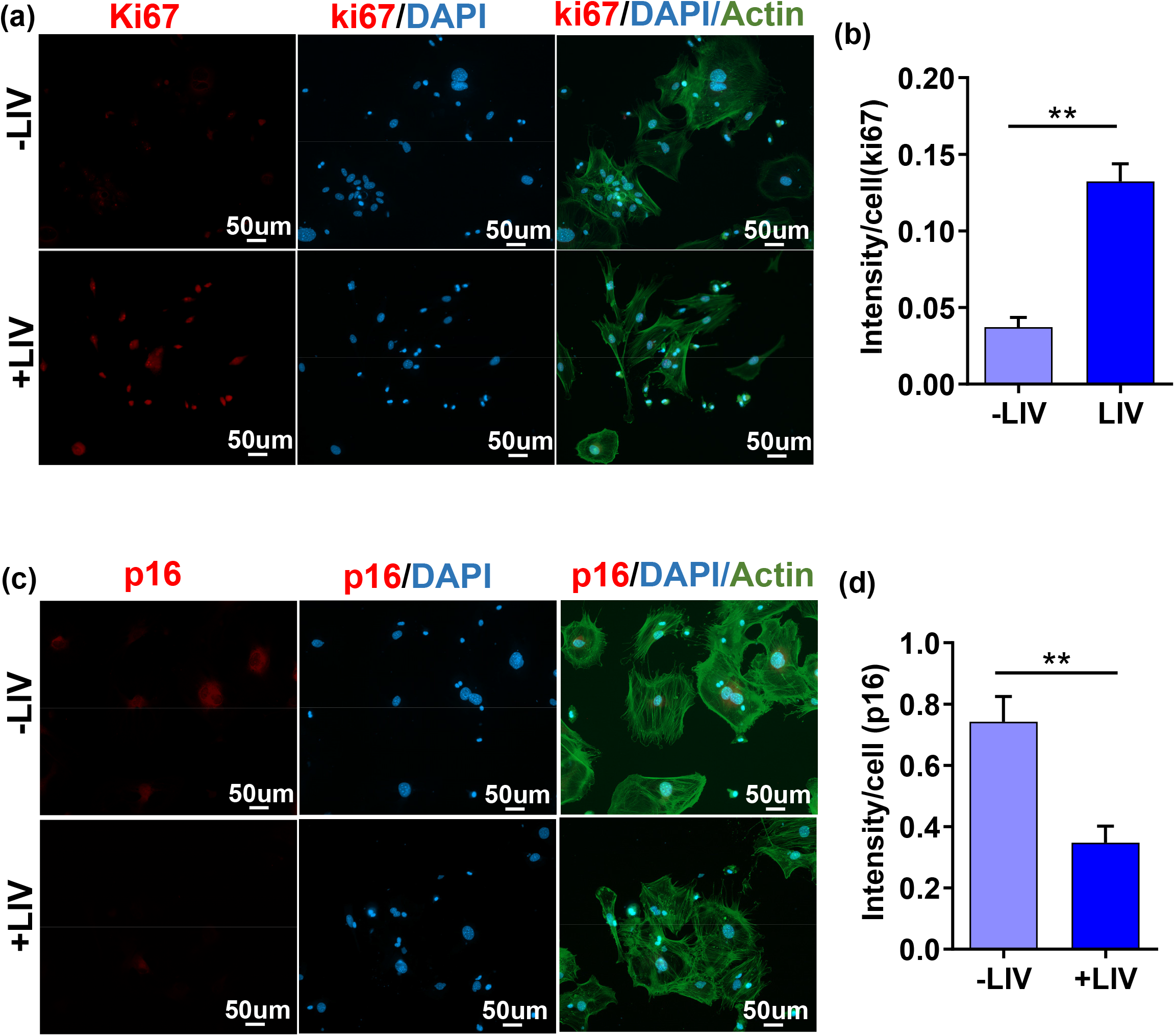
Long-term LIV treatment increases Ki67 expression and reduces p16 expression in bone marrow-derived MSCs from aged mice. Representative immunofluorescence images showing the expression of (a) Ki67, a marker of cellular proliferation, and (c) p16, a marker of cellular senescence, in bone marrow-derived MSCs isolated from +LIV and −LIV groups. Quantitative analyses of (b) Ki67 and (d) p16 fluorescence intensity demonstrate significantly increased Ki67 expression and decreased p16 expression in MSCs from the +LIV group compared with the −LIV group (both p < 0.001). Images were acquired from multiple randomly selected fields of view for each biological replicate. Fluorescence intensity was quantified and normalized to cell number. All experiments were performed using passage 1 MSCs from three independent biological replicates per group (n = 3 mice per group). Values are presented as mean ± SD. Statistical comparisons between the +LIV and −LIV groups were performed using an unpaired, two-tailed Student’s t-test.. *p < 0.05, **p < 0.01, ***p < 0.001, and ****p < 0.0001.

### 3.4. LIV attenuates adipogenic differentiation in aged MSCs

To assess the practical implications of LIV in aged MSCs, we first measured adipogenic differentiation potential following *in vivo* LIV. As shown in **Figure 4a**, Oil Red O staining revealed lipid droplet formation in MSCs from both the −LIV and +LIV groups following adipogenic induction. Quantitative analysis demonstrated a significant reduction in lipid droplet area fraction, from 7.59 ± 1.17% in the −LIV group to 3.76 ± 1.63% in the +LIV group (**Figure 4b**, p < 0.001). To further investigate the LIV-associated transcriptional changes in adipogenesis, we surveyed the RNA-sequencing (RNA-seq) data. Gene Ontology (GO) enrichment analysis using the Database for Annotation, Visualization and Integrated Discovery (DAVID) was performed to identify biological processes associated with the observed changes in adipogenic function. The pathway analysis revealed downregulation of biological processes associated with adipogenesis in the +LIV group (**Figure 4c**). In contrast, LIV-associated changes in extracellular matrix and cell–matrix interaction pathways suggested potential alterations in the cellular microenvironment that may contribute to reduced adipogenic differentiation (**Figure 4d**).

**Figure 4.**
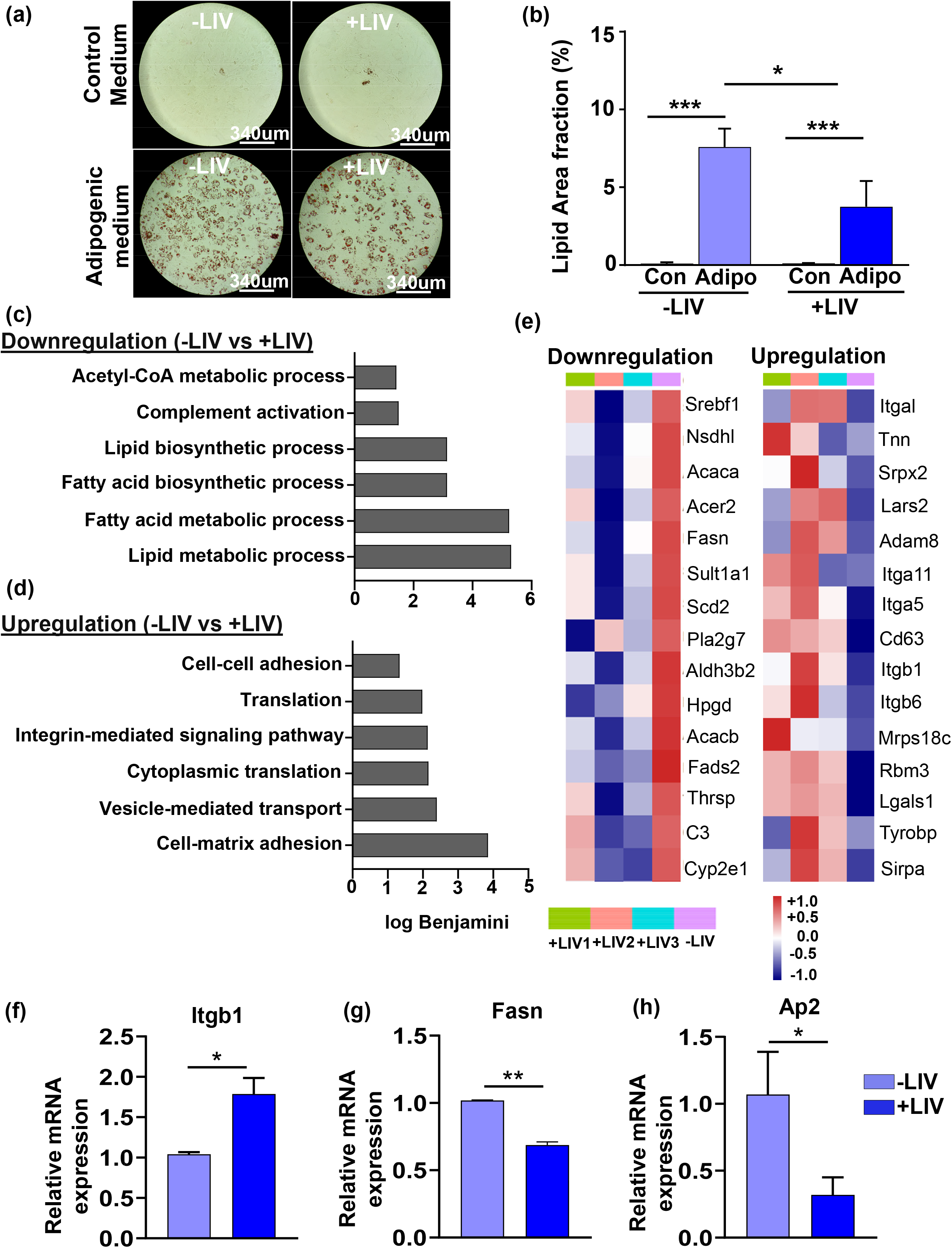
Long-term LIV treatment attenuates adipogenic differentiation and modulates adipogenesis-associated gene expression in aged MSCs. (a) Representative images of Oil Red O staining showing adipogenic differentiation and intracellular lipid accumulation in bone marrow-derived MSCs isolated from +LIV and −LIV groups under normal growth and adipogenic induction conditions. (b) Quantitative analysis of adipogenic differentiation based on the percentage of Oil Red O-positive area, demonstrating significantly reduced lipid accumulation in the +LIV group compared with the −LIV group (p < 0.05). DAVID functional enrichment analysis showing significantly enriched Gene Ontology (GO) terms associated with (c) downregulated and (d) upregulated differentially expressed genes (DEGs) in the +LIV group relative to the −LIV group. The x-axis represents the −log10 of the Benjamini-adjusted p-value, and the y-axis lists the corresponding enriched GO categories. (e) Heatmaps showing the expression patterns of selected adipogenesis-associated genes, highlighting differentially expressed genes in the +LIV group compared with the −LIV group. Based on the RNA-sequencing results and adipogenic differentiation findings, further performed quantitative real-time PCR (qPCR). Relative mRNA expression levels of (f) Itgb6, (g) Fasn, and (h) AP2 demonstrate significant changes in adipogenesis-associated gene expression following long-term LIV treatment. These expression changes are consistent with the reduced adipogenic potential observed in MSCs from the +LIV group. All experiments were performed using passage 2 MSCs derived from three independent biological replicates per group (n = 3 mice per group). Values are presented as mean ± SD. Statistical comparisons between the +LIV and −LIV groups were performed using an unpaired, two-tailed Student’s t-test. *p < 0.05, **p < 0.01, ***p < 0.001, and ****p < 0.0001.

Based on the RNA-seq findings (**Figure 4e**), we selected Fasn (fatty acid synthase) and Itgb1 (integrin β1) for further validation via RT-qPCR analysis. Fasn, which encodes fatty acid synthase, plays an important role in fatty acid biosynthesis and lipid accumulation during adipocyte differentiation. Consistent with the RNA-seq results, Fasn expression was significantly reduced in the +LIV group compared with the −LIV group (**Figure 4g**, p <0.01). Conversely, Itgb1, which encodes the integrin β1 subunit and is involved in cell–matrix interactions and signaling, was significantly upregulated following LIV treatment (**Figure 4f**, p <0.05). In addition, expression of Ap2 (fatty acid-binding protein 4), a marker associated with adipocyte differentiation and lipid metabolism, was significantly decreased in the +LIV group (**Figure 4h**, p < 0.001).

Collectively, these findings demonstrate that *in vivo* LIV treatment attenuates the adipogenic differentiation potential of aged MSCs, as evidenced by reduced lipid accumulation, downregulation of adipogenesis-associated biological pathways, and changes in the expression of genes involved in lipid biosynthesis.

### 3.5. LIV potentiates osteogenic differentiation in aged MSCs

We next evaluated MSCs osteogenic differentiation potential following *in vivo* LIV treatment. As shown in **Figure 5a**, Alizarin Red staining revealed mineralized nodule formation in MSCs from both the −LIV and +LIV groups following osteogenic induction. Mineralized nodule area was significantly greater in the +LIV group (0.66 ± 0.09 µm²) than in the −LIV group (0.19 ± 0.08 µm^²;^ **Figure 5b**, p < 0.01).

**Figure 5.**
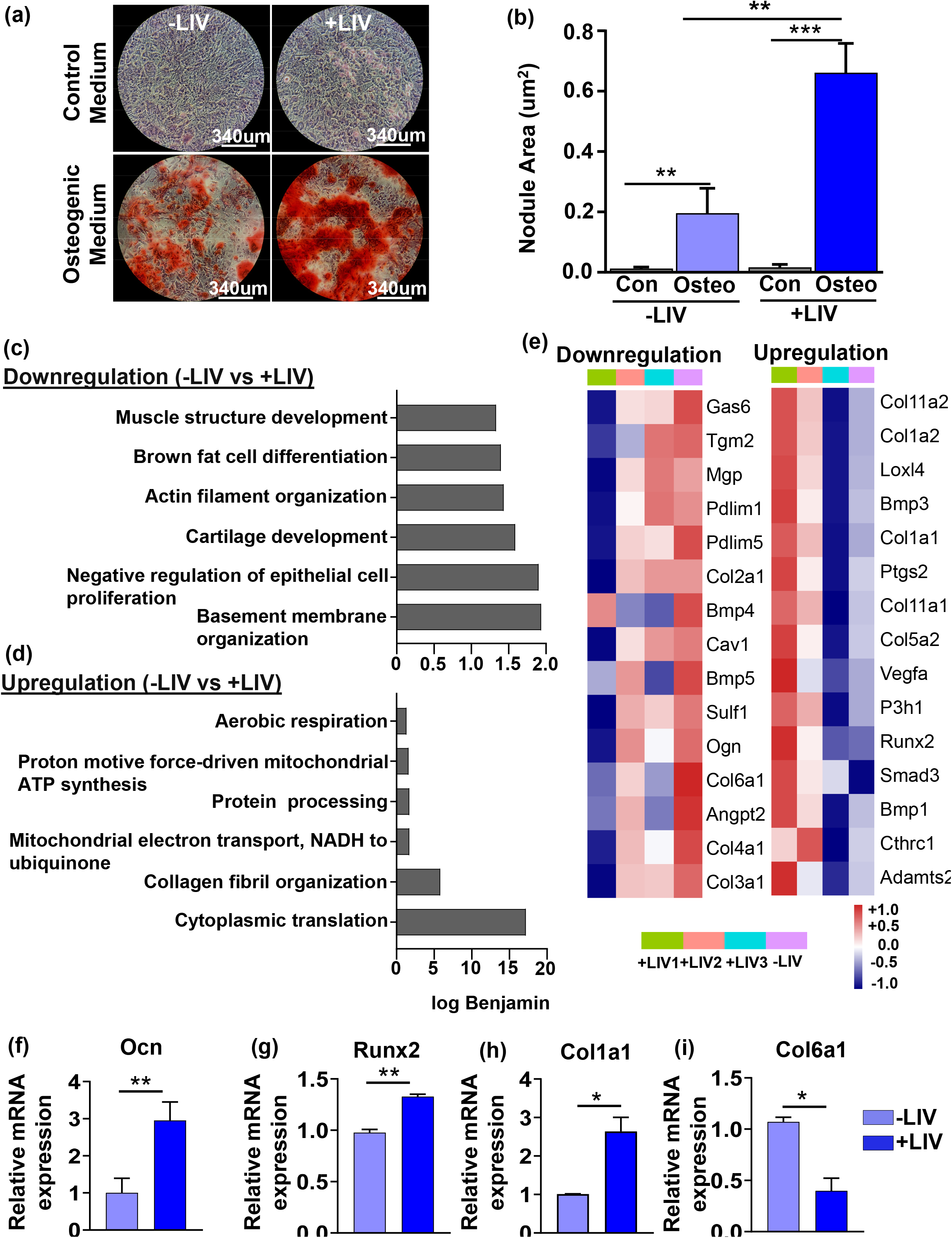
Long-term LIV treatment enhances osteogenic differentiation and modulates osteogenesis-associated gene expression in aged MSCs. (a) Representative images of Alizarin Red S staining showing osteogenic differentiation and mineralized nodule formation in bone marrow-derived MSCs isolated from +LIV and −LIV groups under normal growth and osteogenic induction conditions. (b) Quantitative analysis of osteogenic differentiation based on the percentage of Alizarin Red S-positive area, demonstrating significantly increased mineralized nodule formation in the +LIV group compared with the −LIV group (p < 0.01). DAVID functional enrichment analysis showing significantly enriched Gene Ontology (GO) terms associated with (c) downregulated and (d) upregulated differentially expressed genes (DEGs) in the +LIV group relative to the −LIV group. The x-axis represents the −log10 of the Benjamini-adjusted p-value, and the y-axis lists the corresponding enriched GO categories. (e) Heatmaps showing the expression patterns of selected osteogenesis-associated genes, highlighting differentially expressed genes in the +LIV group compared with the −LIV group. Based on the RNA-sequencing results and osteogenic differentiation findings, selected genes were further examined by quantitative real-time PCR (qPCR). Relative mRNA expression levels of (f) Ocn, (g) Runx2, (h) Col1a1, and (i) Col6a1 demonstrate significant changes following long-term LIV treatment, with increased expression of osteogenic markers (Ocn, Runx2, and Col1a1) and decreased expression of Col6a1. These expression changes are consistent with the enhanced osteogenic differentiation observed in MSCs from the +LIV group. All experiments were performed using passage 2 MSCs derived from three independent biological replicates per group (n = 3 mice per group). Values are presented as mean ± SD. Statistical comparisons between the +LIV and −LIV groups were performed using an unpaired, two-tailed Student’s t-test. *p < 0.05, **p < 0.01, ***p < 0.001, and ****p < 0.0001.

We examined the RNA-seq data to investigate gene expression changes under osteogenic differentiation conditions in MSCs isolated from −LIV and +LIV mice. The heatmap in **Figure 5e** illustrates representative genes downregulated and upregulated in response to LIV. Furthermore, DAVID analysis identified biological pathways associated with the LIV-induced changes in osteogenic function (**Figure 5c–d)**. The analysis revealed changes in the expression of genes associated with extracellular matrix remodeling, mitochondrial and growth factor-related signaling. Notably, LIV was associated with the upregulation of genes involved in osteogenic differentiation and extracellular matrix formation, including Runx2, Col1a1, and Col1a2.

RT-qPCR confirmed increased expression of the osteogenic markers Ocn (osteocalcin, p < 0.01), Runx2 (runt-related transcription factor 2, p < 0.01), and Col1a1 (collagen type I alpha 1, p < 0.05) in the +LIV group compared with the −LIV group. (**Figure 5f-h**) In contrast, Col6a1 (collagen type VI alpha 1) expression was decreased following LIV treatment. These transcriptional changes, together with the increased mineralized nodule area, suggest that LIV treatment is associated with enhanced osteogenic differentiation in aged MSCs (**Figure 5i**, p < 0.05).

### 3.6. LIV remodels F-actin and focal adhesion architecture, increasing cell stiffness in aged MSCs

To investigate the effects of LIV on cytoskeletal organization and focal adhesions, we surveyed the F-actin architecture via phalloidin staining and focal adhesions via probing vinculin in MSCs from the −LIV and +LIV groups. As shown in **Figure 6a**, immunofluorescence staining revealed differences in F-actin organization between the two groups. F-actin fiber number was significantly higher in +LIV MSCs (1278.74 ± 101.43 per cell) than in −LIV MSCs (995.02 ± 80.91 per cell; **Figure 6b**, p < 0.05). No changes in F-actin fiber length were found. Furthermore, vinculin immunofluorescence (**Figure 6d**) revealed significant increases in both the number of vinculin-associated structures and mean fluorescence intensity in +LIV MSCs. Vinculin number increased from 421.17 ± 32.98 per cell in the −LIV group to 578.58 ± 41.67 per cell in the +LIV group (**Figure 6e**, p < 0.01), while mean fluorescence intensity increased from 0.014 ± 0.001 to 0.017 ± 0.0005 (**Figure 6f**, p < 0.05).

**Figure 6.**
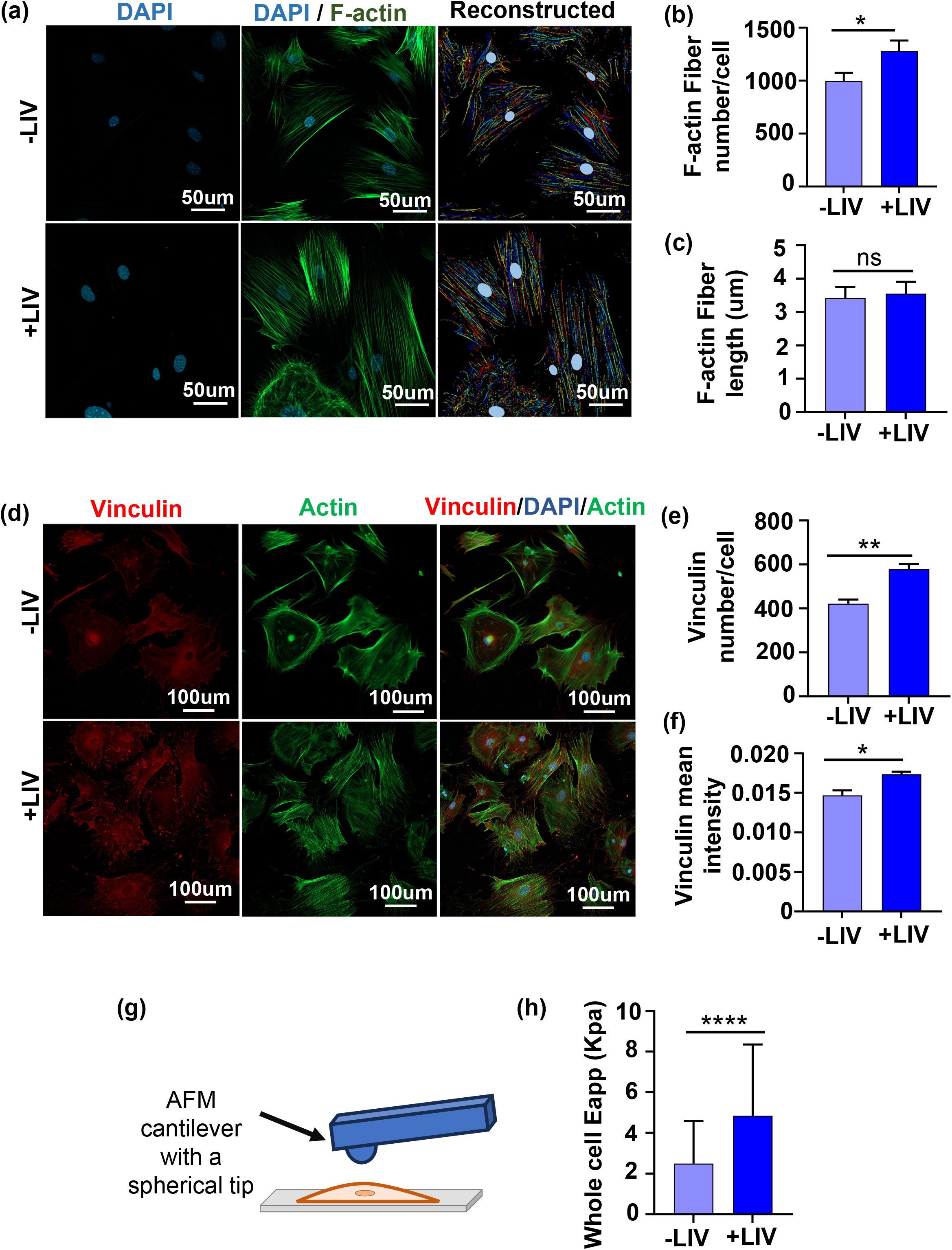
Long-term LIV treatment enhances cytoskeletal organization, focal adhesion-associated protein expression, and cellular stiffness in bone marrow-derived MSCs. (a) Representative fluorescence microscopy images and corresponding reconstructed images illustrating F-actin cytoskeletal organization (green) and nuclear staining (blue) in MSCs isolated from the −LIV and +LIV groups. (b) Quantification of F-actin fiber number per cell, showing a significant increase in the +LIV group compared with the −LIV group (p < 0.05). (c) Quantification of average F-actin fiber length, showing no significant difference between groups. (d) Representative immunofluorescence images showing vinculin localization (red), F-actin organization (green), and nuclei (blue) in MSCs isolated from the −LIV and +LIV groups. (e, f) Quantification of vinculin expression, demonstrating significant increases in (e) vinculin-positive focal adhesion number per cell and (f) mean fluorescence intensity in the +LIV group compared with the −LIV group. (g) Schematic representation of the atomic force microscopy (AFM) approach used to assess cellular mechanical properties. (h) Quantification of the apparent Young’s modulus (E_app_), demonstrating significantly increased whole-cell stiffness in the +LIV group compared with the −LIV group (p < 0.0001). All experiments were performed using passage 2 MSCs derived from three independent biological replicates per group (n = 3 mice per group). Data are presented as mean ± SD. Statistical comparisons between the +LIV and −LIV groups were performed using an unpaired, two-tailed Student’s t-test. *p < 0.05, **p < 0.01, ***p < 0.001, and ****p < 0.0001.

Given the important roles of the F-actin cytoskeleton and focal adhesions in regulating cellular structure, we further assessed whole-cell stiffness using atomic force microscopy (AFM) in live cells. As illustrated in **Figure 6g**, AFM measurements were performed by indenting individual cells with an AFM probe to characterize their apparent Young’s modulus (Eapp). Whole-cell Eapp was significantly higher in +LIV MSCs (4.843 ± 3.51 kPa) than in −LIV MSCs (2.596 ± 2.09 kPa; **Figure 6h**, p <0.0001). Collectively, these findings suggest that LIV treatment is associated with cytoskeletal remodeling, an increased number of vinculin-associated focal adhesions, and elevated apparent whole-cell stiffness in aged MSCs.

### 3.7. LIV enhances lamin A/C expression and alters the dynamic frequency response of the nucleus and epigenetic marks in aged MSCs

To investigate the effects of LIV on nuclear remodeling and mechanics, we first assessed lamin A/C expression and dynamic nuclear properties using AFM-based frequency sweep measurements. **Figure 7a** shows representative immunofluorescence images of lamin A/C in the −LIV and +LIV groups. Quantitative analysis revealed a significant 126.3% increase in lamin A/C fluorescence intensity, from 114.4 ± 23.17 in the −LIV group to 258.9 ± 17.18 in the +LIV group (**Figure 7b**, p < 0.001).

**Figure 7.**
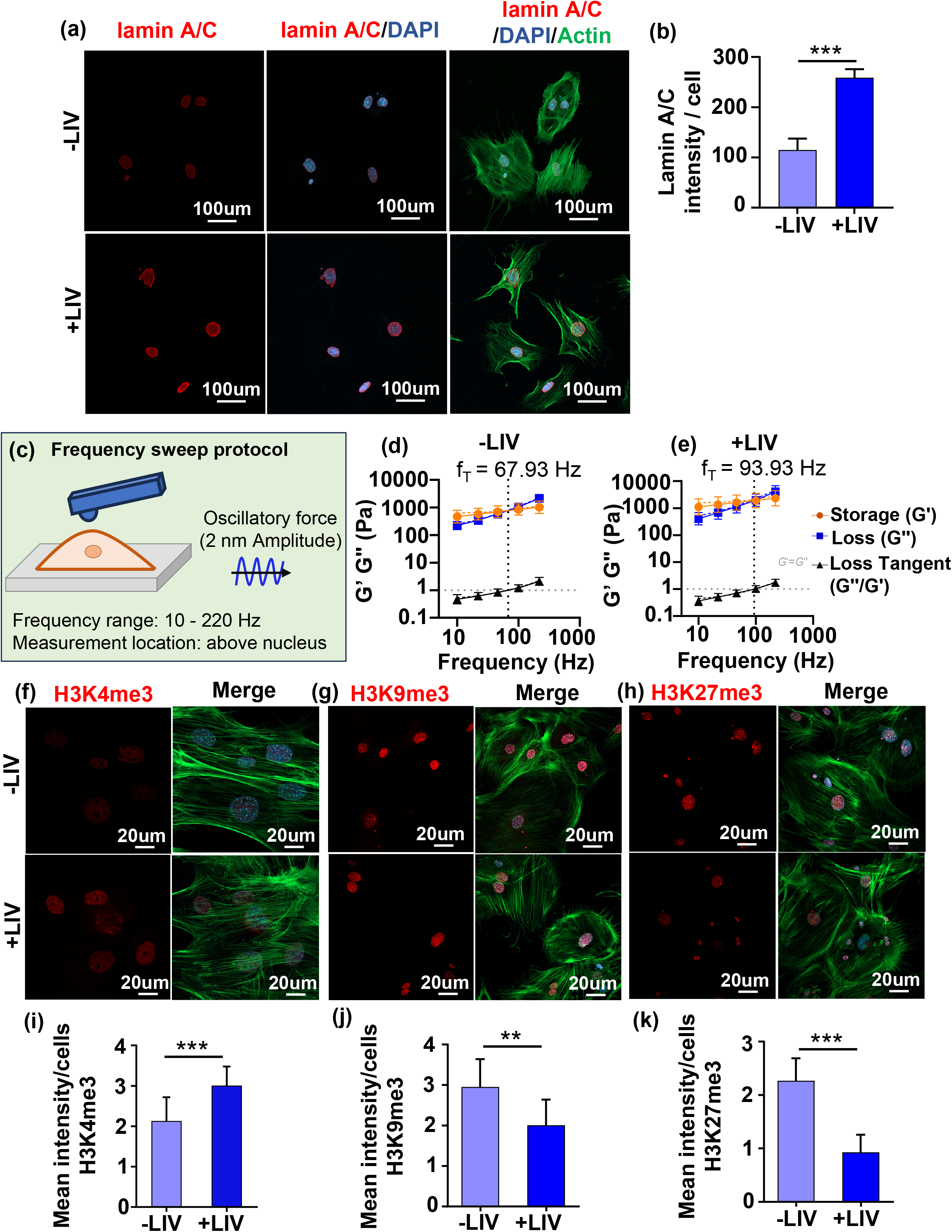
Long-term LIV treatment enhances nuclear lamin A/C expression, modulates nuclear viscoelastic properties, and alters global histone modifications in aged bone marrow-derived MSCs. (a) Representative fluorescence microscopy images showing lamin A/C expression (red), F-actin organization (green), and nuclear staining (blue) in MSCs isolated from the −LIV and +LIV groups. (b) Quantification of lamin A/C integrated fluorescence intensity, showing a significant increase in the +LIV group compared with the −LIV group (p < 0.01). (c) Schematic representation of the atomic force microscopy (AFM)-based approach used to characterize nuclear mechanical properties under oscillatory loading at different frequencies. Representative nuclear-positioned microrheology measurements of the −LIV (d,e) and +LIV (f) groups, showing linear regressions of storage (G′) and loss (G″) moduli and loss tangent (tanδ, G”/G’) across oscillatory loading frequencies (10–220 Hz). The transition frequency (fT), defined by the crossover of storage and loss moduli (G′ = G″) and corresponding to the intersection of the loss tangent regression with y = 1, was increased in the +LIV group compared with the −LIV group, indicating a shift toward elastic-dominant behavior at higher frequencies. (f–h) Representative immunofluorescence images showing global histone modifications, including H3K4me3 (f), H3K9me3 (g), and H3K27me3 (h), in MSCs from the −LIV and +LIV groups. (i–k) Quantification of mean fluorescence intensity demonstrating significantly increased H3K4me3 (i) and significantly decreased H3K9me3 (j) and H3K27me3 (k) in the +LIV group compared with the −LIV group. All experiments were performed using passage 2 MSCs derived from three independent biological replicates per group (n = 3 mice per group).. Data are presented as mean ± SD. Statistical comparisons between the +LIV and −LIV groups were performed using an unpaired, two-tailed Student’s t-test. *p < 0.05, **p < 0.01, ***p < 0.001, and ****p < 0.0001.

To further characterize the effects of LIV on nuclear mechanical properties, we performed AFM-based frequency sweep measurements at the nuclear position of aged MSCs over a frequency range of 10–220 Hz (**Figure 7c**). The crossover frequency (f_T_), defined as the frequency at which G′ = G″ and tanδ = **1**, shifted from 67.93 Hz in the −LIV group to 93.81 Hz in the +LIV group (**Figure 7d–e**). This 38% increase in crossover frequency indicates that elastic-dominant behavior persisted to a higher frequency in +LIV MSCs when measured over the nuclear region. Collectively, these findings suggest that LIV modifies lamin A/C expression and alters the dynamic mechanical behavior of the nucleus in aged MSCs.

To investigate possible histone modifications following LIV treatment in aged mice, we examined three histone methylation marks: H3K4me3, H3K9me3, and H3K27me3. Representative immunofluorescence images are shown in **Figure 7f–h**. Quantitative analysis revealed a significant increase in H3K4me3 fluorescence intensity following LIV treatment, from 2.12 ± 0.59 in the −LIV group to 3.007 ± 0.47 in the +LIV group (**Figure 7i**, p < 0.001). In contrast, H3K9me3 intensity decreased from 2.95 ± 0.68 in the −LIV group to 2.00 ± 0.63 in the +LIV group (**Figure 7j**, p < 0.001,). Similarly, H3K27me3 intensity significantly decreased from 2.26 ± 0.42 in the −LIV group to 0.92 ± 0.33 in the +LIV group (**Figure 7k**, p < 0.001). Together, these results indicate that LIV is associated with changes in global histone methylation patterns in aged MSCs, including increased H3K4me3 and decreased H3K9me3 and H3K27me3.

## 4. Discussion

Our findings indicate that 19 months of LIV treatment, from 5 to 24 months of age, improved average voluntary wheel-running distance and velocity in female mice. While these values did not fully return to those observed at 5 months of age, **[25]** LIV-treated mice showed significantly improved physical activity levels and a trend toward increased bone volume compared with age-matched controls. Once MSCs are isolated from animals with or without LIV, LIV treated MSCs show a positive bias toward osteogenesis and away from adipogesis, indicating at least a partial reversal of age associated adipogenic drift **[29]** suggesting LIV may have the potential to improve age related outcomes. In this study we only used female mice, and conclusions cannot be generalized to male mice.

Our findings further demonstrate that LIV treatment is associated with decreased expression of the senescence-associated marker p16 and increased expression of the proliferation-associated marker Ki67. Increased expression of p16 has been observed in aged MSCs, linking cellular senescence to persistent activation of the DNA damage response. **[30,31]** Moreover, the activation of DNA damage response pathways in cells from older donors has been associated with decreased expression of the proliferation marker Ki67.**[32]** Growing evidence indicates that cellular senescence contributes to the decline in MSCs functional capacity and impaired differentiation potential during aging. Consistent with the functional importance of senescent cells in aging, Farr et al. used a transgenic mouse model to selectively eliminate p16-positive senescent cells, resulting in increased bone mass, bone formation, and bone strength. **[33]** Furthermore, Wen et al. demonstrated that low-magnitude vibration could restore impaired bone formation in aged rats, potentially through the inhibition of senescence in osteogenic cells. **[34]** Their findings indicated that vibration treatment reduced the senescent cell burden and attenuated senescence through the Sirt1/p53/p21 signaling axis. Other studies have also suggested that mechanical loading can influence the cellular response to DNA damage and nuclear stress. Taken together, our findings suggest that long-term LIV treatment may modulate senescence-associated changes in MSCs during chronological aging. The increased expression of Ki67 and decreased expression of p16 indicate that LIV may help maintain the proliferative state and functional potential of aged MSCs. These changes may contribute to the improved osteogenic differentiation observed following LIV treatment. However, further studies are needed to determine whether LIV directly regulates cellular senescence and whether these changes causally contribute to the preservation of MSCs differentiation capacity.

The differentiation capacity of MSCs changes with age, often favoring adipogenesis over osteogenesis, which may contribute to reduced bone formation and the development of osteoporosis. **[35,36]** Age-related changes in MSCs include the downregulation of osteogenic genes, such as Runx2, collagen-related genes, and osteocalcin, alongside the upregulation of adipogenic markers, including Adipoq, Pparγ, and Fasn. In addition to these molecular changes, previous studies have reported increased adipocyte size and decreased mineralized matrix formation in late-passage MSCs compared with early-passage MSCs. **[37]** Similarly, studies of bone marrow-derived MSCs have demonstrated age-associated shifts in lineage commitment, characterized by reduced osteoblastogenesis and increased adipogenesis both *in vivo* and *in vitro*. **[38,39]** In the present study, we investigated whether LIV treatment could modulate the lineage commitment of MSCs during chronological aging. Our findings suggest that LIV may help preserve the osteogenic potential of aged MSCs while suppressing their adipogenic phenotype. To further explore the molecular pathways potentially associated with these changes, we used RNA sequencing to characterize gene expression profiles associated with LIV-mediated lineage commitment. Expression of genes associated with adipogenesis and lipid formation was lower in the +LIV group than in the −LIV reference, suggesting a potential shift in transcriptional programs related to adipogenic differentiation.

During osteogenesis, RNA-seq analysis revealed that LIV was associated with the upregulation of genes involved in osteogenic differentiation and extracellular matrix (ECM) formation, including Runx2, Col1a1, and Col1a2. **[40]** Additionally, the upregulation of genes associated with collagen processing and maturation, including Loxl4, Bmp1, Adamts2, and P3h1, suggests enhanced collagen biosynthesis and ECM remodeling. Consistent with these findings, qPCR results showed that LIV significantly increased the expression of Runx2, a key transcription factor that regulates osteogenic differentiation and the expression of osteogenesis-related genes. **[41]**

The cytoskeleton and focal adhesions play essential roles in mechanotransduction by transmitting mechanical signals that regulate osteogenic differentiation. Actin filaments form the structural framework of the cytoskeleton, maintaining cell shape, generating intracellular tension, and facilitating the transmission of mechanical forces throughout the cell. Vinculin, a key focal adhesion protein, strengthens the connection between the actin cytoskeleton and integrin-associated complexes, facilitating the transmission of mechanical signals from the extracellular matrix (ECM) into the cell. **[42]** Previous studies have demonstrated that actin and vinculin collectively contribute to focal adhesion organization and influence mechanosensitive signaling pathways, including YAP/TAZ and RhoA–ROCK, which can regulate osteogenic gene expression. **[43,44]** In the present study, LIV treatment remodeled actin fiber organization and vinculin distribution, with significant increases in their respective numbers in MSCs from aged mice compared with the -LIV group. These changes may facilitate the observed changes in MSCs lineage commitment. **[10,45]** Previous studies have also shown that mechanical stress influences G-protein signaling, which regulates cytoskeletal organization and can facilitate ERK translocation to the nucleus, thereby activating transcription factors and altering cellular processes, including proliferation, differentiation, and gene expression. **[46]** Regarding focal adhesions, Xiong et al. demonstrated that LIV increased vinculin levels in MSCs in response to increasing vibration intensity. **[47]** Similarly, other studies have reported that mechanical strain increases vinculin expression, which may subsequently promote osteogenic differentiation in MSCs. **[48,49]**

The cytoskeleton and focal adhesions also function as a coordinated mechanical system that contributes to cellular stiffness. In the present study, we assessed cellular stiffness following LIV treatment in aged MSCs using atomic force microscopy (AFM), alongside changes in actin fiber organization and vinculin distribution. Our results demonstrated that LIV treatment increased cellular stiffness, which may be associated with cytoskeletal remodeling and focal adhesion reinforcement. This interpretation is supported by previous studies showing that increased actin filament organization can enhance cellular stiffness, whereas actin disruption reduces cellular stiffness. **[50–52**] Yi et al. further demonstrated that LIV increased the stiffness of breast cancer cells in association with changes in actin organization. **[53]** Another study reported a 24% increase in stiffness following vibration stimulation, with this mechanical alteration associated with signaling pathways linked to osteogenic potential. **[54]** Collectively, aged MSCs isolated from LIV-treated female mice showed altered cytoskeletal organization and focal adhesion architecture, along with increased cellular stiffness, compared with MSCs from age-matched −LIV mice. Further mechanistic studies are needed to establish whether these changes directly mediate the osteogenic or adipogenic phenotypes observed in these cells.

Changes in cytoskeletal organization are also linked to nuclear structure. In our study, +LIV MSCs showed increased lamin A/C and H3K4me3 levels, together with decreased H3K9me3 and H3K27me3 levels. The nuclear lamina and epigenetic regulation represent interconnected components of cellular mechanoadaptation, potentially linking mechanical signals to changes in gene expression. **[55,56]** Further, LIV expands the frequency range of elastic-dominant mechanical response in aged MSCs, suggesting that +LIV treated MSCs may be responsive to a broader frequency range of mechanical signals before viscoelastic loss dominates their structural behavior. While the physiological benefits of this frequency shift and increased lamin A/C levels remain unclear, these changes are consistent with altered nuclear mechanical adaptation and may influence how aged MSCs transmit and dissipate mechanical forces within the bone marrow niche.

Histone modifications have been increasingly recognized as important regulators of MSCs fate determination and lineage differentiation. H3K4me3 is generally associated with transcriptionally active promoters and gene expression. **[57–59]** Several previous studies have also implicated H3K4me3 in osteogenic differentiation. For example, reduced H3K4me3 has been shown to impair osteogenesis, whereas increased H3K4me3 enrichment at the Runx2 promoter has been associated with increased Runx2 expression and enhanced osteogenic differentiation. **[60–62]** In contrast, H3K9me3 and H3K27me3 are generally associated with transcriptional repression and play important roles in regulating MSCs lineage commitment. Kang et al. demonstrated that demethylation of H3K9 at the Runx2 promoter promotes osteogenesis in bone marrow-derived MSCs. **[63]** Furthermore, increased H3K9 methylation has been observed in aged and osteoporotic mice and has been implicated in the reduced osteogenic potential of MSCs. Consistent with these findings, our RNA-seq data revealed that LIV increased the expression of osteogenic genes while reducing the expression of adipogenic genes compared with the non-LIV control group. Collectively, these findings suggest that LIV-associated changes in histone modifications may contribute to the transcriptional shift favoring osteogenesis over adipogenesis in aged MSCs. However, because histone modifications were assessed globally, their genomic distribution and specific relationships with lineage-associated genes remain unknown. Further studies are needed to determine whether the observed increases in lamin A/C and changes in mechanical behavior directly contribute to LIV-induced epigenetic remodeling or represent parallel adaptations to mechanical stimulation.

In conclusion, our findings demonstrate that *in vivo* LIV promotes osteogenic differentiation while suppressing adipogenesis in aged MSCs. Accompanying changes in cellular mechanics, histone modifications, and gene expression suggest that LIV may improve age-associated MSCs dysfunction through coordinated epigenetic and mechanical adaptations. The trend toward increased bone volume and the improvement in exercise performance highlight the potential physiological relevance of LIV in maintaining skeletal health and physical function during aging. Although the causal mechanisms linking these cellular adaptations to organism-level outcomes remain to be established, this study provides evidence supporting LIV as a potential nonpharmacological approach to modulating age-associated changes in MSCs function and promoting skeletal health.

## Author contributions

Omar Faruq: concept/design, data analysis/interpretation, manuscript writing. Calvin L Jones: concept/design, data analysis/interpretation, manuscript writing. Nilson R.S Silva: concept/design, data analysis/interpretation. Scott Birks: concept/design, data analysis/interpretation. Sean Howard: concept/design, data analysis/interpretation. Gunes Uzer: concept/design, data analysis/interpretation, financial support, manuscript writing, final approval of manuscript.

## Data accessibility statement

The RNA-sequencing data are being deposited in the NCBI Gene Expression Omnibus (GEO) database, and the accession number will be provided upon acceptance of the manuscript. The data supporting the findings of this study are available in the supplementary Materials. Other data that support the findings of this study are available on request from the corresponding author.

---

**Table S1.**
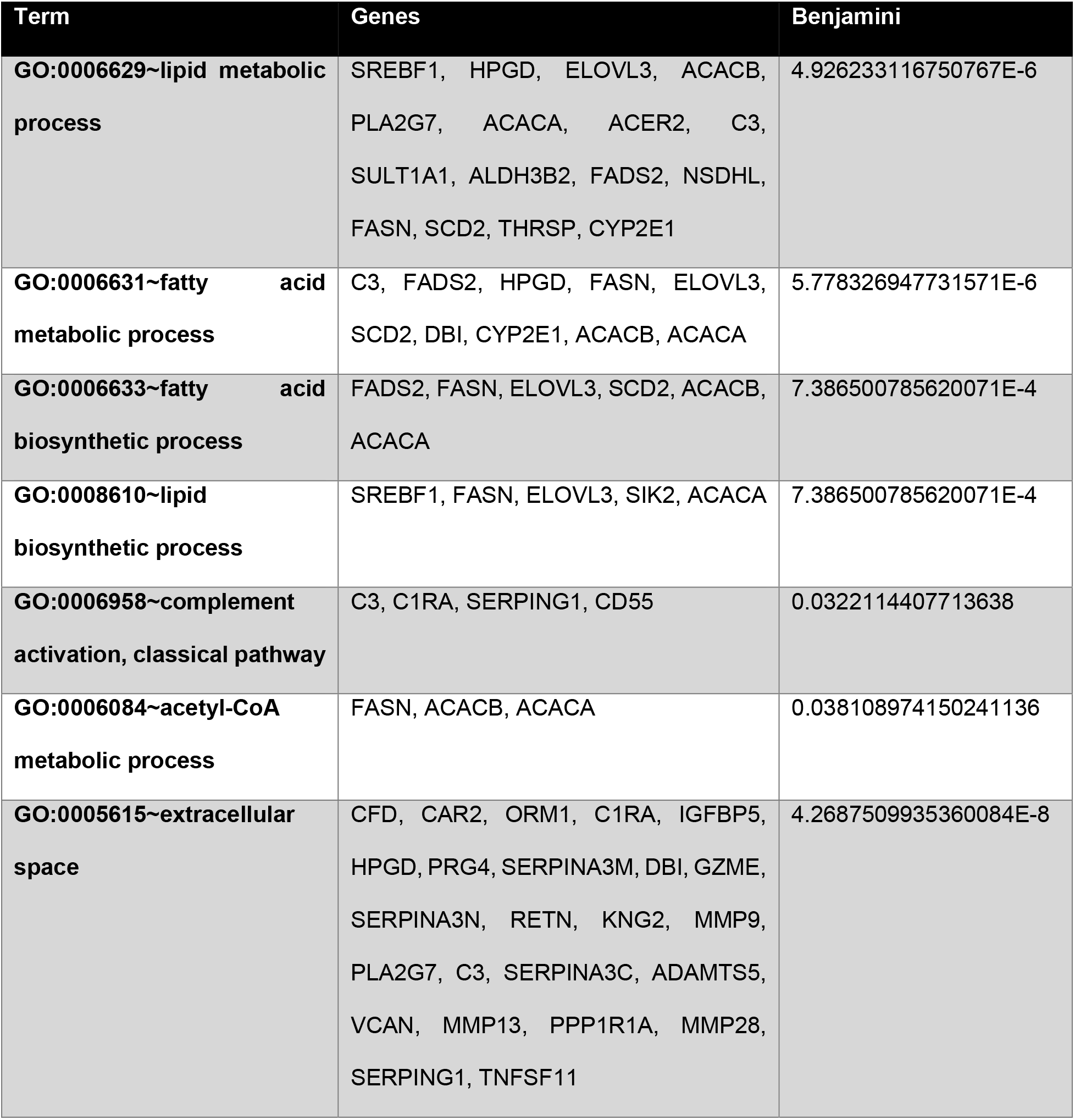
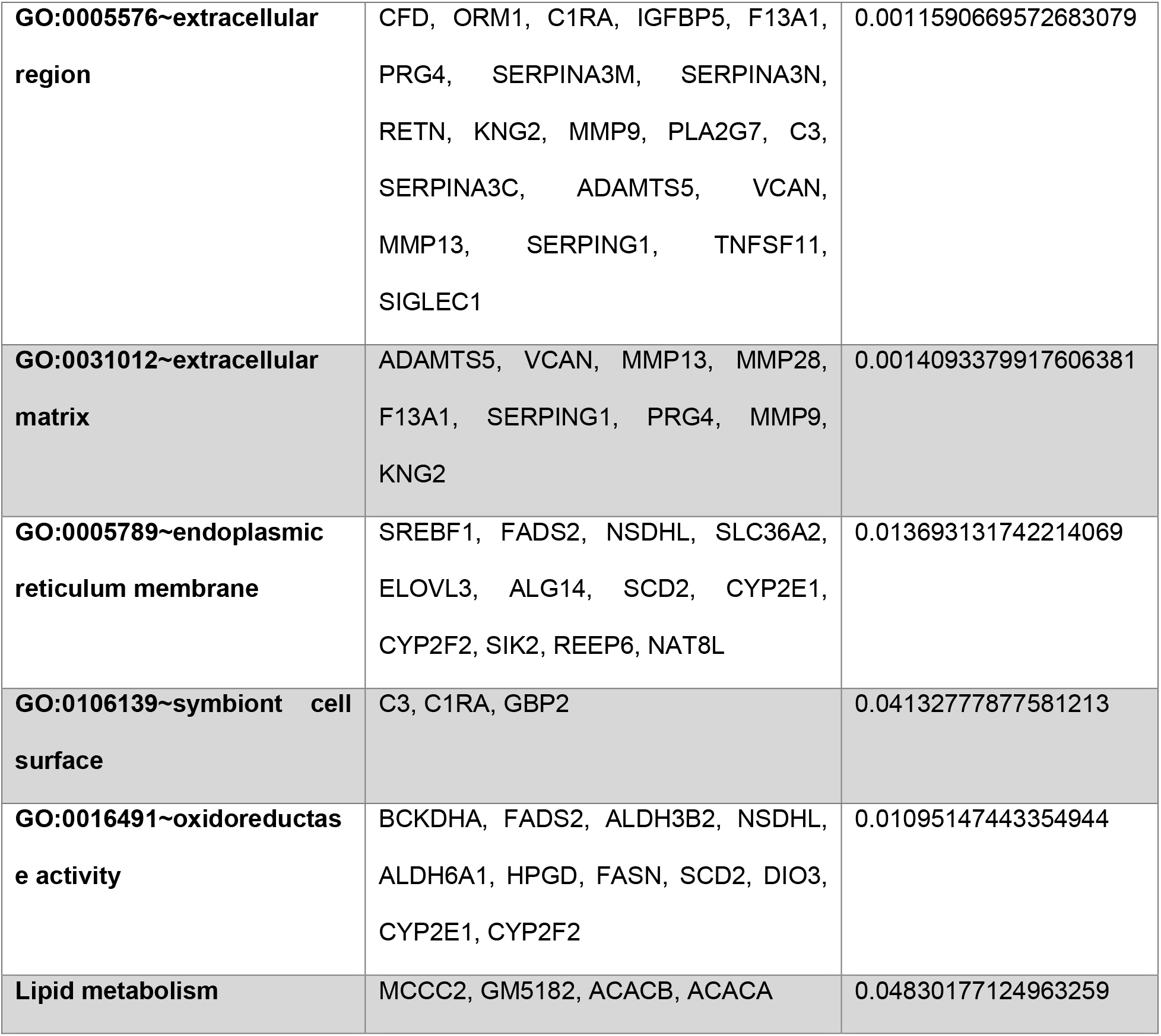
DAVID Gene Ontology (GO) enrichment analysis of genes downregulated in adipogenic medium following LIV treatment compared with untreated controls. Genes included in the analysis exhibited at least a 25% decrease in expression in the +LIV group relative to the −LIV group.

| **Term** | **Genes** | **Benjamini** |
|---|---|---|
| **GO:0006629~lipid metabolic process** | SREBF1, HPGD, ELOVL3, ACACB, PLA2G7, ACACA, ACER2, C3, SULT1A1, ALDH3B2, FADS2, NSDHL, FASN, SCD2, THRSP, CYP2E1 | 4.92623116750767E-6 |
| **GO:0006631~fatty acid metabolic process** | C3, FADS2, HPGD, FASN, ELOVL3, SCD2, DBI, CYP2E1, ACACB, ACACA | 5.77832694773157​1E-6 |
| **GO:0006633~fatty acid biosynthetic process** | FADS2, FASN, ELOVL3, SCD2, ACACB, ACACA | 7.38650078562007​1E-4 |
| **GO:0008610~lipid biosynthetic process** | SREBF1, FASN, ELOVL3, SIK2, ACACA | 7.38650078562007​1E-4 |
| **GO:0006958~complement activation, classical pathway** | C3, C1RA, SERPING1, CD55 | 0.0322114407713638 |
| **GO:0006084~acetyl-CoA metabolic process** | FASN, ACACB, ACACA | 0.0381089741502411​36 |
| **GO:0005615~extracellular space** | CFD, CAR2, ORM1, C1RA, IGFBP5, HPGD, PRG4, SERPINA3M, DBI, GZME, SERPINA3N, RETN, KNG2, MMP9, PLA2G7, C3, SERPINA3C, ADAMTS5, VCAN, MMP13, PPP1R1A, MMP28, SERPING1, TNFSF11 | 4.26875099353600​84E-8 |
| **GO:0005576~extracellular region** | CFD, ORM1, C1RA, IGFBP5, F13A1, PRG4, SERPINA3M, SERPINA3N, RETN, KNG2, MMP9, PLA2G7, C3, SERPINA3C, ADAMTS5, VCAN, MMP13, SERPING1, TNFSF11, SIGLEC1 | 0.00115906695726​83079 |
| **GO:0031012~extracellular matrix** | ADAMTS5, VCAN, MMP13, MMP28, F13A1, SERPING1, PRG4, MMP9, KNG2 | 0.00140933799176​06381 |
| **GO:0005789~endoplasmic reticulum membrane** | SREBF1, FADS2, NSDHL, SLC36A2, ELOVL3, ALG14, SCD2, CYP2E1, CYP2F2, SIK2, REEP6, NAT8L | 0.0136931317422​14069 |
| **GO:0106139~symbiont cell surface** | C3, C1RA, GBP2 | 0.0413277787758​1213 |
| **GO:0016491~oxidoreductase activity** | BCKDHA, FADS2, ALDH3B2, NSDHL, ALDH6A1, HPGD, FASN, SCD2, DIO3, CYP2E1, CYP2F2 | 0.0109514744335​4944 |
| **Lipid metabolism** | MCCC2, GM5182, ACACB, ACACA | 0.0483017712496​3259 |

**Table S2.**
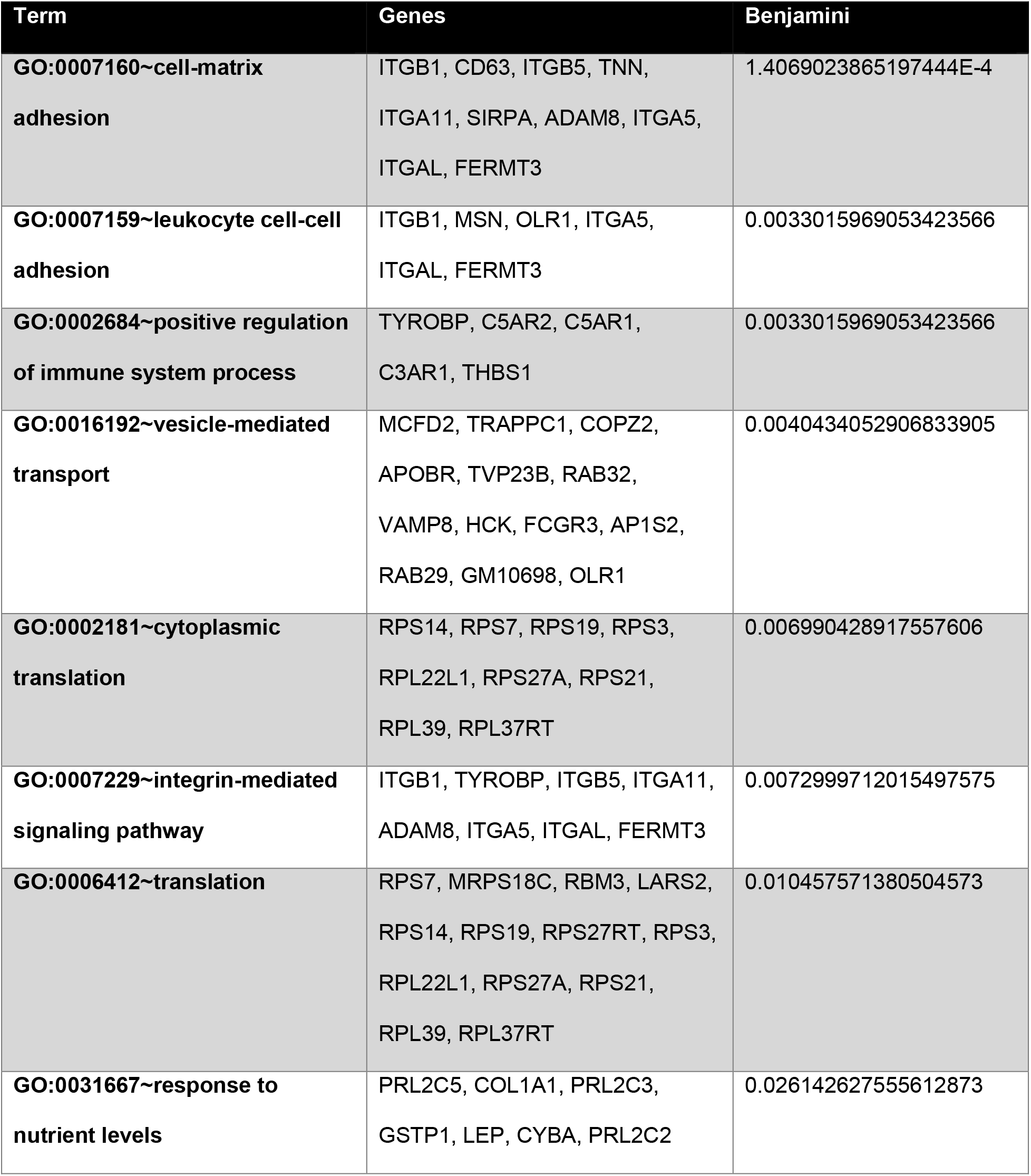
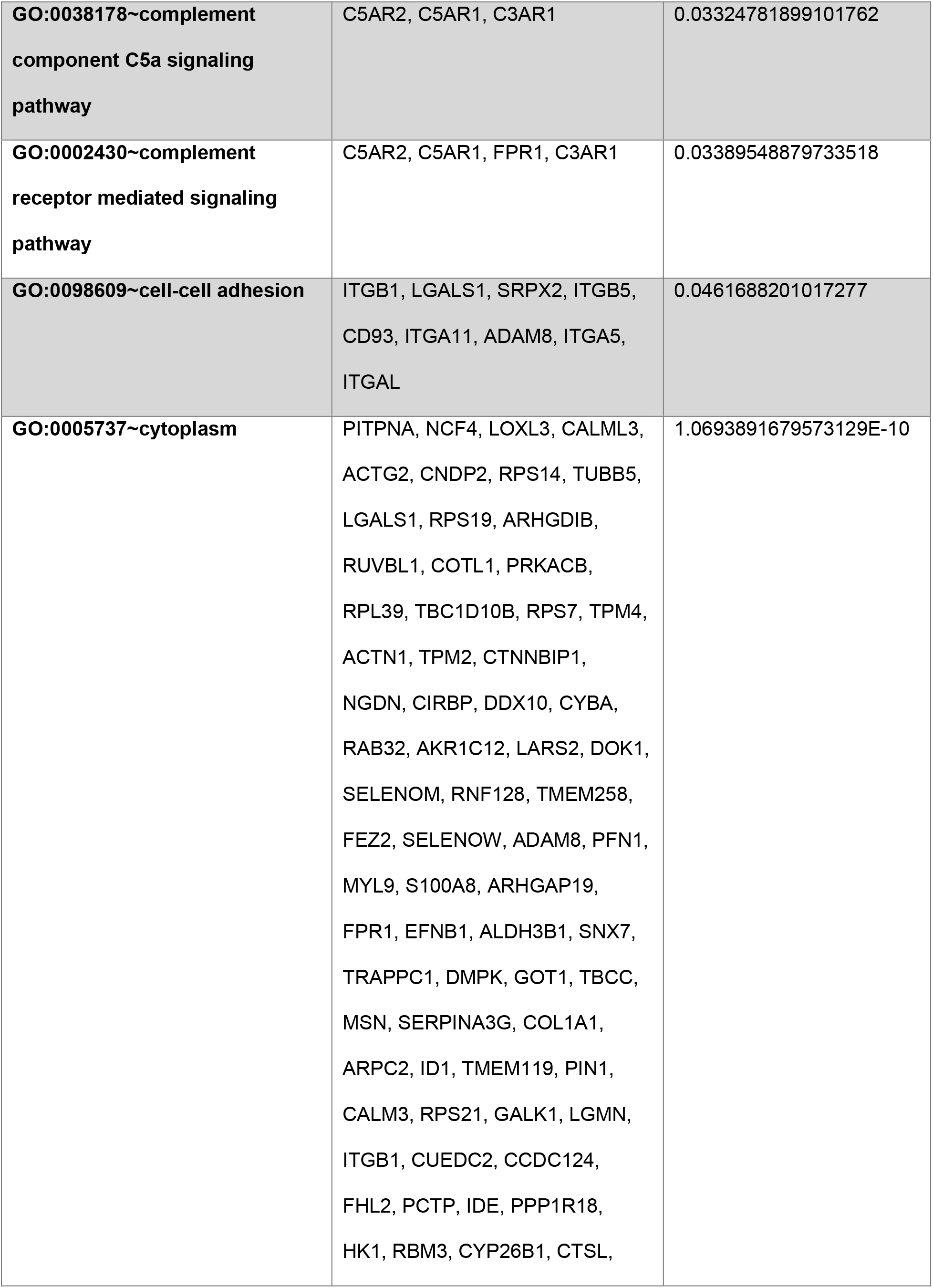
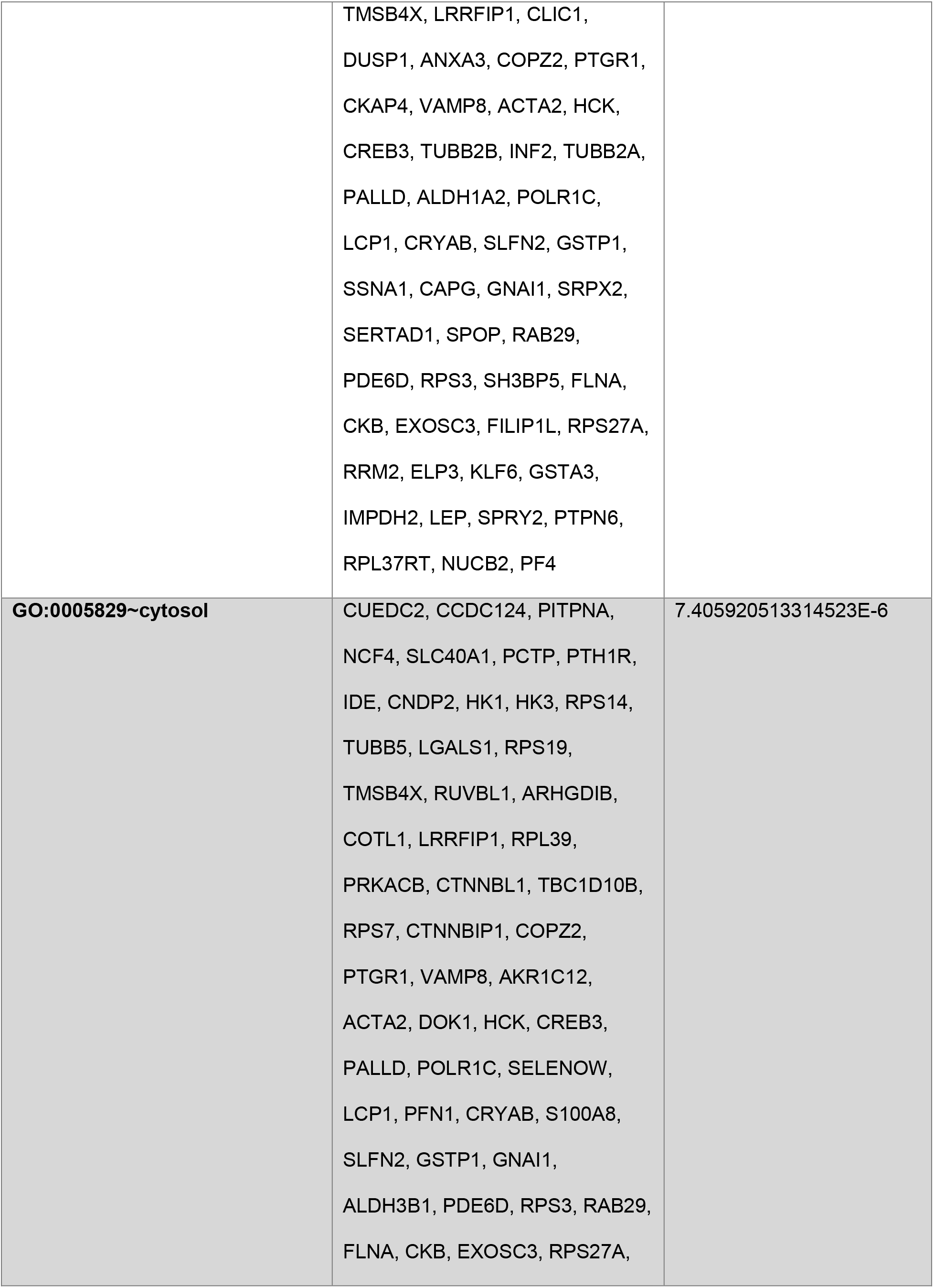
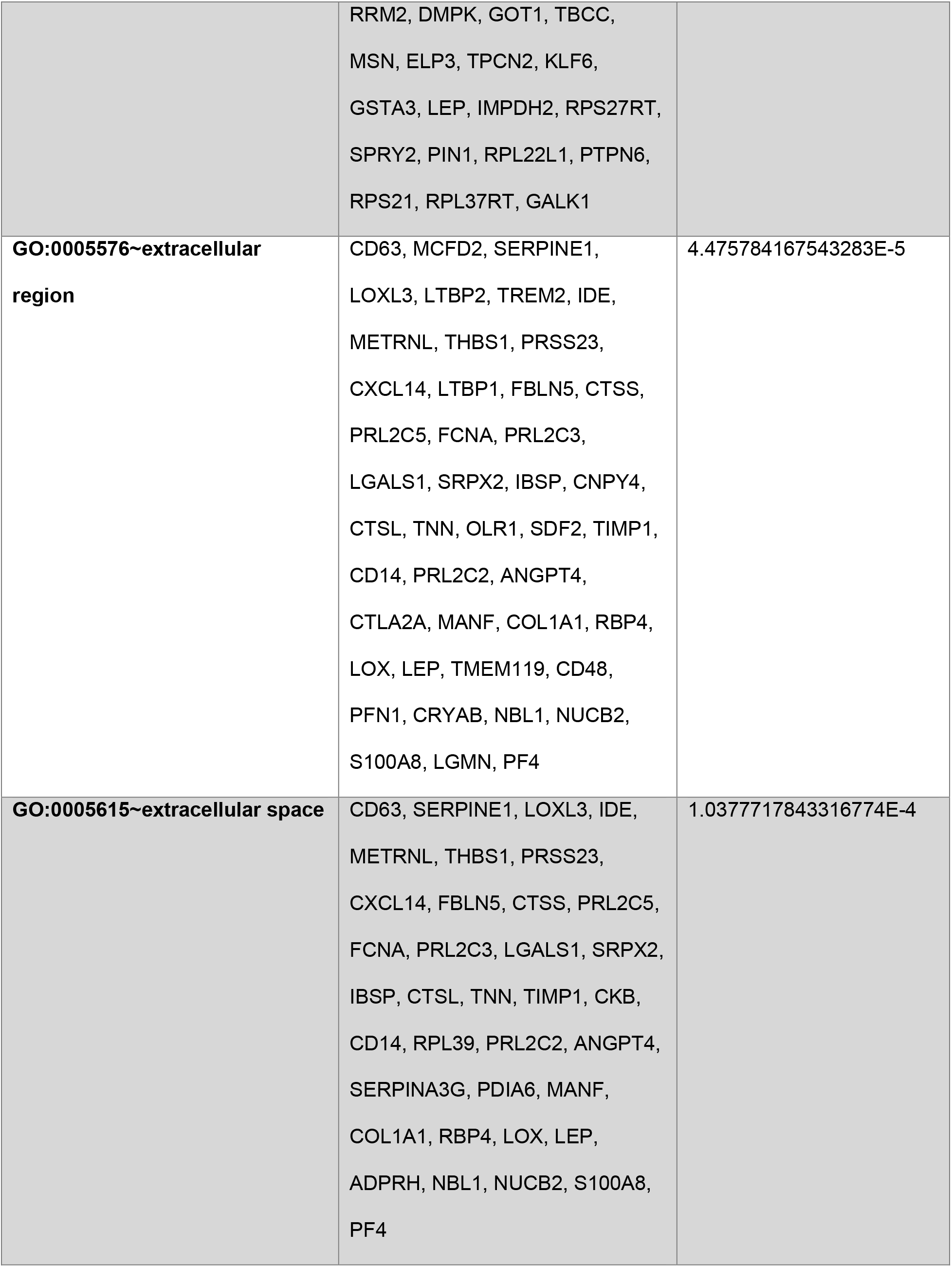
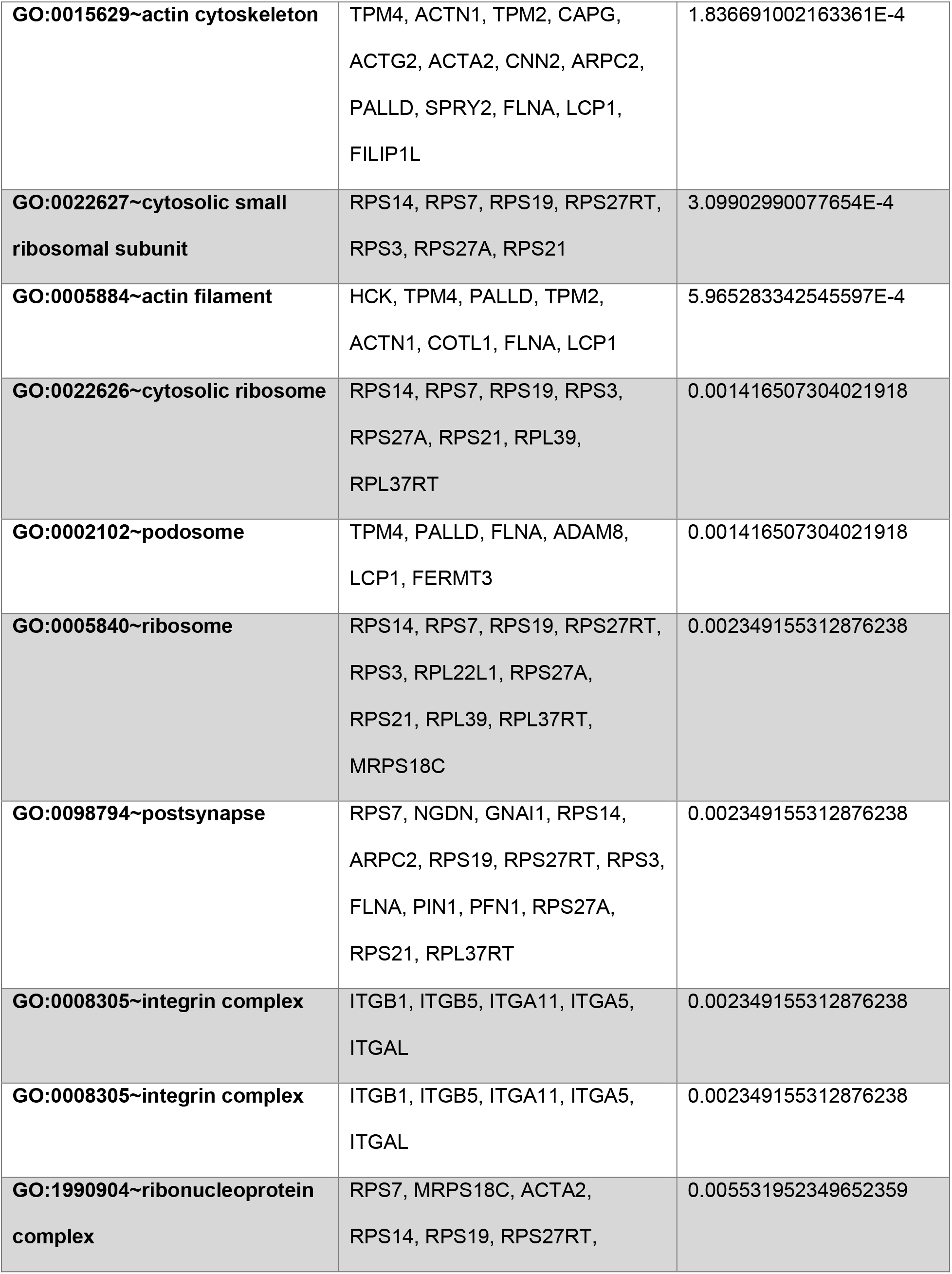
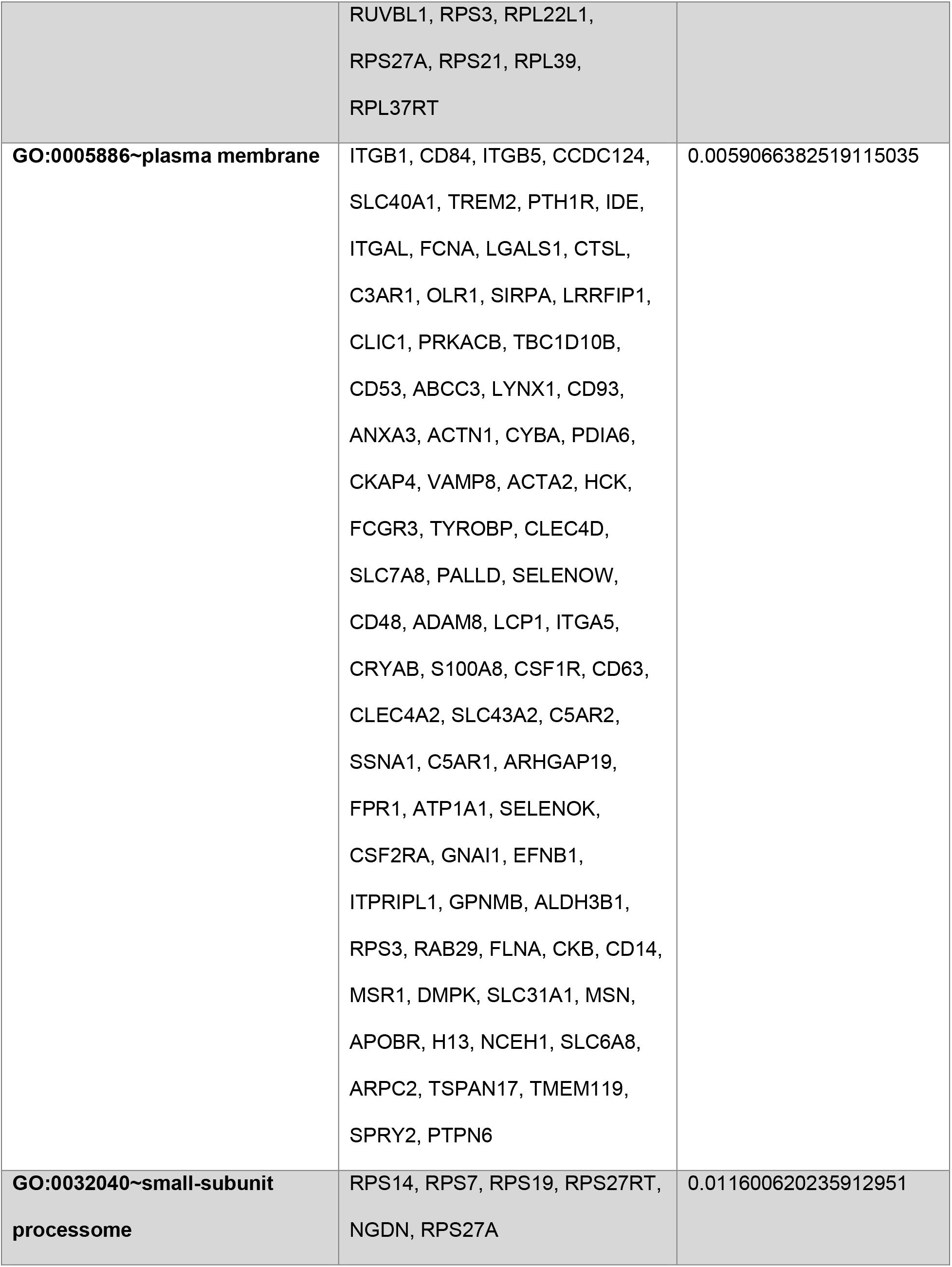
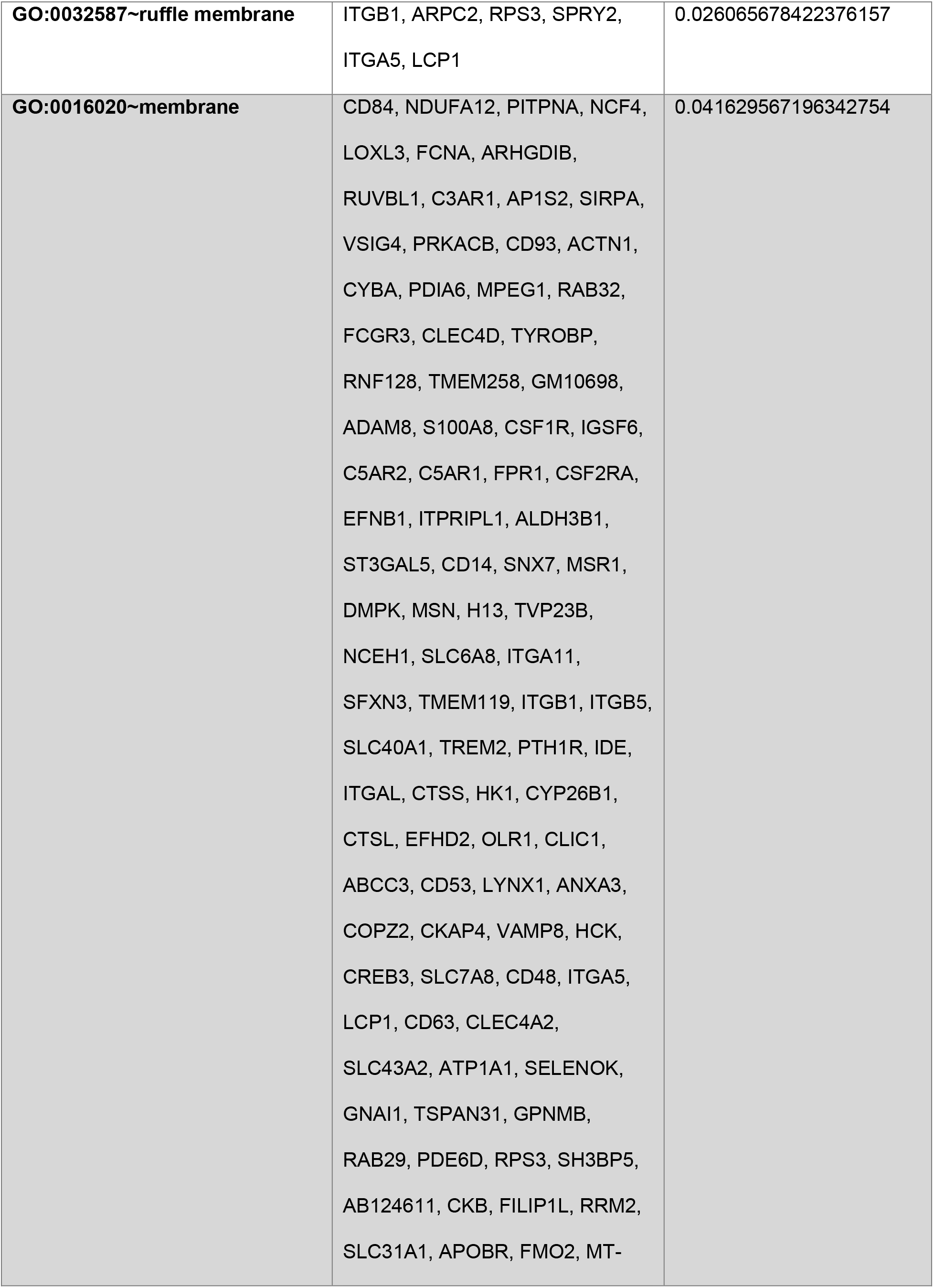
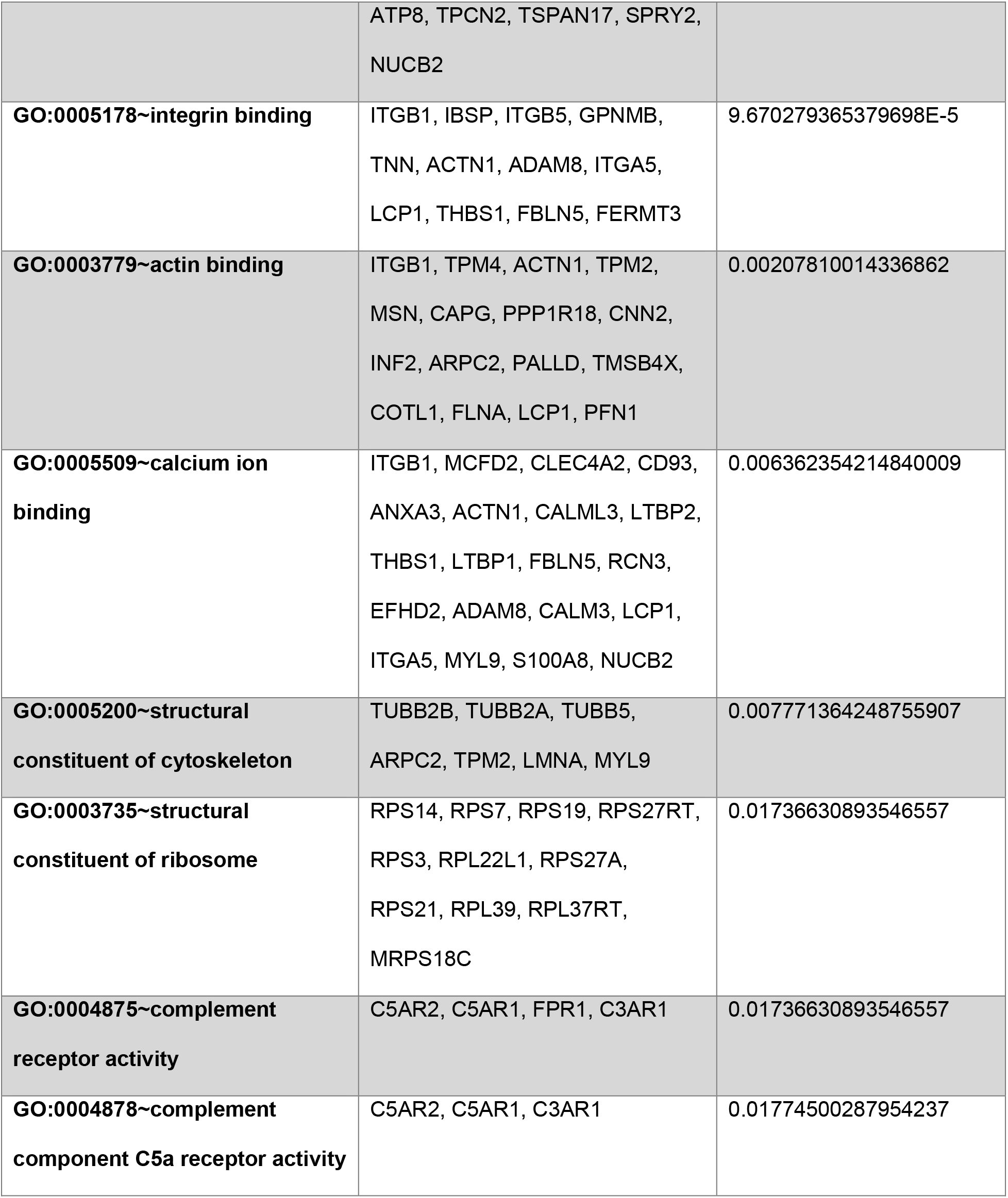
DAVID Gene Ontology (GO) enrichment analysis of genes upregulated in adipogenic medium following LIV treatment compared with untreated controls. Genes included in the analysis exhibited at least a 25% increase in expression in the +LIV group relative to the −LIV group.

| **Term** | **Genes** | **Benjamini** |
|---|---|---|
| **GO:0007160~cell-matrix adhesion** | ITGB1, CD63, ITGB5, TNN, ITGA11, SIRPA, ADAM8, ITGA5, ITGAL, FERMT3 | 1.4069023865197444E-4 |
| **GO:0007159~leukocyte cell-cell adhesion** | ITGB1, MSN, OLR1, ITGA5, ITGAL, FERMT3 | 0.003301596905342356 |
| **GO:0002684~positive regulation of immune system process** | TYROBP, C5AR2, C5AR1, C3AR1, THBS1 | 0.003301596905342356 |
| **GO:0016192~vesicle-mediated transport** | MCFD2, TRAPPC1, COPZ2, APOBR, TVP23B, RAB32, VAMP8, HCK, FCGR3, AP1S2, RAB29, GM10698, OLR1 | 0.004043405290683905 |
| **GO:0002181~cytoplasmic translation** | RPS14, RPS7, RPS19, RPS3, RPL22L1, RPS27A, RPS21, RPL39, RPL37RT | 0.00699042891755760​6 |
| **GO:0007229~integrin-mediated signaling pathway** | ITGB1, TYROBP, ITGB5, ITGA11, ADAM8, ITGA5, ITGAL, FERMT3 | 0.007299971201549757​5 |
| **GO:0006412~translation** | RPS7, MRPS18C, RBM3, LARS2, RPS14, RPS19, RPS27RT, RPS3, RPL22L1, RPS27A, RPS21, RPL39, RPL37RT | 0.010457571380504573 |
| **GO:0031667~response to nutrient levels** | PRL2C5, COL1A1, PRL2C3, GSTP1, LEP, CYBA, PRL2C2 | 0.026142627555612873 |
| **GO:0038178~complement component C5a signaling pathway** | C5AR2, C5AR1, C3AR1 | 0.03324781899101762 |
| **GO:0002430~complement receptor mediated signaling pathway** | C5AR2, C5AR1, FPR1, C3AR1 | 0.03389548879733518 |
| **GO:0098609~cell-cell adhesion** | ITGB1, LGALS1, SRPX2, ITGB5, CD93, ITGA11, ADAM8, ITGA5, ITGAL | 0.0461688201017277 |
| **GO:0005737~cytoplasm** | PITPNA, NCF4, LOXL3, CALML3, ACTG2, CNDP2, RPS14, TUBB5, LGALS1, RPS19, ARHGDIB, RUVBL1, COTL1, PRKACB, RPL39, TBC1D10B, RPS7, TPM4, ACTN1, TPM2, CTNNBIP1, NGDN, CIRBP, DDX10, CYBA, RAB32, AKR1C12, LARS2, DOK1, SELENOM, RNF128, TMEM258, FEZ2, SELENOW, ADAM8, PFN1, MYL9, S100A8, ARHGAP19, FPR1, EFNB1, ALDH3B1, SNX7, TRAPPC1, DMPK, GOT1, TBCC, MSN, SERPINA3G, COL1A1, ARPC2, ID1, TMEM119, PIN1, CALM3, RPS21, GALK1, LGMN, ITGB1, CUEDC2, CCDC124, FHL2, PCTP, IDE, PPP1R18, HK1, RBM3, CYP26B1, CTSL, TMSB4X, LRRFIP1, CLIC1, DUSP1, ANXA3, COPZ2, PTGR1, CKAP4, VAMP8, ACTA2, HCK, CREB3, TUBB2B, INF2, TUBB2A, PALLD, ALDH1A2, POLR1C, LCP1, CRYAB, SLFN2, GSTP1, SSNA1, CAPG, GNAI1, SRPX2, SERTAD1, SPOP, RAB29, PDE6D, RPS3, SH3BP5, FLNA, CKB, EXOSC3, FILIP1L, RPS27A, RRM2, ELP3, KLF6, GSTA3, IMPDH2, LEP, SPRY2, PTPN6, RPL37RT, NUCB2, PF4 | 1.0693891679573129E-10 |
| **GO:0005829~cytosol** | CUEDC2, CCDC124, PITPNA, NCF4, SLC40A1, PCTP, PTH1R, IDE, CNDP2, HK1, HK3, RPS14, TUBB5, LGALS1, RPS19, TMSB4X, RUVBL1, ARHGDIB, COTL1, LRRFIP1, RPL39, PRKACB, CTNNBL1, TBC1D10B, RPS7, CTNNBIP1, COPZ2, PTGR1, VAMP8, AKR1C12, ACTA2, DOK1, HCK, CREB3, PALLD, POLR1C, SELENOW, LCP1, PFN1, CRYAB, S100A8, SLFN2, GSTP1, GNAI1, ALDH3B1, PDE6D, RPS3, RAB29, FLNA, CKB, EXOSC3, RPS27A, RRM2, DMPK, GOT1, TBCC, MSN, ELP3, TPCN2, KLF6, GSTA3, LEP, IMPDH2, RPS27RT, SPRY2, PIN1, RPL22L1, PTPN6, RPS21, RPL37RT, GALK1 | 7.405920513314523E-6 |
| **GO:0005576~extracellular region** | CD63, MCFD2, SERPINE1, LOXL3, LTBP2, TREM2, IDE, METRNL, THBS1, PRSS23, CXCL14, LTBP1, FBLN5, CTSS, PRL2C5, FCNA, PRL2C3, LGALS1, SRPX2, IBSP, CNPY4, CTSL, TNN, OLR1, SDF2, TIMP1, CD14, PRL2C2, ANGPT4, CTLA2A, MANF, COL1A1, RBP4, LOX, LEP, TMEM119, CD48, PFN1, CRYAB, NBL1, NUCB2, S100A8, LGMN, PF4 | 4.475784167543283E-5 |
| **GO:0005615~extracellular space** | CD63, SERPINE1, LOXL3, IDE, METRNL, THBS1, PRSS23, CXCL14, FBLN5, CTSS, PRL2C5, FCNA, PRL2C3, LGALS1, SRPX2, IBSP, CTSL, TNN, TIMP1, CKB, CD14, RPL39, PRL2C2, ANGPT4, SERPINA3G, PDIA6, MANF, COL1A1, RBP4, LOX, LEP, ADPRH, NBL1, NUCB2, S100A8, PF4 | 1.0377717843316774E-4 |
| **GO:0015629~actin cytoskeleton** | TPM4, ACTN1, TPM2, CAPG, ACTG2, ACTA2, CNN2, ARPC2, PALLD, SPRY2, FLNA, LCP1, FILIP1L | 1.836691002163361E-4 |
| **GO:0022627~cytosolic small ribosomal subunit** | RPS14, RPS7, RPS19, RPS27RT, RPS3, RPS27A, RPS21 | 3.09902990077654E-4 |
| **GO:0005884~actin filament** | HCK, TPM4, PALLD, TPM2, ACTN1, COTL1, FLNA, LCP1 | 5.965283342545597E-4 |
| **GO:0022626~cytosolic ribosome** | RPS14, RPS7, RPS19, RPS3, RPS27A, RPS21, RPL39, RPL37RT | 0.001416507304021918 |
| **GO:0002102~podosome** | TPM4, PALLD, FLNA, ADAM8, LCP1, FERMT3 | 0.001416507304021918 |
| **GO:0005840~ribosome** | RPS14, RPS7, RPS19, RPS27RT, RPS3, RPL22L1, RPS27A, RPS21, RPL39, RPL37RT, MRPS18C | 0.002349155312876238 |
| **GO:0098794~postsynapse** | RPS7, NGDN, GNAI1, RPS14, ARPC2, RPS19, RPS27RT, RPS3, FLNA, PIN1, PFN1, RPS27A, RPS21, RPL37RT | 0.002349155312876238 |
| **GO:0008305~integrin complex** | ITGB1, ITGB5, ITGA11, ITGA5, ITGAL | 0.002349155312876238 |
| **GO:0008305~integrin complex** | ITGB1, ITGB5, ITGA11, ITGA5, ITGAL | 0.002349155312876238 |
| **GO:1990904~ribonucleoprotein complex** | RPS7, MRPS18C, ACTA2, RPS14, RPS19, RPS27RT, RUVBL1, RPS3, RPL22L1, RPS27A, RPS21, RPL39, RPL37RT | 0.00553195234965235​9 |
| **GO:0005886~plasma membrane** | ITGB1, CD84, ITGB5, CCDC124, SLC40A1, TREM2, PTH1R, IDE, ITGAL, FCNA, LGALS1, CTSL, C3AR1, OLR1, SIRPA, LRRFIP1, CLIC1, PRKACB, TBC1D10B, CD53, ABCC3, LYNX1, CD93, ANXA3, ACTN1, CYBA, PDIA6, CKAP4, VAMP8, ACTA2, HCK, FCGR3, TYROBP, CLEC4D, SLC7A8, PALLD, SELENOW, CD48, ADAM8, LCP1, ITGA5, CRYAB, S100A8, CSF1R, CD63, CLEC4A2, SLC43A2, C5AR2, SSNA1, C5AR1, ARHGAP19, FPR1, ATP1A1, SELENOK, CSF2RA, GNAI1, EFNB1, ITPRIPL1, GPNMB, ALDH3B1, RPS3, RAB29, FLNA, CKB, CD14, MSR1, DMPK, SLC31A1, MSN, APOBR, H13, NCEH1, SLC6A8, ARPC2, TSPAN17, TMEM119, SPRY2, PTPN6 | 0.005906638251911503​5 |
| **GO:0032040~small-subunit processome** | RPS14, RPS7, RPS19, RPS27RT, NGDN, RPS27A | 0.011600620235912951 |
| **GO:0032587~ruffle membrane** | ITGB1, ARPC2, RPS3, SPRY2, ITGA5, LCP1 | 0.026065678422376157 |
| **GO:0016020~membrane** | CD84, NDUFA12, PITPNA, NCF4, LOXL3, FCNA, ARHGDIB, RUVBL1, C3AR1, AP1S2, SIRPA, VSIG4, PRKACB, CD93, ACTN1, CYBA, PDIA6, MPEG1, RAB32, FCGR3, CLEC4D, TYROBP, RNF128, TMEM258, GM10698, ADAM8, S100A8, CSF1R, IGSF6, C5AR2, C5AR1, FPR1, CSF2RA, EFNB1, ITPRIPL1, ALDH3B1, ST3GAL5, CD14, SNX7, MSR1, DMPK, MSN, H13, TVP23B, NCEH1, SLC6A8, ITGA11, SFXN3, TMEM119, ITGB1, ITGB5, SLC40A1, TREM2, PTH1R, IDE, ITGAL, CTSS, HK1, CYP26B1, CTSL, EFHD2, OLR1, CLIC1, ABCC3, CD53, LYNX1, ANXA3, COPZ2, CKAP4, VAMP8, HCK, CREB3, SLC7A8, CD48, ITGA5, LCP1, CD63, CLEC4A2, SLC43A2, ATP1A1, SELENOK, GNAI1, TSPAN31, GPNMB, RAB29, PDE6D, RPS3, SH3BP5, AB124611, CKB, FILIP1L, RRM2, SLC31A1, APOBR, FMO2, MT-ATP8, TPCN2, TSPAN17, SPRY2, NUCB2 | 0.04162956719634275​4 |
| **GO:0005178~integrin binding** | ITGB1, IBSP, ITGB5, GPNMB, TNN, ACTN1, ADAM8, ITGA5, LCP1, THBS1, FBLN5, FERMT3 | 9.670279365379698E-5 |
| **GO:0003779~actin binding** | ITGB1, TPM4, ACTN1, TPM2, MSN, CAPG, PPP1R18, CNN2, INF2, ARPC2, PALLD, TMSB4X, COTL1, FLNA, LCP1, PFN1 | 0.00207810014336862 |
| **GO:0005509~calcium ion binding** | ITGB1, MCFD2, CLEC4A2, CD93, ANXA3, ACTN1, CALML3, LTBP2, THBS1, LTBP1, FBLN5, RCN3, EFHD2, ADAM8, CALM3, LCP1, ITGA5, MYL9, S100A8, NUCB2 | 0.006362354214840009 |
| **GO:0005200~structural constituent of cytoskeleton** | TUBB2B, TUBB2A, TUBB5, ARPC2, TPM2, LMNA, MYL9 | 0.007771364248755907 |
| **GO:0003735~structural constituent of ribosome** | RPS14, RPS7, RPS19, RPS27RT, RPS3, RPL22L1, RPS27A, RPS21, RPL39, RPL37RT, MRPS18C | 0.01736630893546557 |
| **GO:0004875~complement receptor activity** | C5AR2, C5AR1, FPR1, C3AR1 | 0.01736630893546557 |
| **GO:0004878~complement component C5a receptor activity** | C5AR2, C5AR1, C3AR1 | 0.01774500287954237 |

**Table S3.**
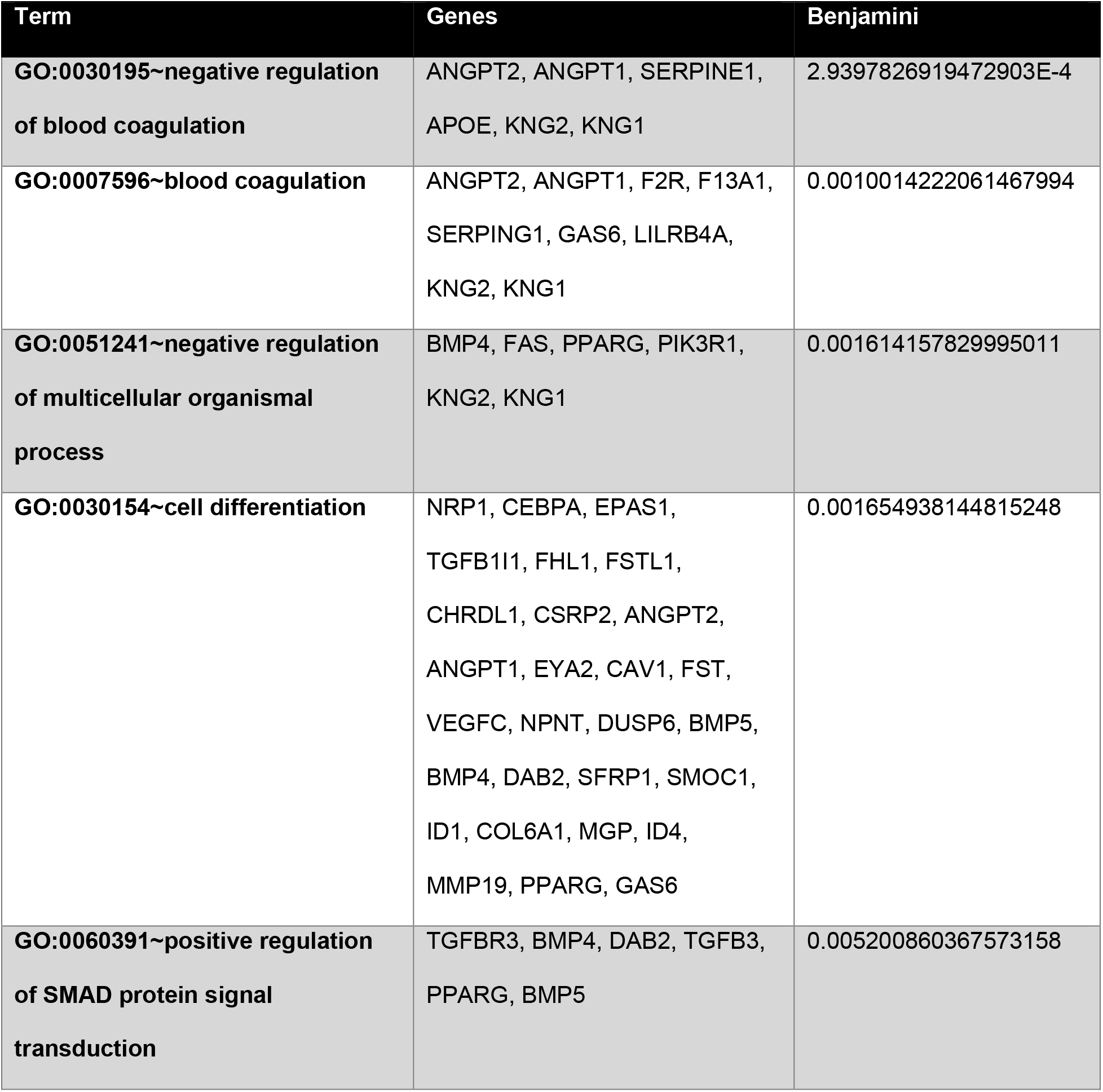
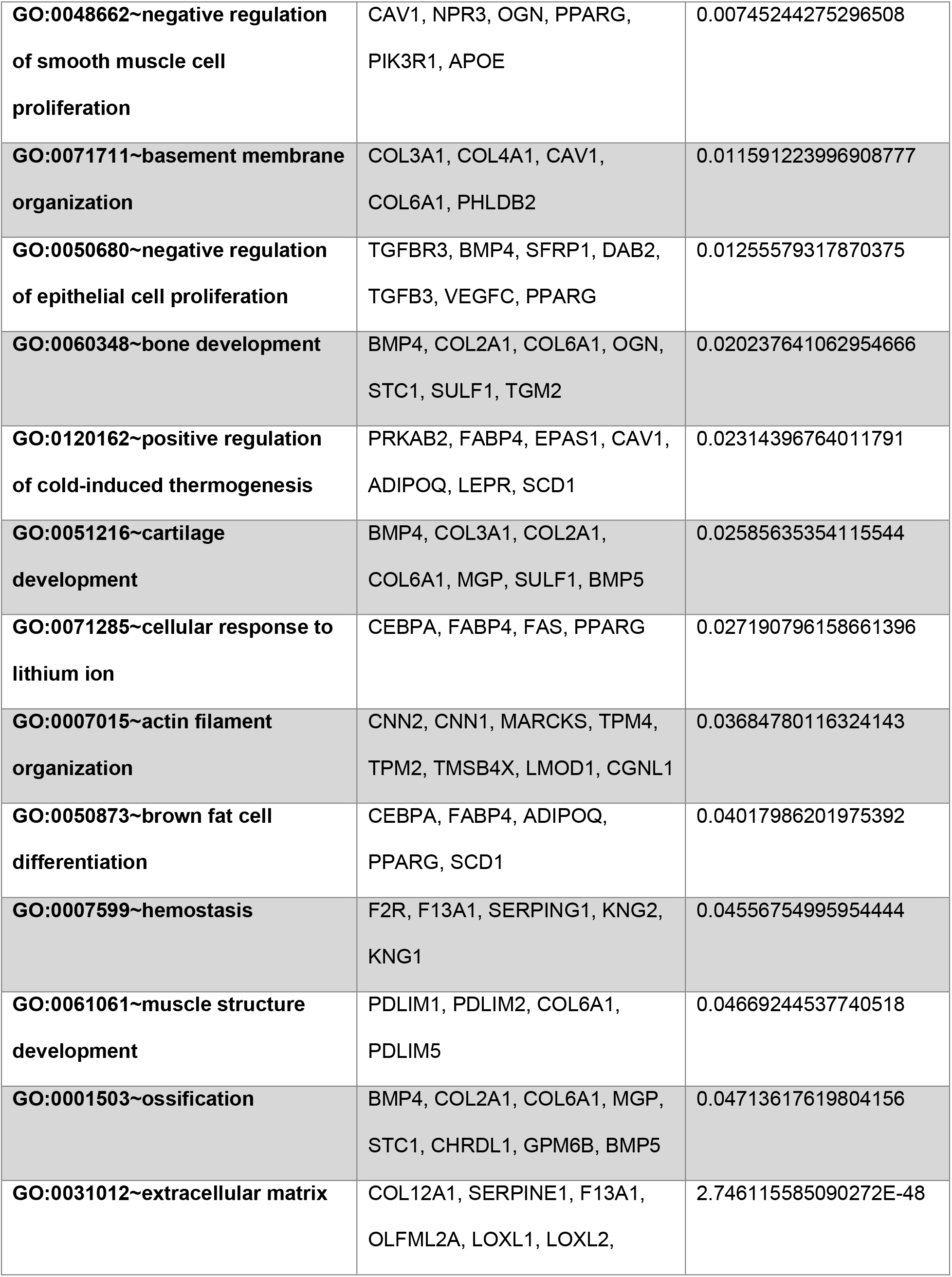
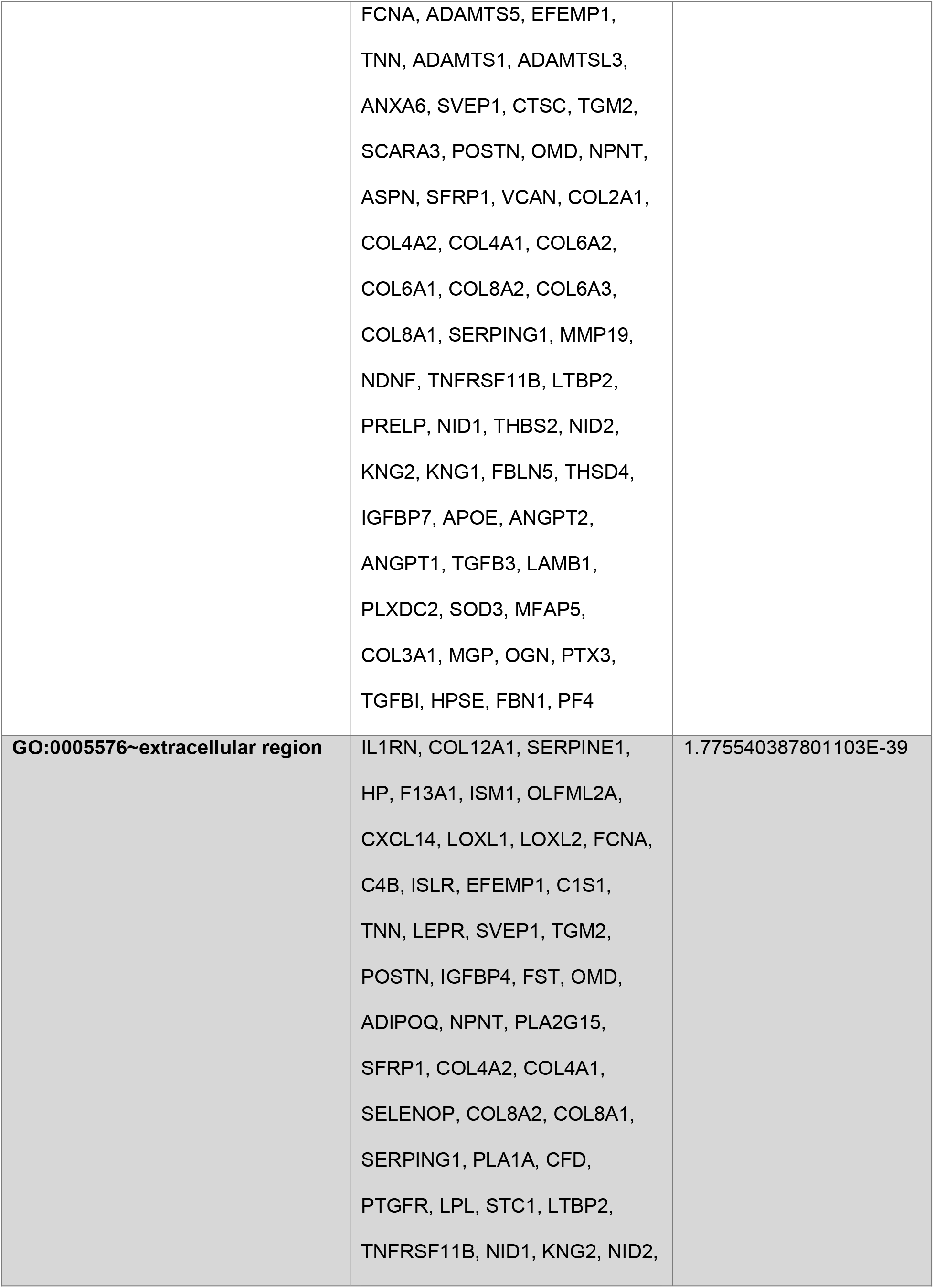
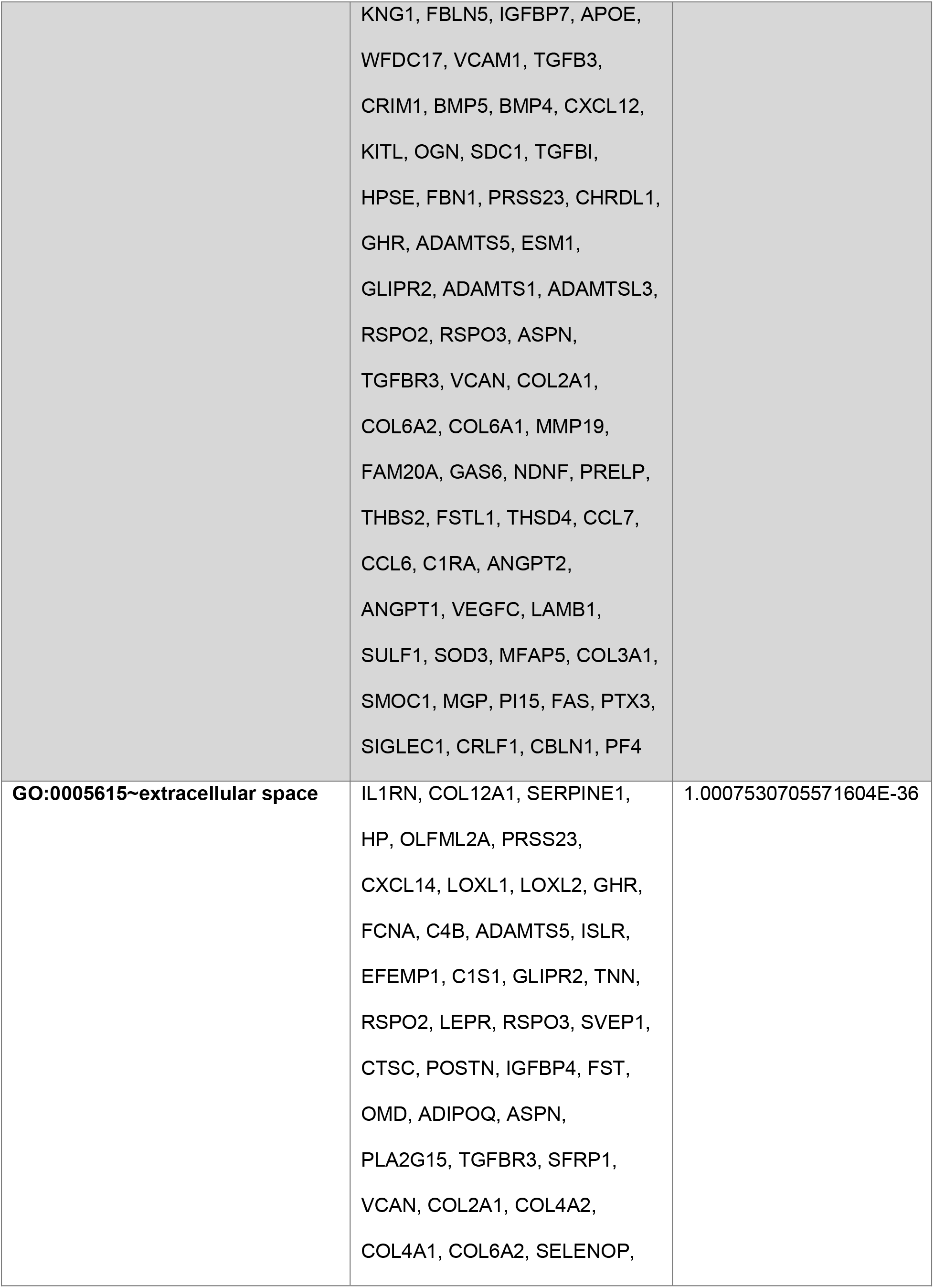
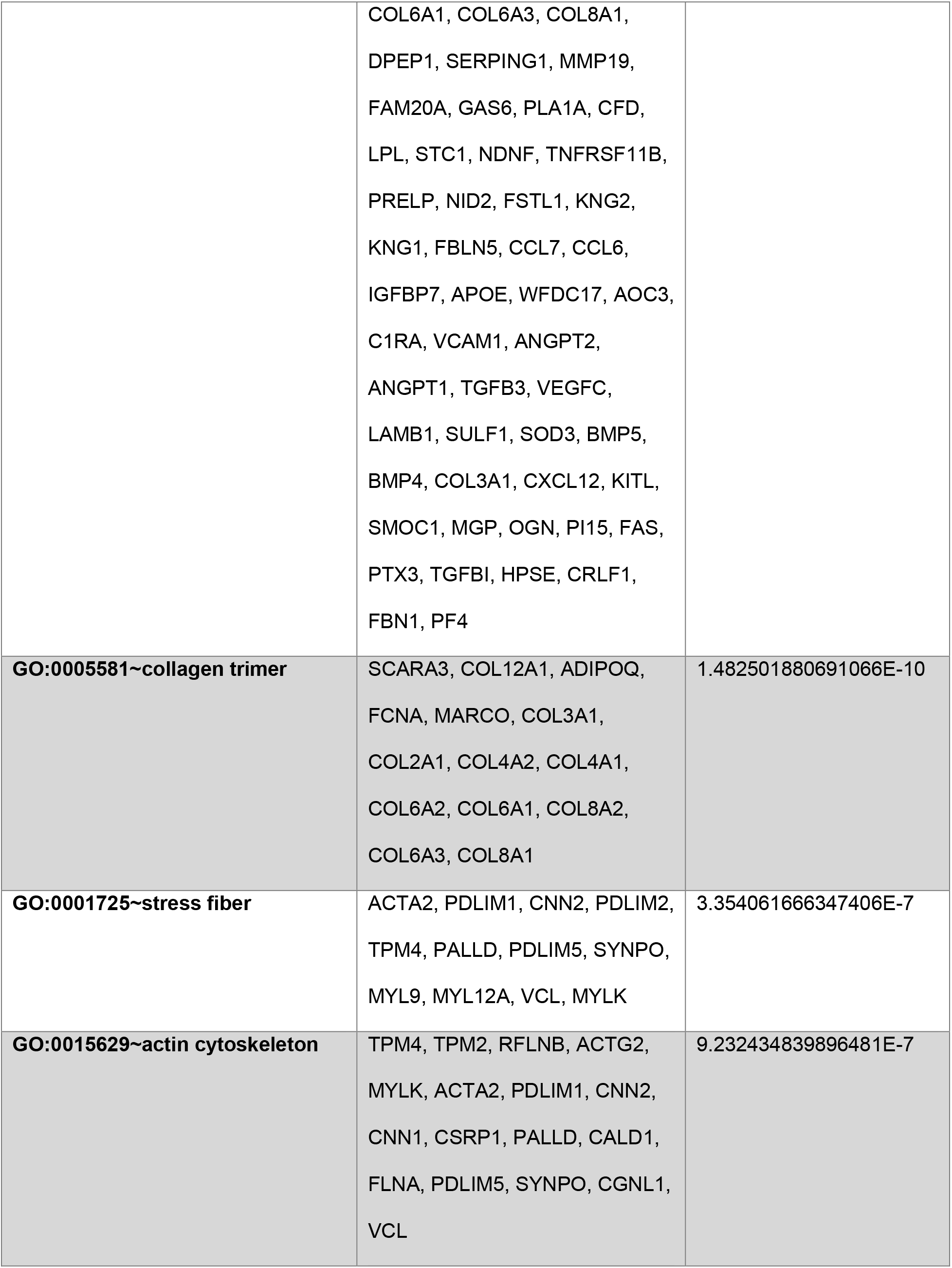
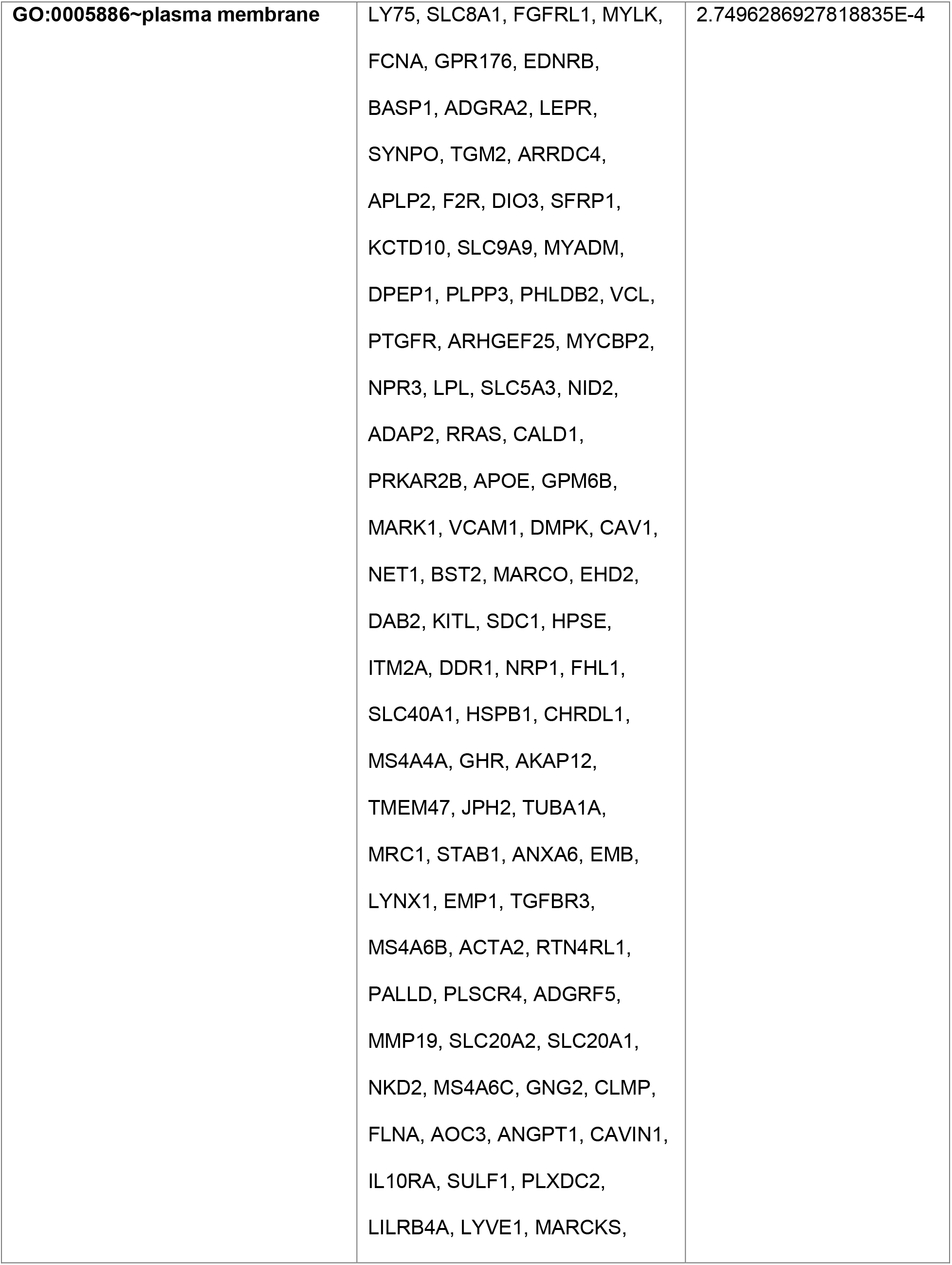
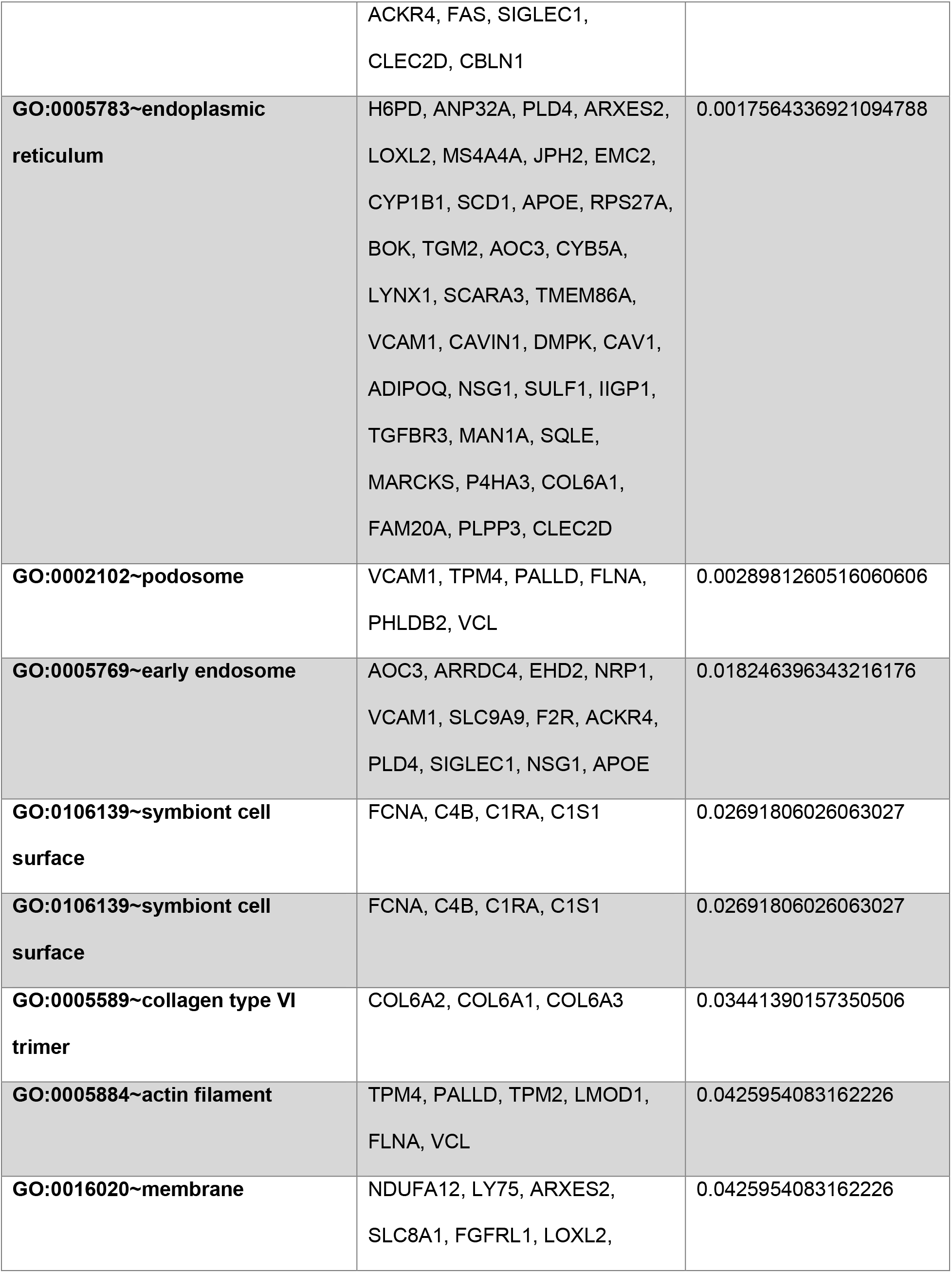
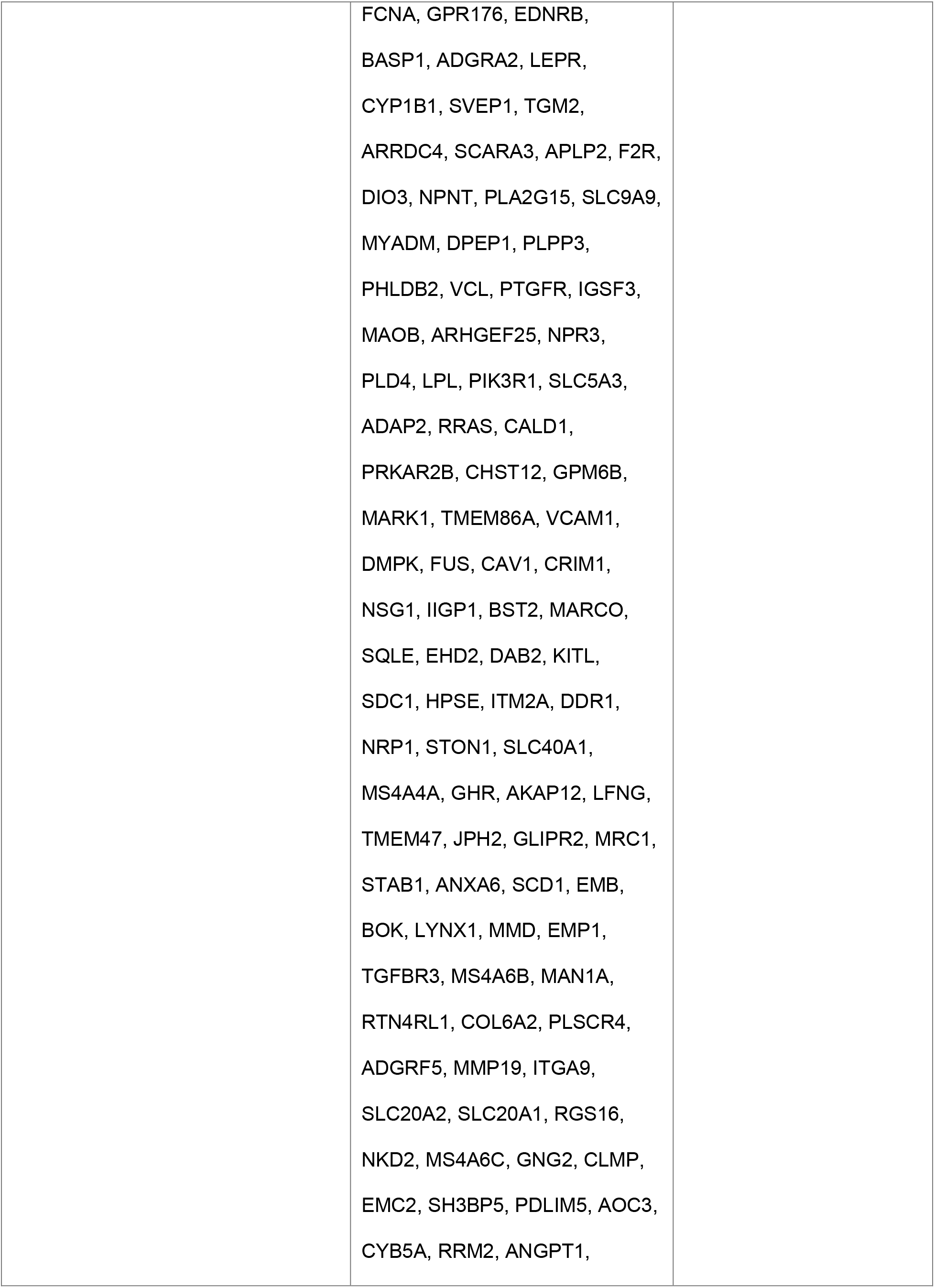
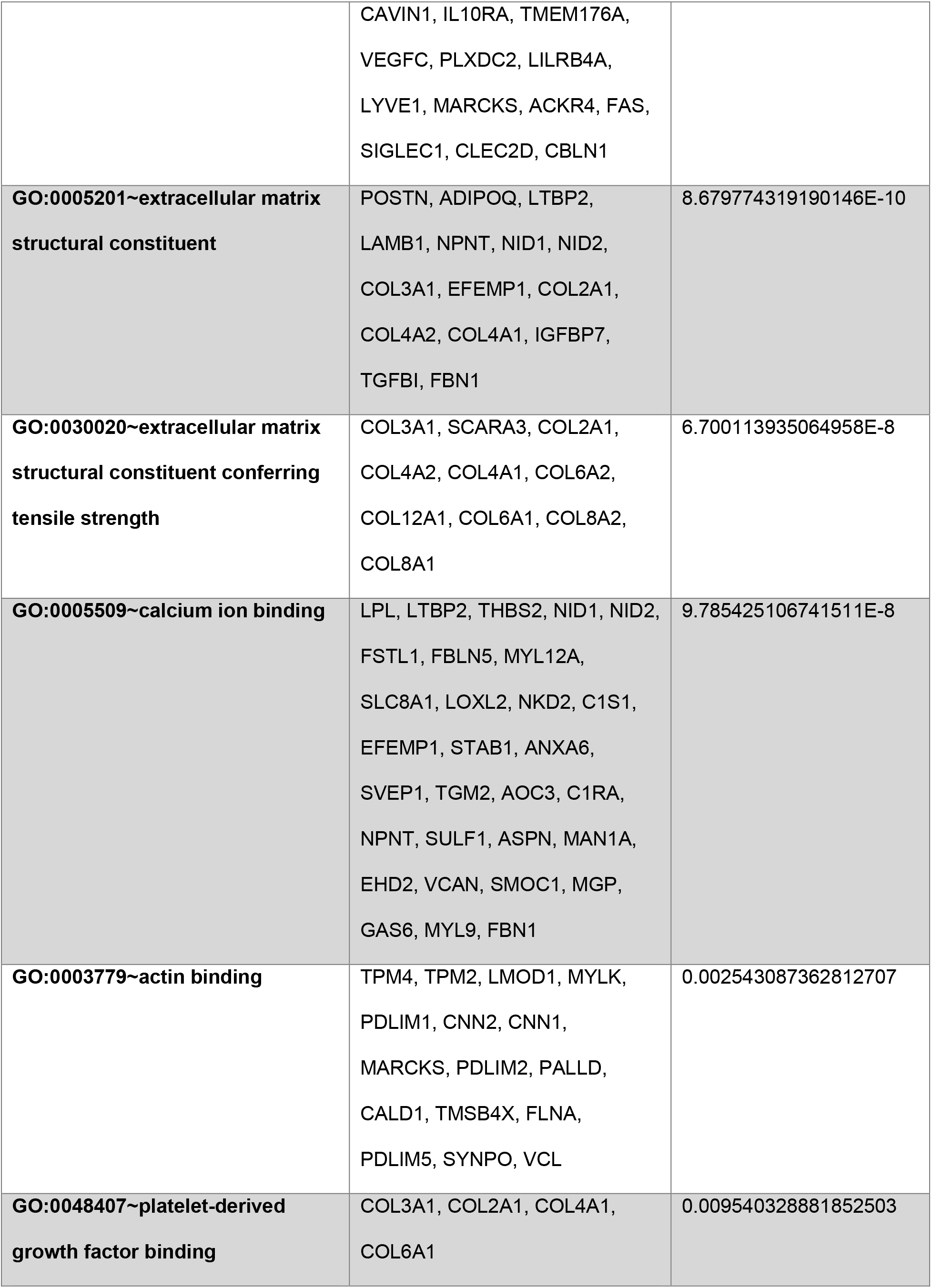
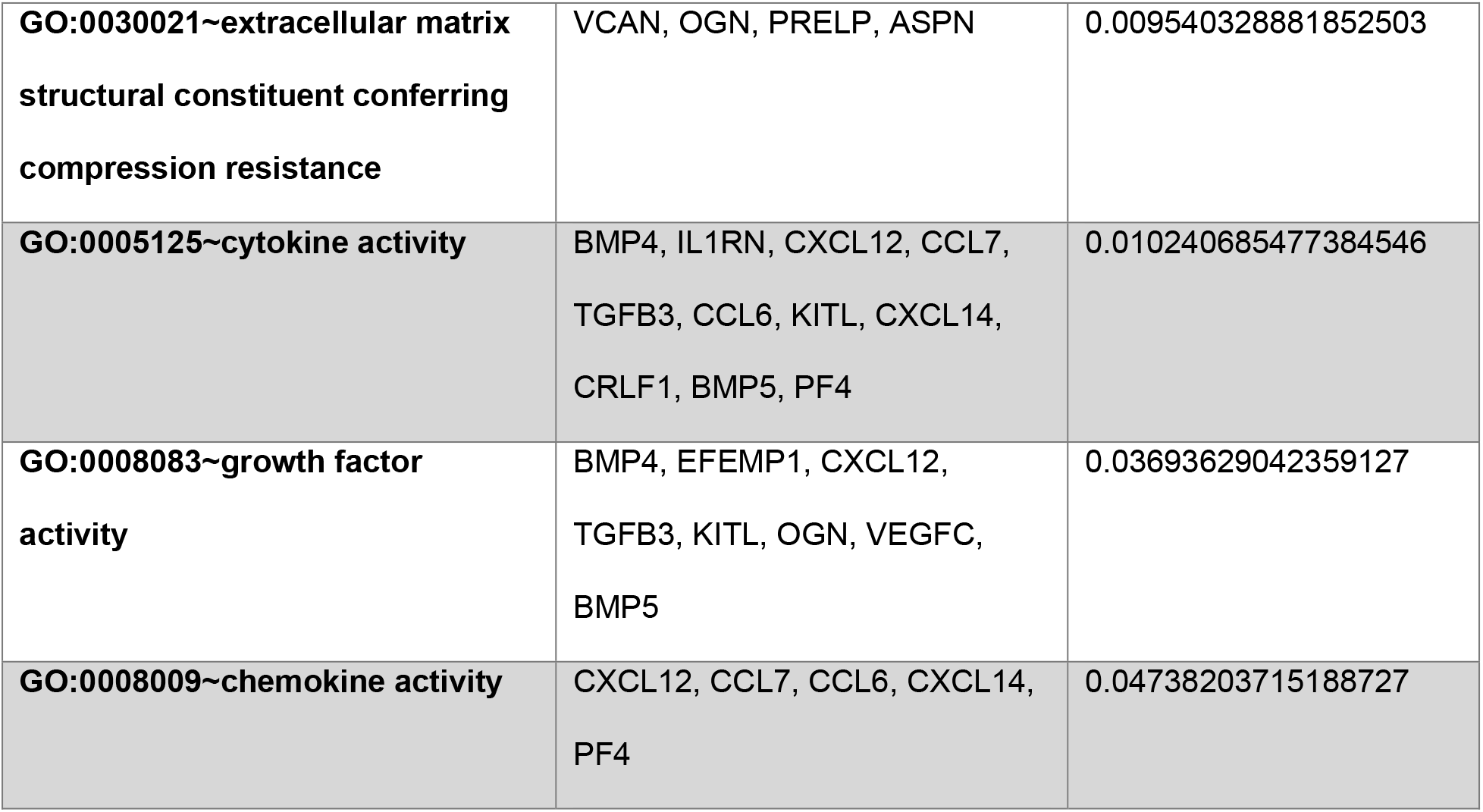
DAVID Gene Ontology (GO) enrichment analysis of genes downregulated in osteogenic medium following LIV treatment compared with untreated controls. Genes included in the analysis exhibited at least a 25% decrease in expression in the +LIV group relative to the −LIV group.

| **Term** | **Genes** | **Benjamini** |
|---|---|---|
| **GO:0030195~negative regulation of blood coagulation** | ANGPT2, ANGPT1, SERPINE1, APOE, KNG2, KNG1 | 2.9397826919472903E-4 |
| **GO:0007596~blood coagulation** | ANGPT2, ANGPT1, F2R, F13A1, SERPING1, GAS6, LILRB4A, KNG2, KNG1 | 0.0010014222061467994 |
| **GO:0051241~negative regulation of multicellular organismal process** | BMP4, FAS, PPARG, PIK3R1, KNG2, KNG1 | 0.001614157829995011 |
| **GO:0030154~cell differentiation** | NRP1, CEBPA, EPAS1, TGFB1I1, FHL1, FSTL1, CHRDL1, CSRP2, ANGPT2, ANGPT1, EYA2, CAV1, FST, VEGFC, NPNT, DUSP6, BMP5, BMP4, DAB2, SFRP1, SMOC1, ID1, COL6A1, MGP, ID4, MMP19, PPARG, GAS6 | 0.001654938144815248 |
| **GO:0060391~positive regulation of SMAD protein signal transduction** | TGFBR3, BMP4, DAB2, TGFB3, PPARG, BMP5 | 0.005200860367573158 |
| **GO:0048662~negative regulation of smooth muscle cell proliferation** | CAV1, NPR3, OGN, PPARG, PIK3R1, APOE | 0.00745244275296508 |
| **GO:0071711~basement membrane organization** | COL3A1, COL4A1, CAV1, COL6A1, PHLDB2 | 0.011591223996908777 |
| **GO:0050680~negative regulation of epithelial cell proliferation** | TGFBR3, BMP4, SFRP1, DAB2, TGFB3, VEGFC, PPARG | 0.01255579317870375 |
| **GO:0060348~bone development** | BMP4, COL2A1, COL6A1, OGN, STC1, SULF1, TGM2 | 0.020237641062954666 |
| **GO:0120162~positive regulation of cold-induced thermogenesis** | PRKAB2, FABP4, EPAS1, CAV1, ADIPOQ, LEPR, SCD1 | 0.02314396764011791 |
| **GO:0051216~cartilage development** | BMP4, COL3A1, COL2A1, COL6A1, MGP, SULF1, BMP5 | 0.02585635354115544 |
| **GO:0071285~cellular response to lithium ion** | CEBPA, FABP4, FAS, PPARG | 0.027190796158661396 |
| **GO:0007015~actin filament organization** | CNN2, CNN1, MARCKS, TPM4, TPM2, TMSB4X, LMOD1, CGNL1 | 0.03684780116324143 |
| **GO:0050873~brown fat cell differentiation** | CEBPA, FABP4, ADIPOQ, PPARG, SCD1 | 0.04017986201975392 |
| **GO:0007599~hemostasis** | F2R, F13A1, SERPING1, KNG2, KNG1 | 0.04556754995954444 |
| **GO:0061061~muscle structure development** | PDLIM1, PDLIM2, COL6A1, PDLIM5 | 0.04669244537740518 |
| **GO:0001503~ossification** | BMP4, COL2A1, COL6A1, MGP, STC1, CHRDL1, GPM6B, BMP5 | 0.04713617619804156 |
| **GO:0031012~extracellular matrix** | COL12A1, SERPINE1, F13A1, OLFML2A, LOXL1, LOXL2, FCNA, ADAMTS5, EFEMP1, TNN, ADAMTS1, ADAMTSL3, ANXA6, SVEP1, CTSC, TGM2, SCARA3, POSTN, OMD, NPNT, ASPN, SFRP1, VCAN, COL2A1, COL4A2, COL4A1, COL6A2, COL6A1, COL8A2, COL6A3, COL8A1, SERPING1, MMP19, NDNF, TNFRSF11B, LTBP2, PRELP, NID1, THBS2, NID2, KNG2, KNG1, FBLN5, THSD4, IGFBP7, APOE, ANGPT2, ANGPT1, TGFB3, LAMB1, PLXDC2, SOD3, MFAP5, COL3A1, MGP, OGN, PTX3, TGFBI, HPSE, FBN1, PF4 | 2.746115585090272E-48 |
| **GO:0005576~extracellular region** | IL1RN, COL12A1, SERPINE1, HP, F13A1, ISM1, OLFML2A, CXCL14, LOXL1, LOXL2, FCNA, C4B, ISLR, EFEMP1, C1S1, TNN, LEPR, SVEP1, TGM2, POSTN, IGFBP4, FST, OMD, ADIPOQ, NPNT, PLA2G15, SFRP1, COL4A2, COL4A1, SELENOP, COL8A2, COL8A1, SERPING1, PLA1A, CFD, PTGFR, LPL, STC1, LTBP2, TNFRSF11B, NID1, KNG2, NID2, KNG1, FBLN5, IGFBP7, APOE, WFDC17, VCAM1, TGFB3, CRIM1, BMP5, BMP4, CXCL12, KITL, OGN, SDC1, TGFBI, HPSE, FBN1, PRSS23, CHRDL1, GHR, ADAMTS5, ESM1, GLIPR2, ADAMTS1, ADAMTSL3, RSPO2, RSPO3, ASPN, TGFBR3, VCAN, COL2A1, COL6A2, COL6A1, MMP19, FAM20A, GAS6, NDNF, PRELP, THBS2, FSTL1, THSD4, CCL7, CCL6, C1RA, ANGPT2, ANGPT1, VEGFC, LAMB1, SULF1, SOD3, MFAP5, COL3A1, SMOC1, MGP, PI15, FAS, PTX3, SIGLEC1, CRLF1, CBLN1, PF4 | 1.775540387801103E-39 |
| **GO:0005615~extracellular space** | IL1RN, COL12A1, SERPINE1, HP, OLFML2A, PRSS23, CXCL14, LOXL1, LOXL2, GHR, FCNA, C4B, ADAMTS5, ISLR, EFEMP1, C1S1, GLIPR2, TNN, RSPO2, LEPR, RSPO3, SVEP1, CTSC, POSTN, IGFBP4, FST, OMD, ADIPOQ, ASPN, PLA2G15, TGFBR3, SFRP1, VCAN, COL2A1, COL4A2, COL4A1, COL6A2, SELENOP, COL6A1, COL6A3, COL8A1, DPEP1, SERPING1, MMP19, FAM20A, GAS6, PLA1A, CFD, LPL, STC1, NDNF, TNFRSF11B, PRELP, NID2, FSTL1, KNG2, KNG1, FBLN5, CCL7, CCL6, IGFBP7, APOE, WFDC17, AOC3, C1RA, VCAM1, ANGPT2, ANGPT1, TGFB3, VEGFC, LAMB1, SULF1, SOD3, BMP5, BMP4, COL3A1, CXCL12, KITL, SMOC1, MGP, OGN, PI15, FAS, PTX3, TGFBI, HPSE, CRLF1, FBN1, PF4 | 1.0007530705571604E-36 |
| **GO:0005581~collagen trimer** | SCARA3, COL12A1, ADIPOQ, FCNA, MARCO, COL3A1, COL2A1, COL4A2, COL4A1, COL6A2, COL6A1, COL8A2, COL6A3, COL8A1 | 1.482501880691066E-10 |
| **GO:0001725~stress fiber** | ACTA2, PDLIM1, CNN2, PDLIM2, TPM4, PALLD, PDLIM5, SYNPO, MYL9, MYL12A, VCL, MYLK | 3.354061666347406E-7 |
| **GO:0015629~actin cytoskeleton** | TPM4, TPM2, RFLNB, ACTG2, MYLK, ACTA2, PDLIM1, CNN2, CNN1, CSRP1, PALLD, CALD1, FLNA, PDLIM5, SYNPO, CGNL1, VCL | 9.232434839896481E-7 |
| **GO:0005886~plasma membrane** | LY75, SLC8A1, FGFRL1, MYLK, FCNA, GPR176, EDNRB, BASP1, ADGRA2, LEPR, SYNPO, TGM2, ARRDC4, APLP2, F2R, DIO3, SFRP1, KCTD10, SLC9A9, MYADM, DPEP1, PLPP3, PHLDB2, VCL, PTGFR, ARHGEF25, MYCBP2, NPR3, LPL, SLC5A3, NID2, ADAP2, RRAS, CALD1, PRKAR2B, APOE, GPM6B, MARK1, VCAM1, DMPK, CAV1, NET1, BST2, MARCO, EHD2, DAB2, KITL, SDC1, HPSE, ITM2A, DDR1, NRP1, FHL1, SLC40A1, HSPB1, CHRDL1, MS4A4A, GHR, AKAP12, TMEM47, JPH2, TUBA1A, MRC1, STAB1, ANXA6, EMB, LYNX1, EMP1, TGFBR3, MS4A6B, ACTA2, RTN4RL1, PALLD, PLSCR4, ADGRF5, MMP19, SLC20A2, SLC20A1, NKD2, MS4A6C, GNG2, CLMP, FLNA, AOC3, ANGPT1, CAVIN1, IL10RA, SULF1, PLXDC2, LILRB4A, LYVE1, MARCKS, ACKR4, FAS, SIGLEC1, CLEC2D, CBLN1 | 2.7496286927818835E-4 |
| **GO:0005783~endoplasmic reticulum** | H6PD, ANP32A, PLD4, ARXES2, LOXL2, MS4A4A, JPH2, EMC2, CYP1B1, SCD1, APOE, RPS27A, BOK, TGM2, AOC3, CYB5A, LYNX1, SCARA3, TMEM86A, VCAM1, CAVIN1, DMPK, CAV1, ADIPOQ, NSG1, SULF1, IIGP1, TGFBR3, MAN1A, SQLE, MARCKS, P4HA3, COL6A1, FAM20A, PLPP3, CLEC2D | 0.0017564336921094788 |
| **GO:0002102~podosome** | VCAM1, TPM4, PALLD, FLNA, PHLDB2, VCL | 0.0028981260516060606 |
| **GO:0005769~early endosome** | AOC3, ARRDC4, EHD2, NRP1, VCAM1, SLC9A9, F2R, ACKR4, PLD4, SIGLEC1, NSG1, APOE | 0.018246396343216176 |
| **GO:0106139~symbiont cell surface** | FCNA, C4B, C1RA, C1S1 | 0.02691806026063027 |
| **GO:0106139~symbiont cell surface** | FCNA, C4B, C1RA, C1S1 | 0.02691806026063027 |
| **GO:0005589~collagen type VI trimer** | COL6A2, COL6A1, COL6A3 | 0.03441390157350506 |
| **GO:0005884~actin filament** | TPM4, PALLD, TPM2, LMOD1, FLNA, VCL | 0.0425954083162226 |
| **GO:0016020~membrane** | NDUFA12, LY75, ARXES2, SLC8A1, FGFRL1, LOXL2, FCNA, GPR176, EDNRB, BASP1, ADGRA2, LEPR, CYP1B1, SVEP1, TGM2, ARRDC4, SCARA3, APLP2, F2R, DIO3, NPNT, PLA2G15, SLC9A9, MYADM, DPEP1, PLPP3, PHLDB2, VCL, PTGFR, IGSF3, MAOB, ARHGEF25, NPR3, PLD4, LPL, PIK3R1, SLC5A3, ADAP2, RRAS, CALD1, PRKAR2B, CHST12, GPM6B, MARK1, TMEM86A, VCAM1, DMPK, FUS, CAV1, CRIM1, NSG1, IIGP1, BST2, MARCO, SQLE, EHD2, DAB2, KITL, SDC1, HPSE, ITM2A, DDR1, NRP1, STON1, SLC40A1, MS4A4A, GHR, AKAP12, LFNG, TMEM47, JPH2, GLIPR2, MRC1, STAB1, ANXA6, SCD1, EMB, BOK, LYNX1, MMD, EMP1, TGFBR3, MS4A6B, MAN1A, RTN4RL1, COL6A2, PLSCR4, ADGRF5, MMP19, ITGA9, SLC20A2, SLC20A1, RGS16, NKD2, MS4A6C, GNG2, CLMP, EMC2, SH3BP5, PDLIM5, AOC3, CYB5A, RRM2, ANGPT1, CAVIN1, IL10RA, TMEM176A, VEGFC, PLXDC2, LILRB4A, LYVE1, MARCKS, ACKR4, FAS, SIGLEC1, CLEC2D, CBLN1 | 0.0425954083162226 |
| **GO:0005201~extracellular matrix structural constituent** | POSTN, ADIPOQ, LTBP2, LAMB1, NPNT, NID1, NID2, COL3A1, EFEMP1, COL2A1, COL4A2, COL4A1, IGFBP7, TGFBI, FBN1 | 8.679774319190146E-10 |
| **GO:0030020~extracellular matrix structural constituent conferring tensile strength** | COL3A1, SCARA3, COL2A1, COL4A2, COL4A1, COL6A2, COL12A1, COL6A1, COL8A2, COL8A1 | 6.700113935064958E-8 |
| **GO:0005509~calcium ion binding** | LPL, LTBP2, THBS2, NID1, NID2, FSTL1, FBLN5, MYL12A, SLC8A1, LOXL2, NKD2, C1S1, EFEMP1, STAB1, ANXA6, SVEP1, TGM2, AOC3, C1RA, NPNT, SULF1, ASPN, MAN1A, EHD2, VCAN, SMOC1, MGP, GAS6, MYL9, FBN1 | 9.785425106741511E-8 |
| **GO:0003779~actin binding** | TPM4, TPM2, LMOD1, MYLK, PDLIM1, CNN2, CNN1, MARCKS, PDLIM2, PALLD, CALD1, TMSB4X, FLNA, PDLIM5, SYNPO, VCL | 0.002543087362812707 |
| **GO:0048407~platelet-derived growth factor binding** | COL3A1, COL2A1, COL4A1, COL6A1 | 0.009540328881852503 |
| **GO:0030021~extracellular matrix structural constituent conferring compression resistance** | VCAN, OGN, PRELP, ASPN | 0.009540328881852503 |
| **GO:0005125~cytokine activity** | BMP4, IL1RN, CXCL12, CCL7, TGFB3, CCL6, KITL, CXCL14, CRLF1, BMP5, PF4 | 0.010240685477384546 |
| **GO:0008083~growth factor activity** | BMP4, EFEMP1, CXCL12, TGFB3, KITL, OGN, VEGFC, BMP5 | 0.03693629042359127 |
| **GO:0008009~chemokine activity** | CXCL12, CCL7, CCL6, CXCL14, PF4 | 0.04738203715188727 |

**Table S4.**
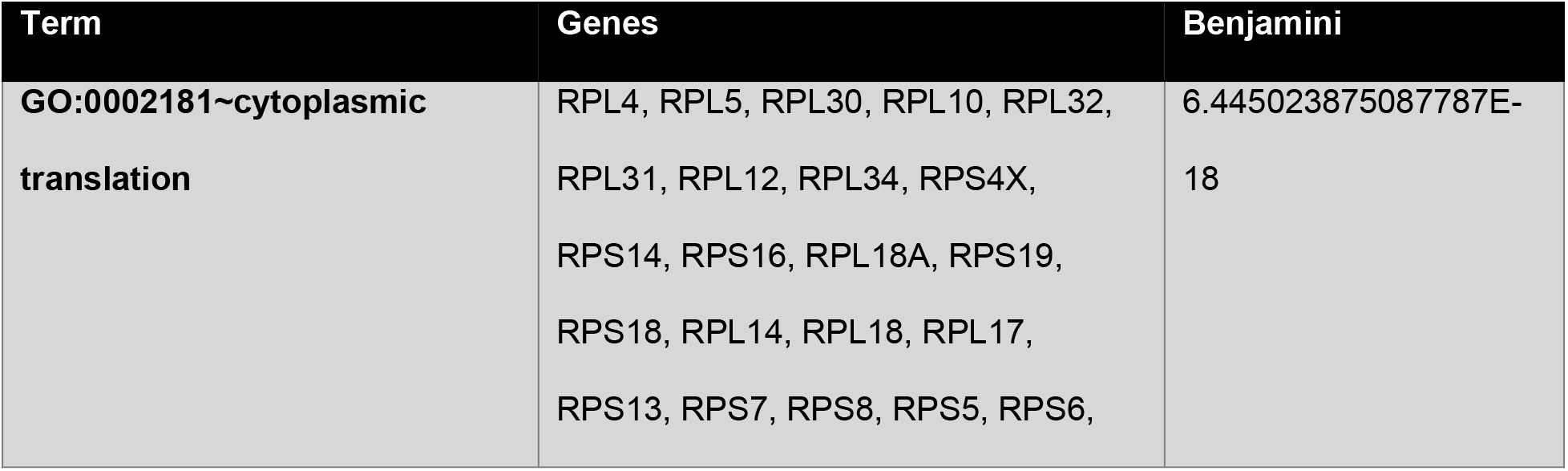
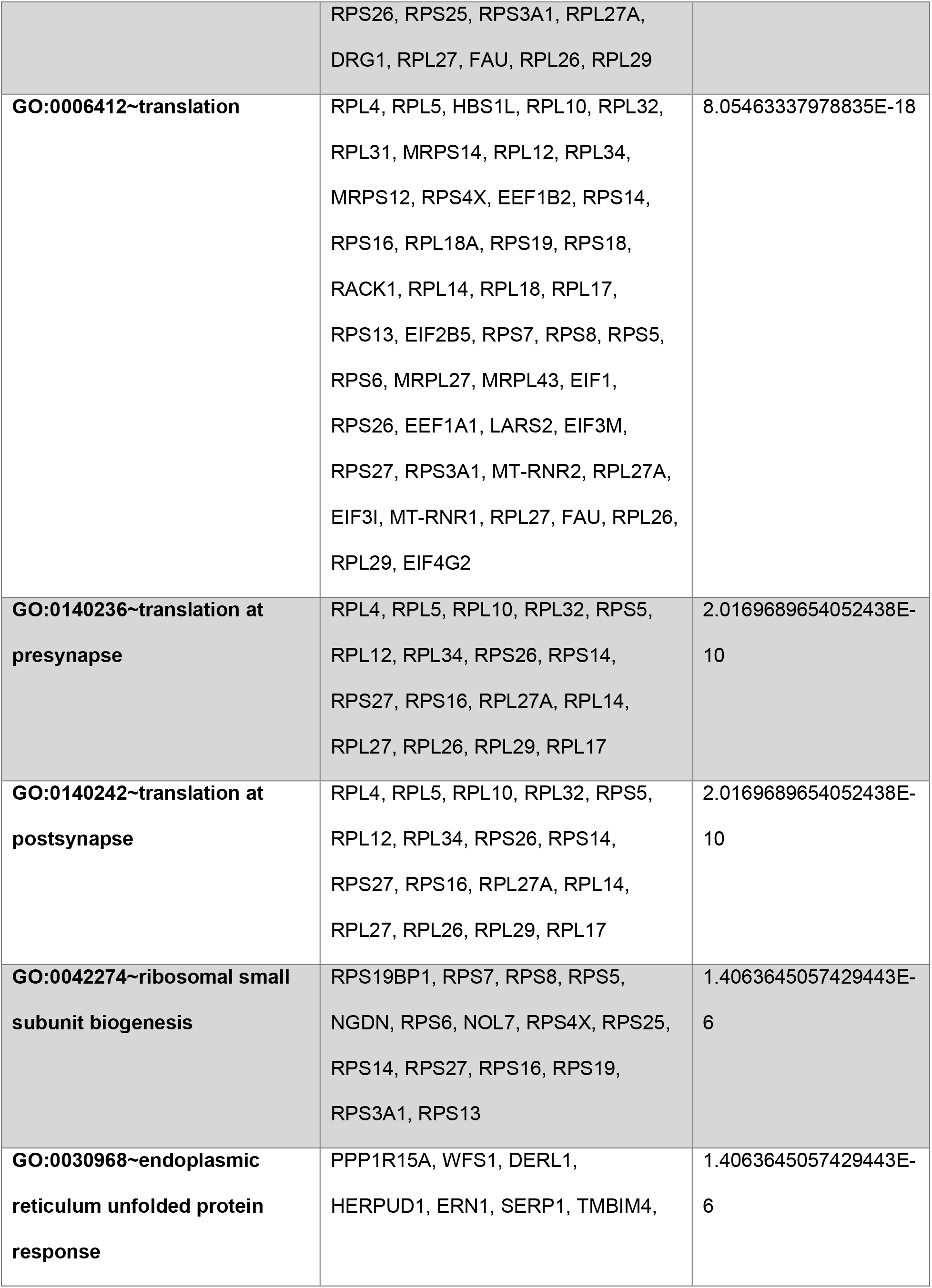
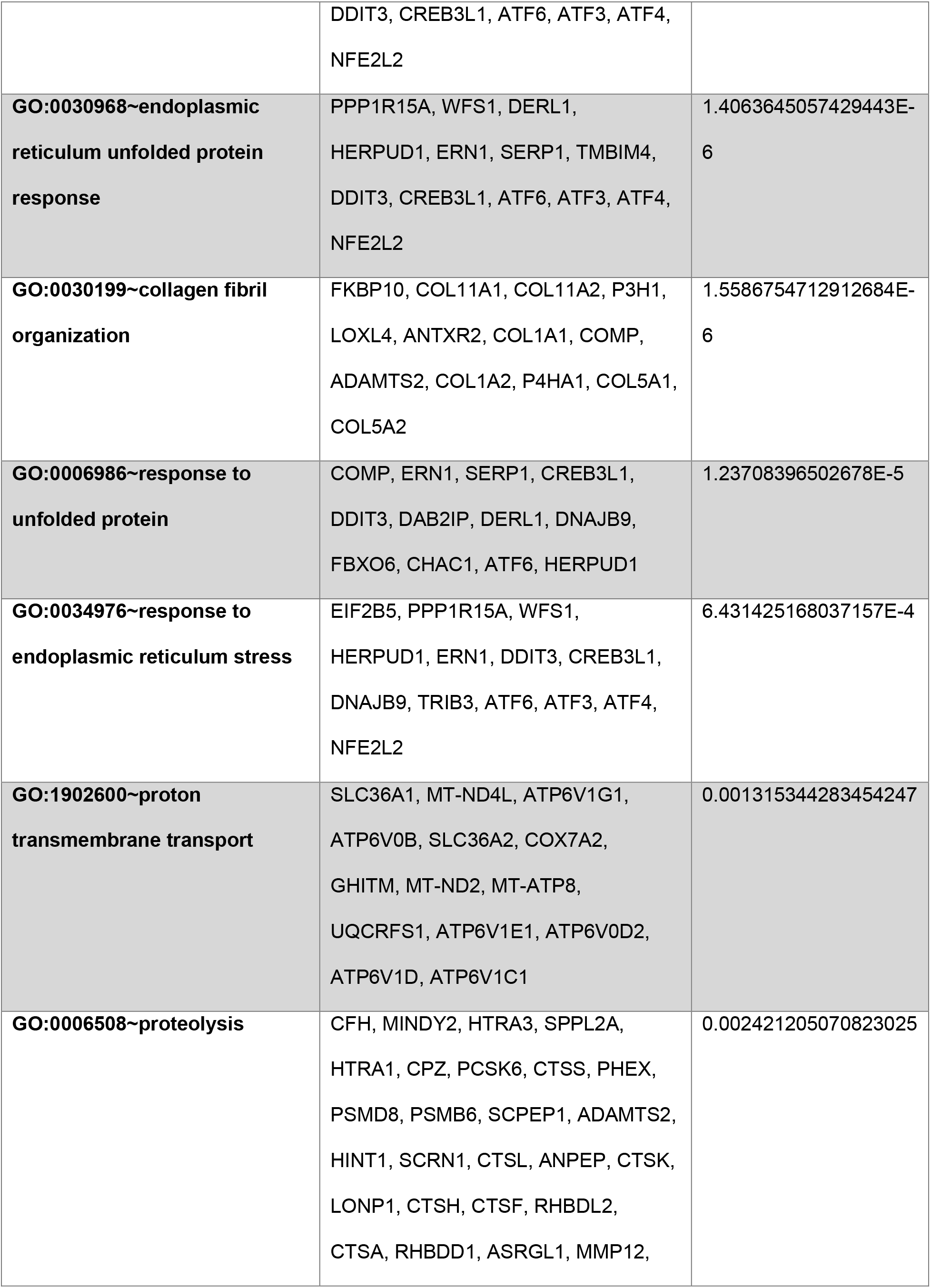
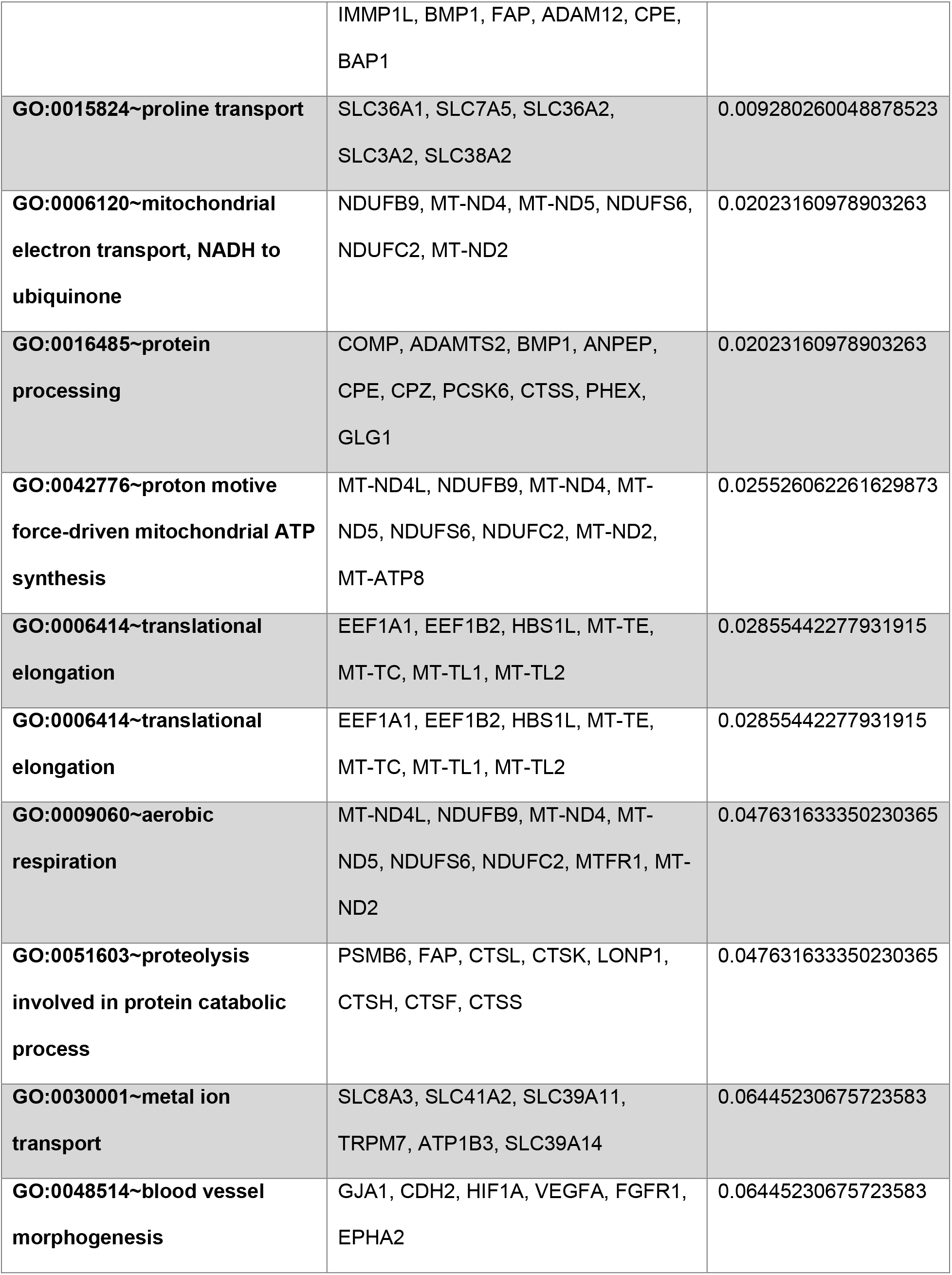
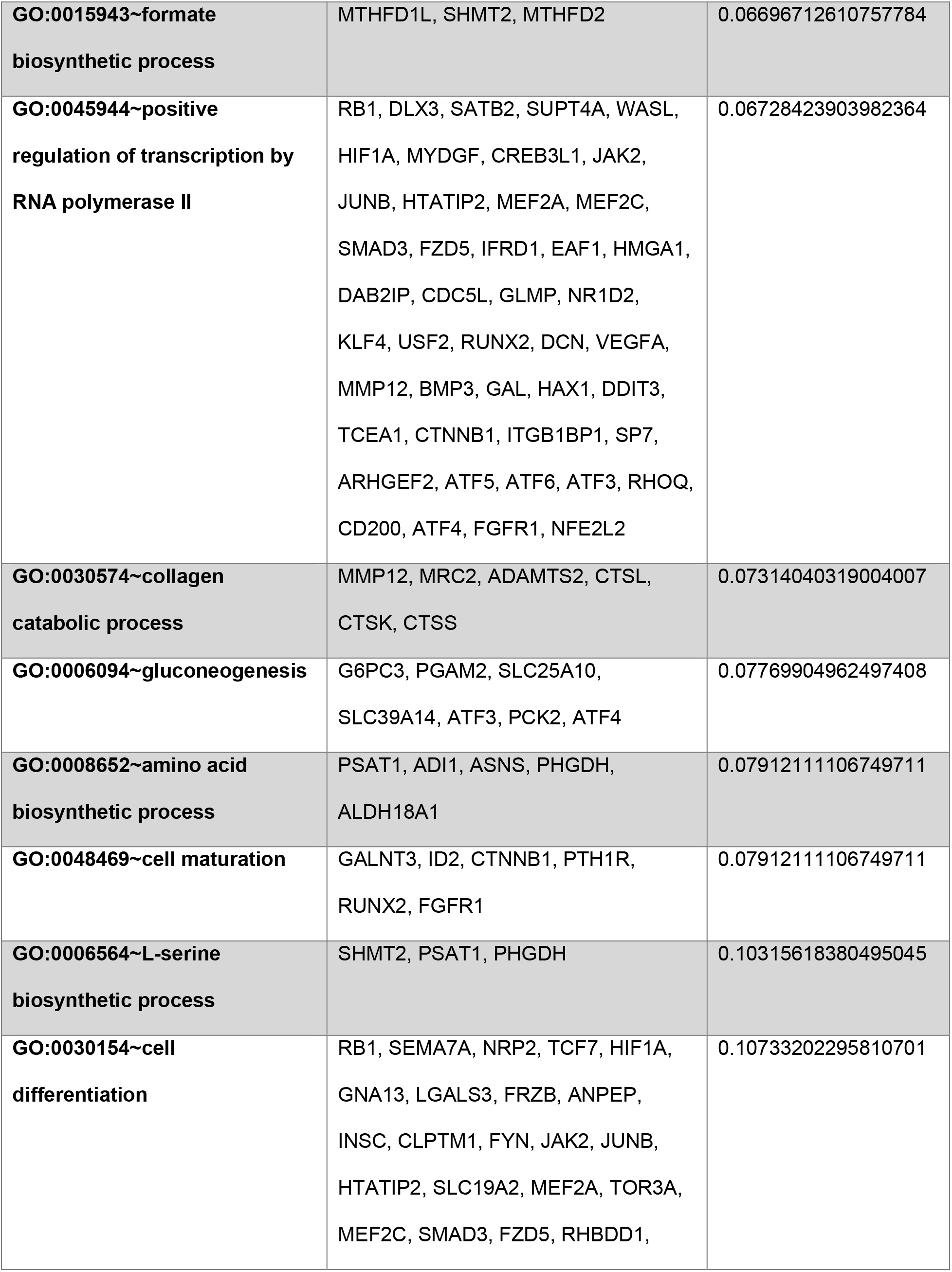
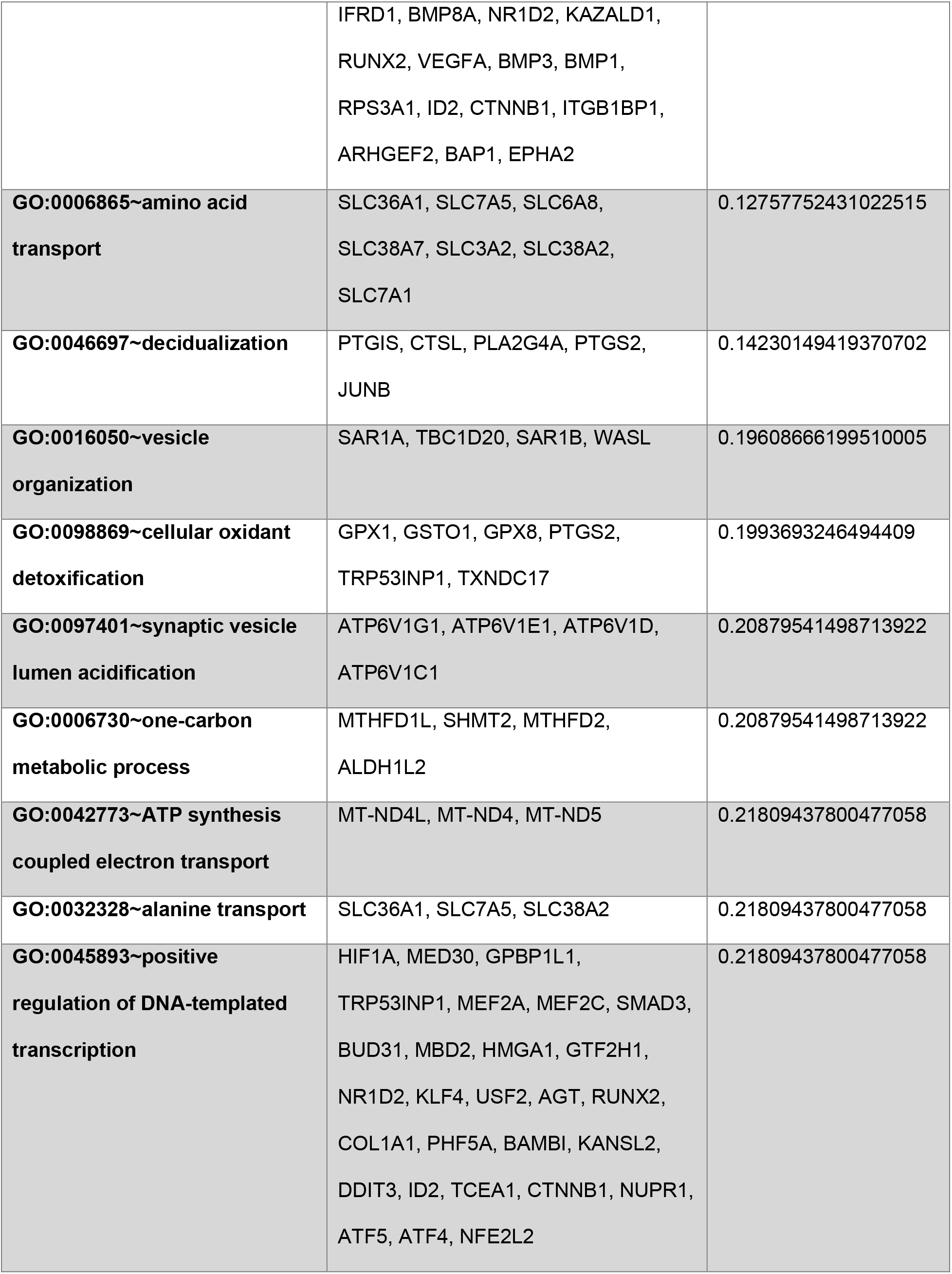
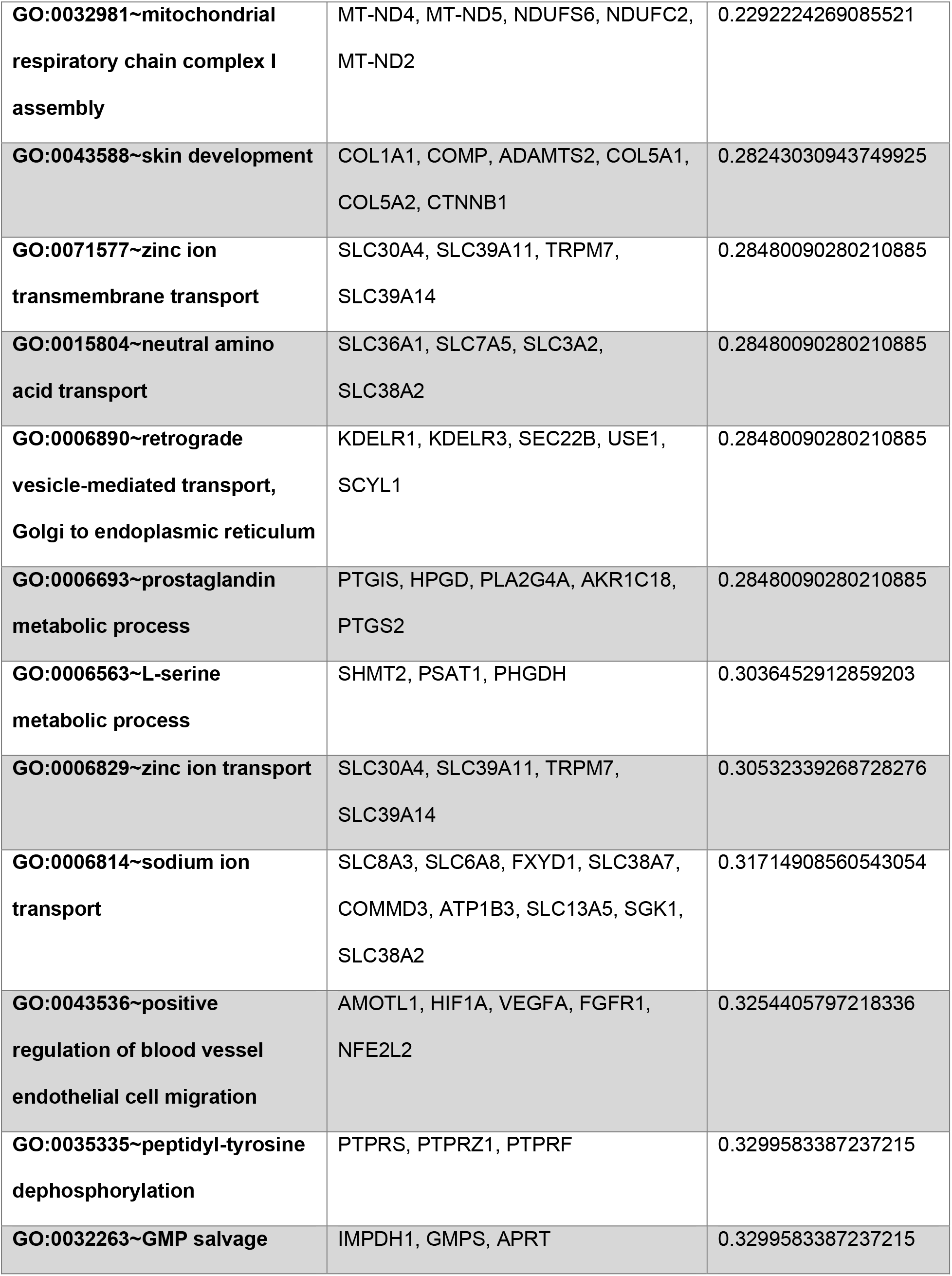
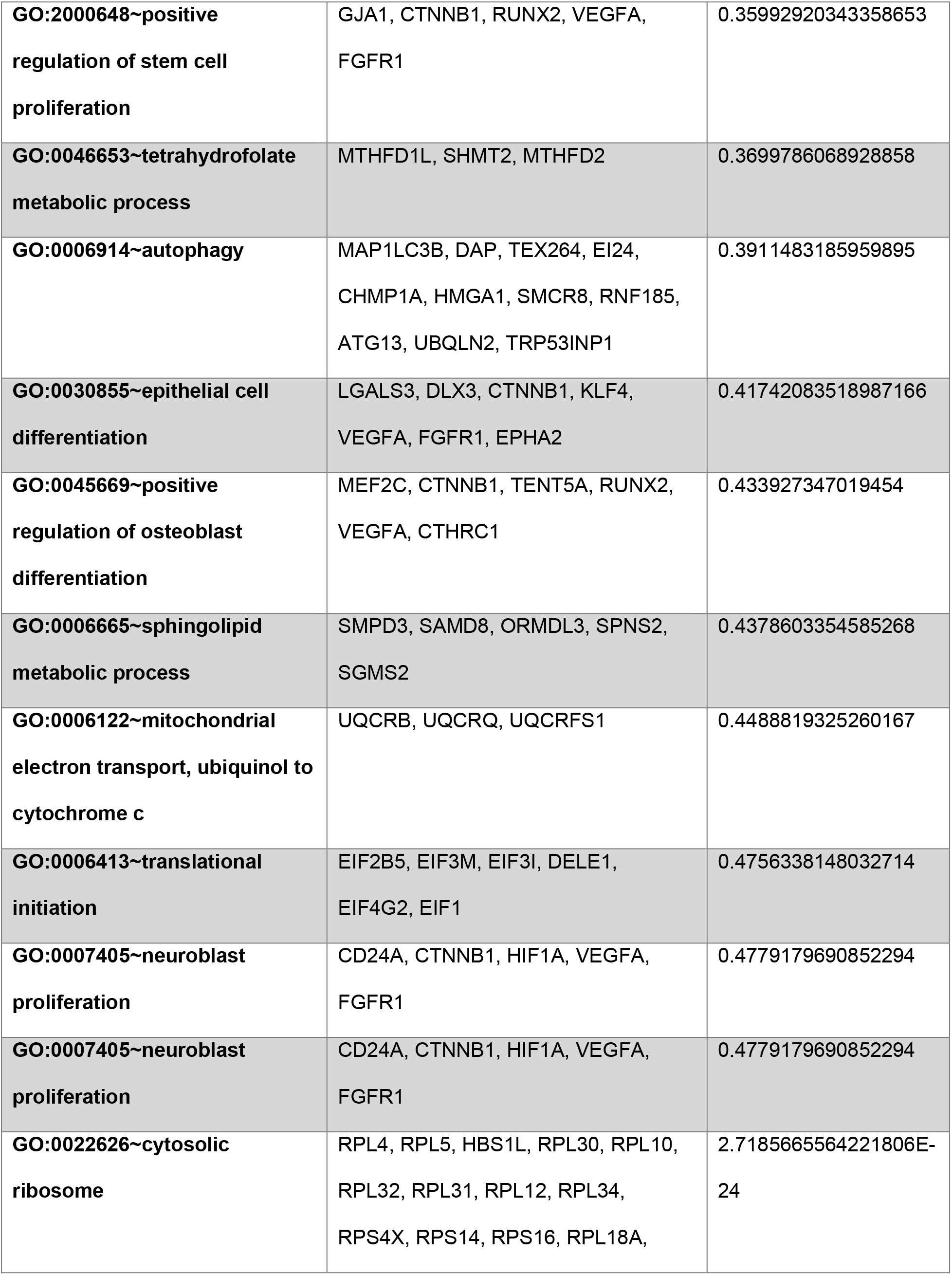
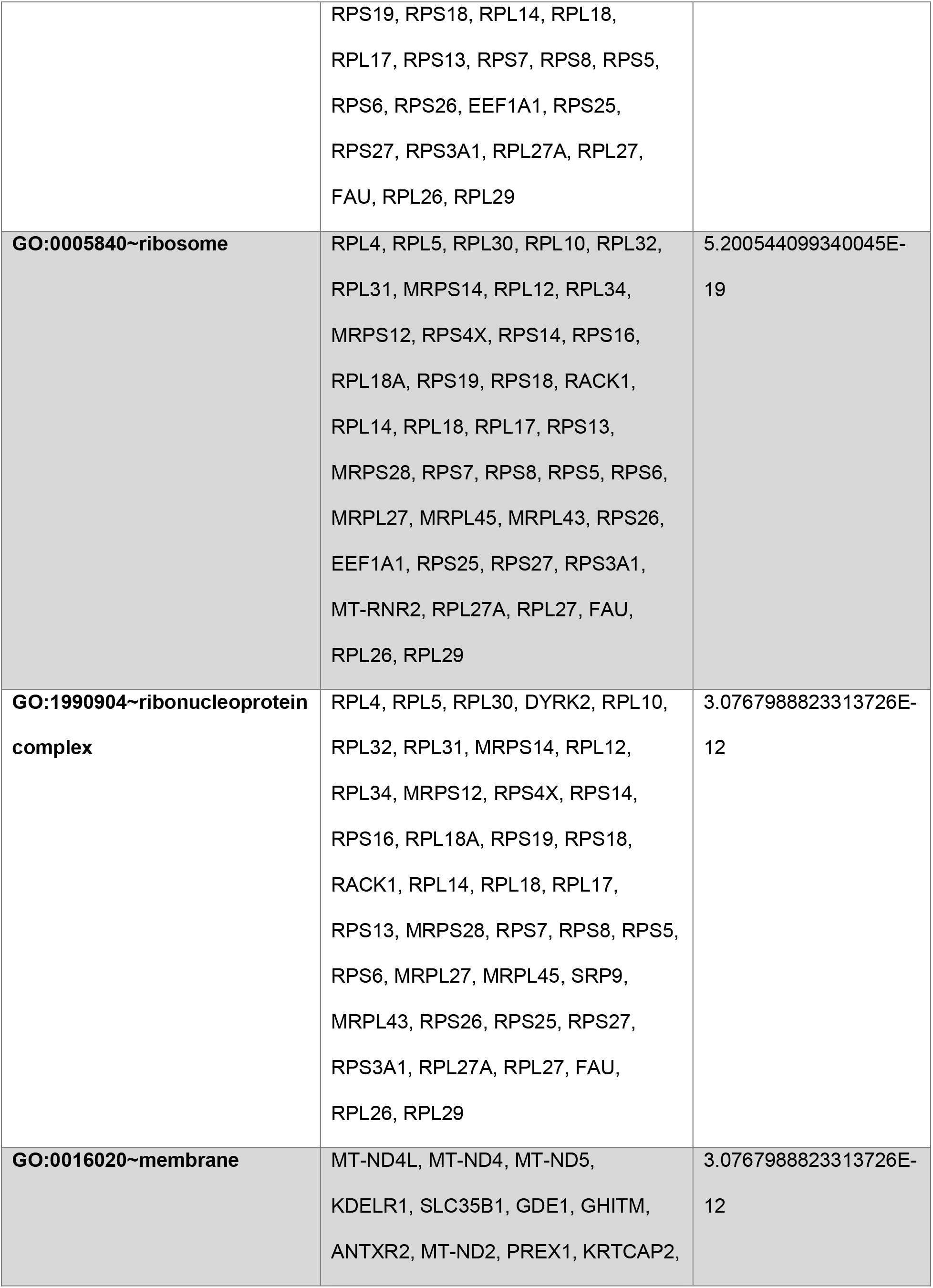
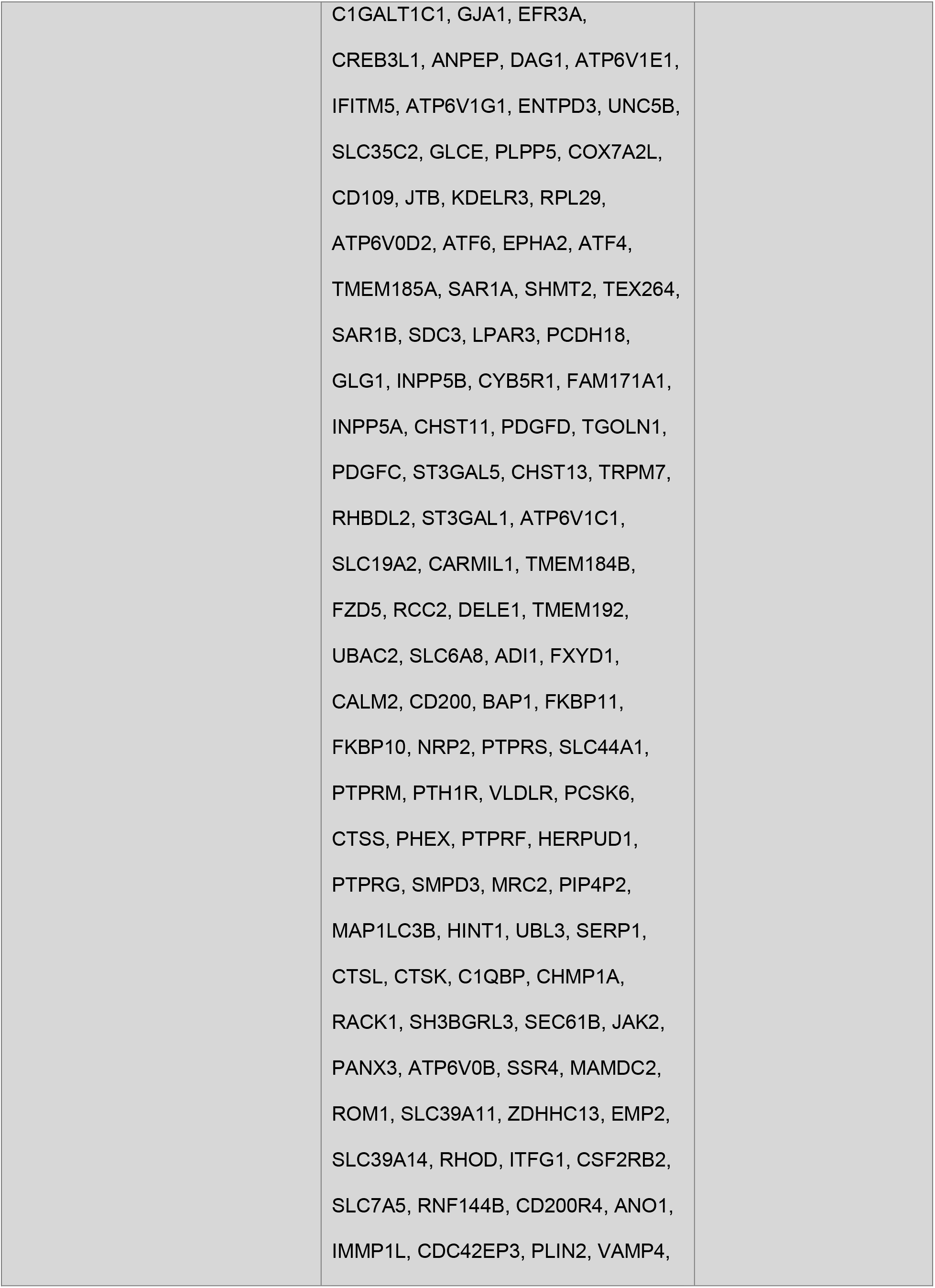
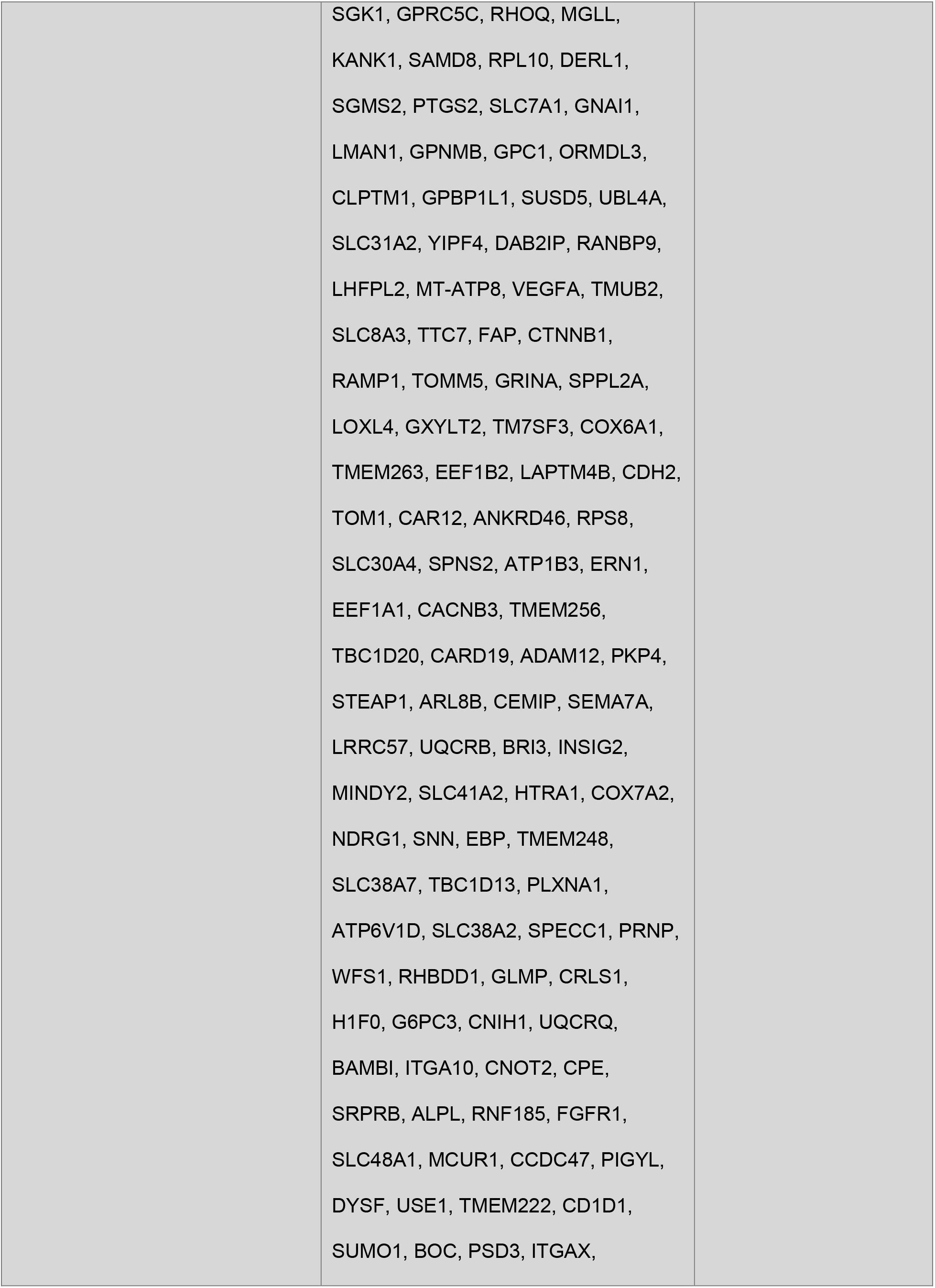
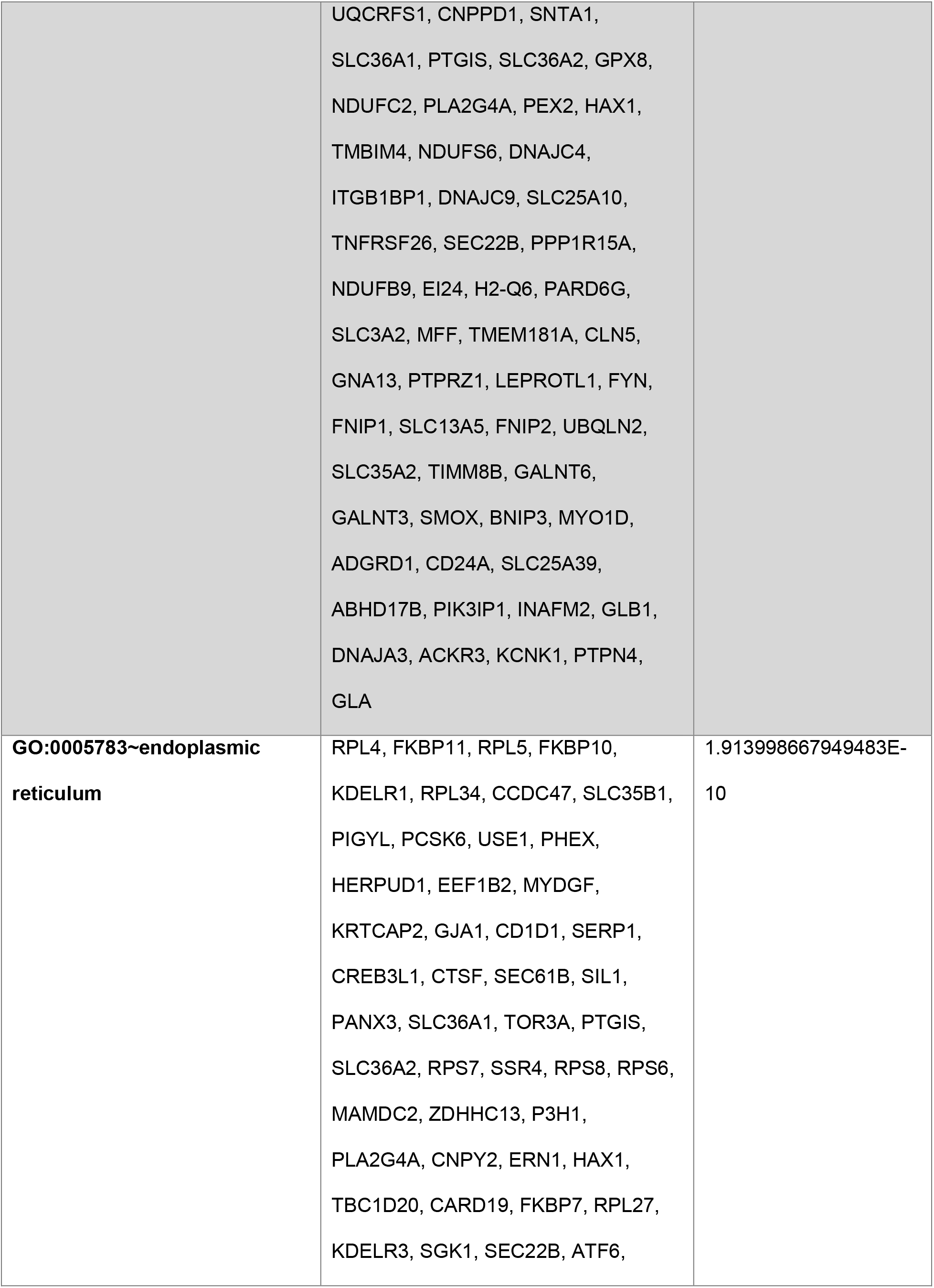
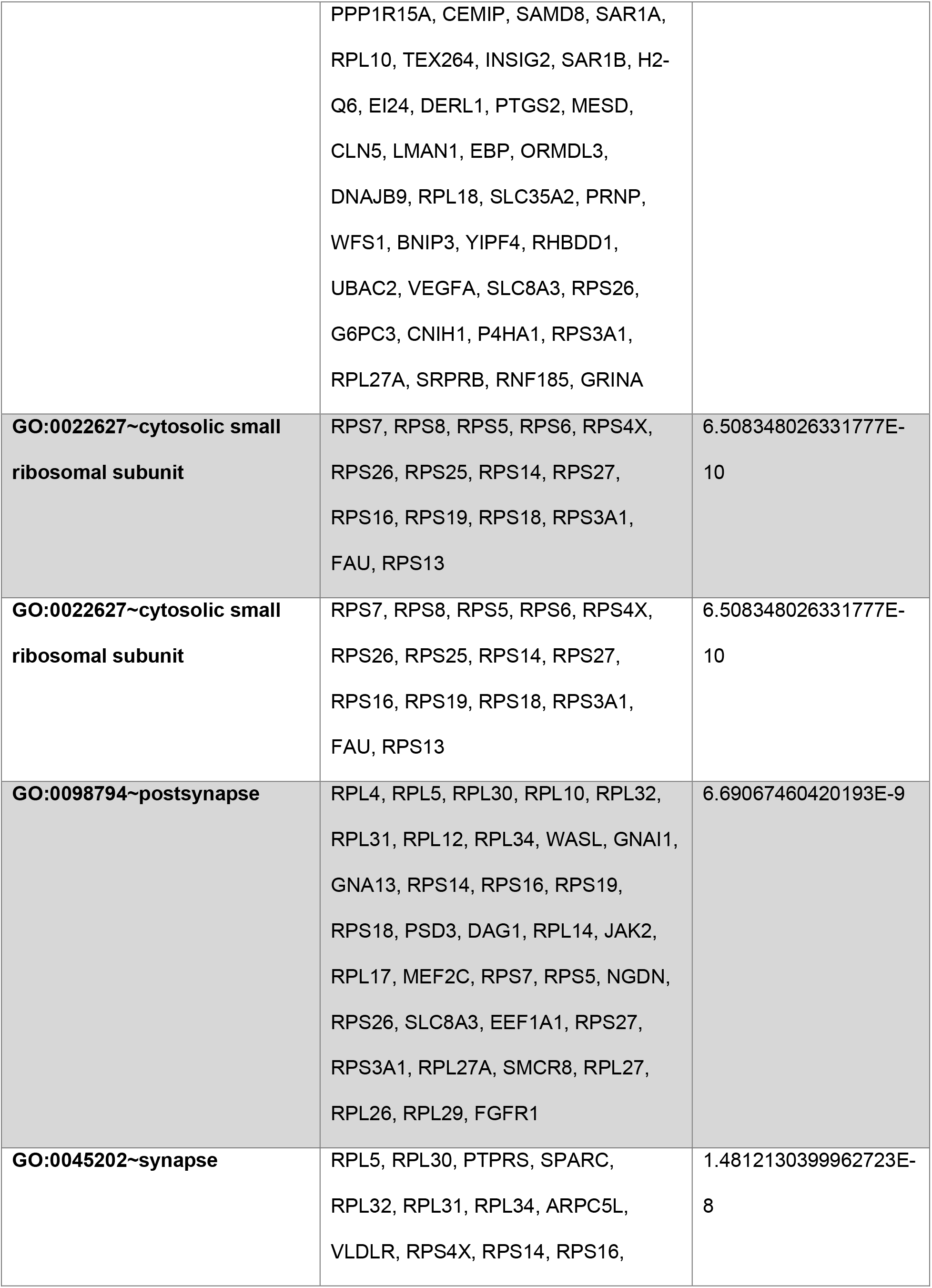
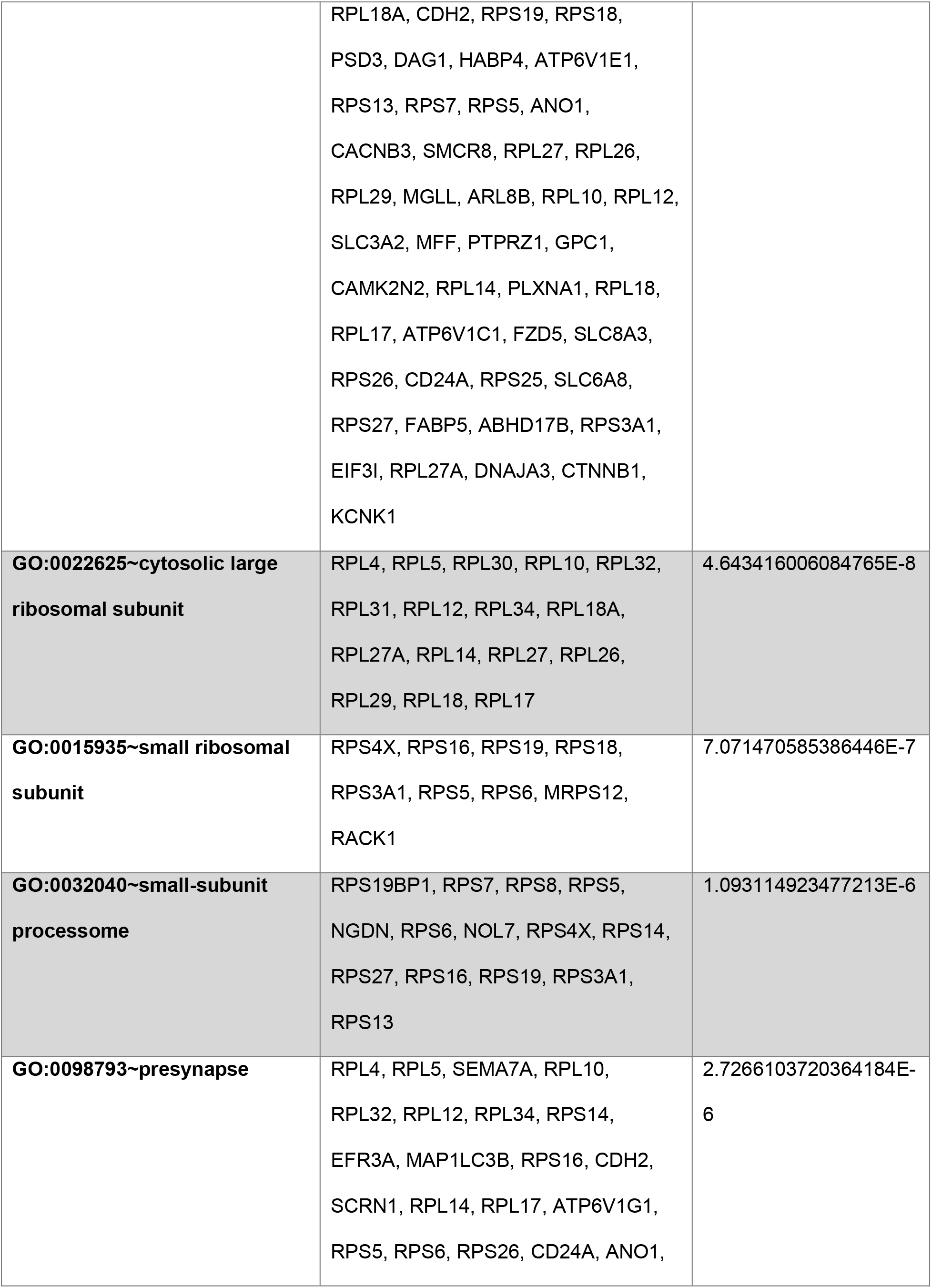
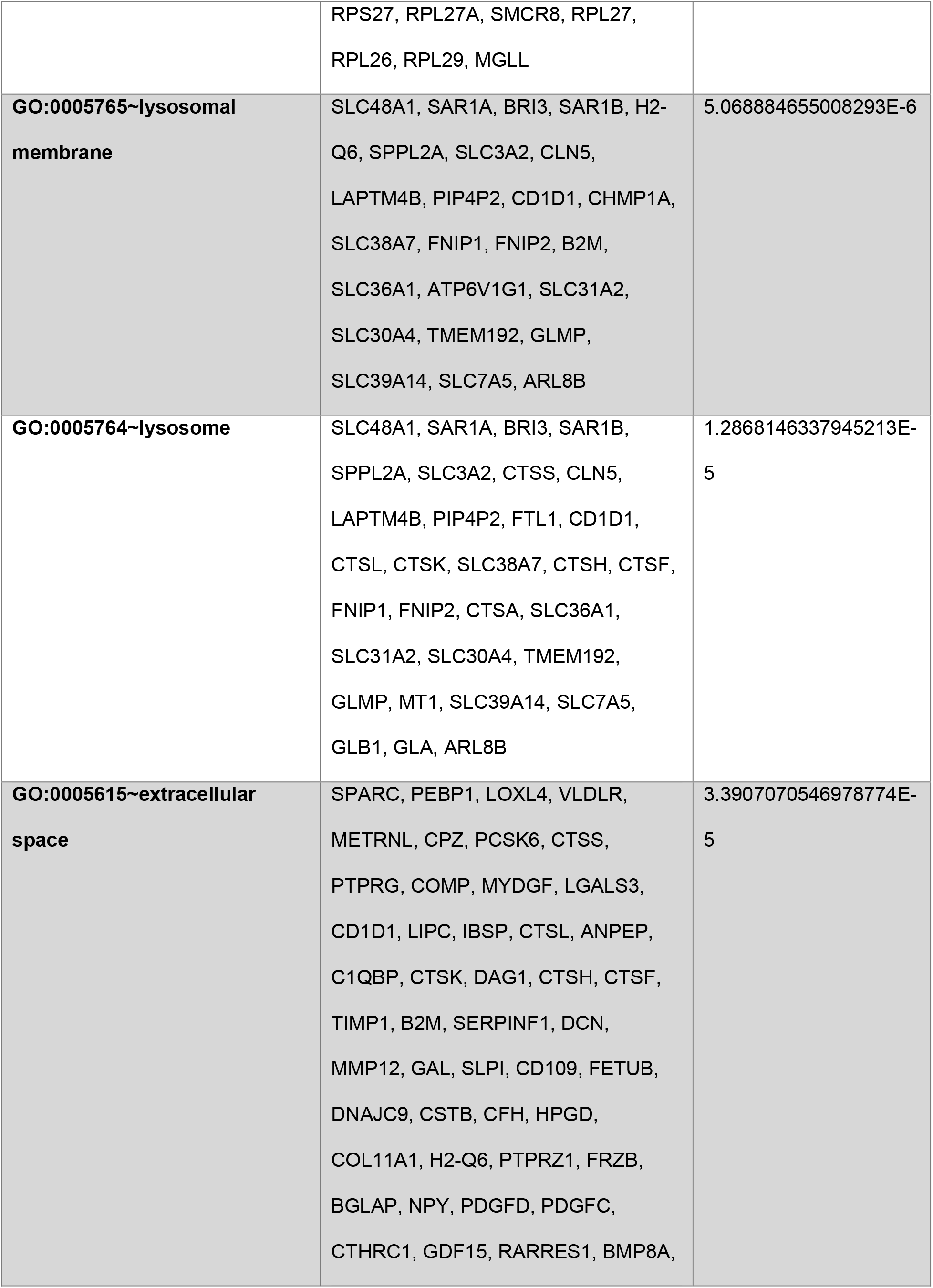
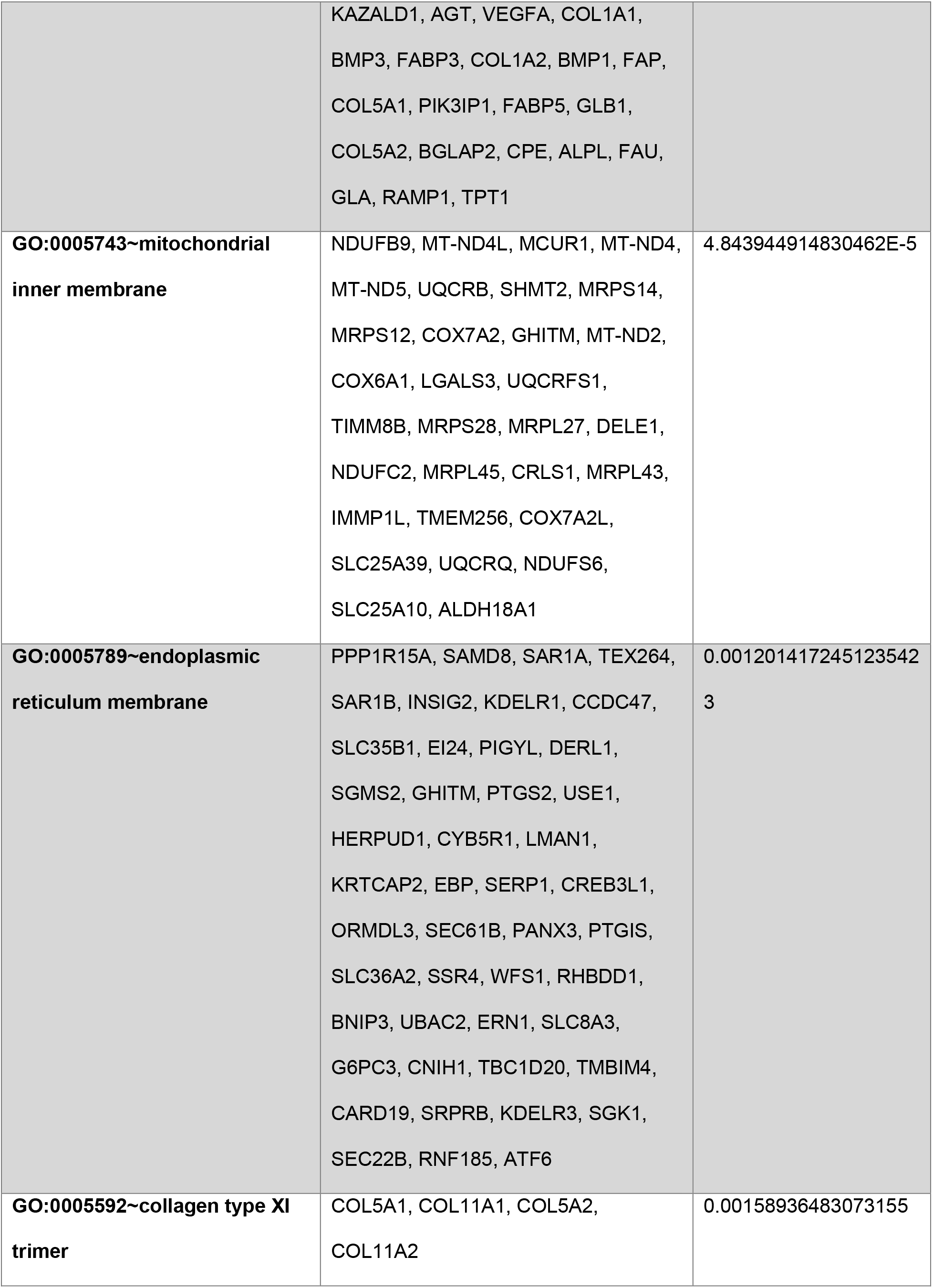
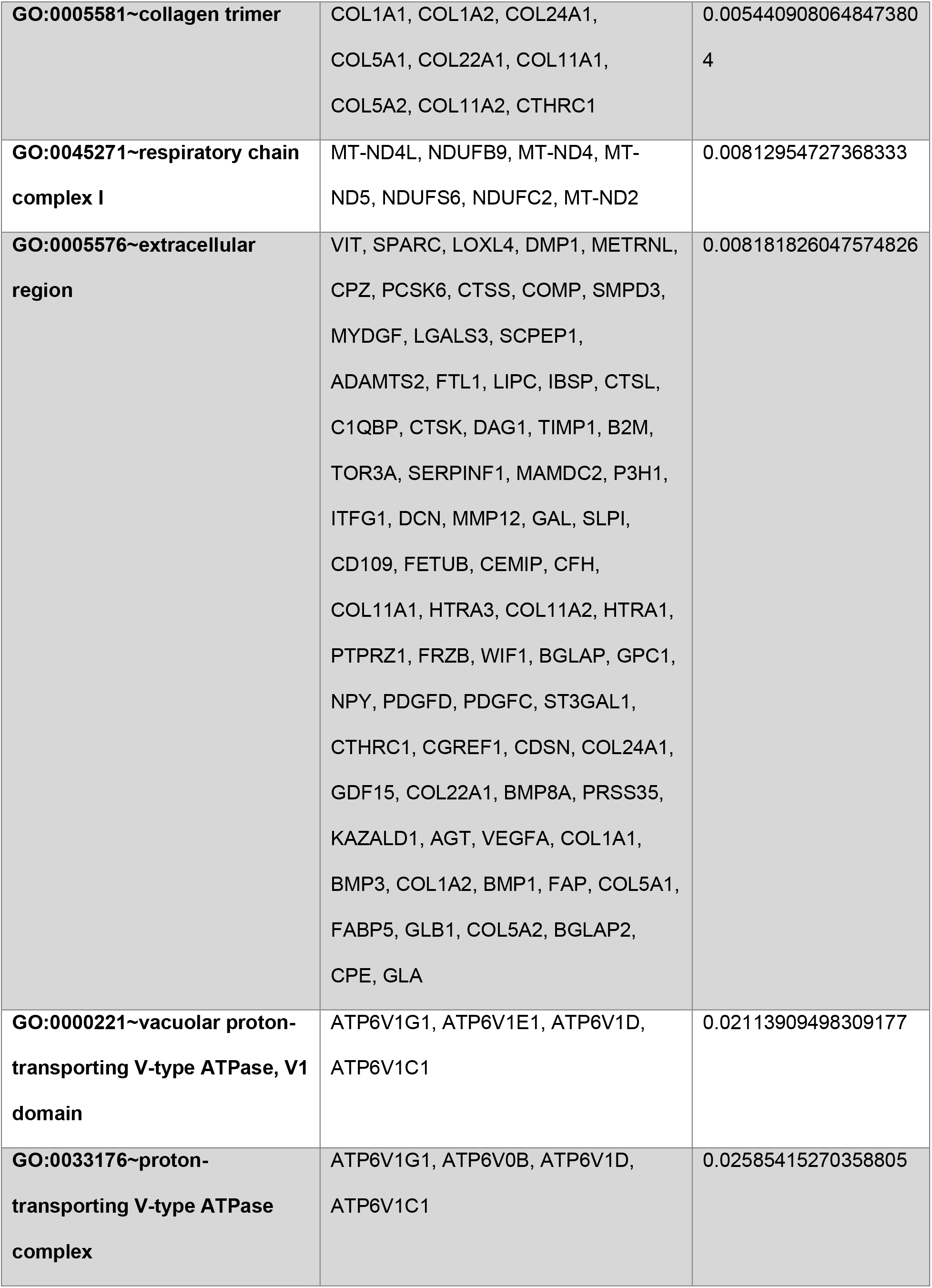
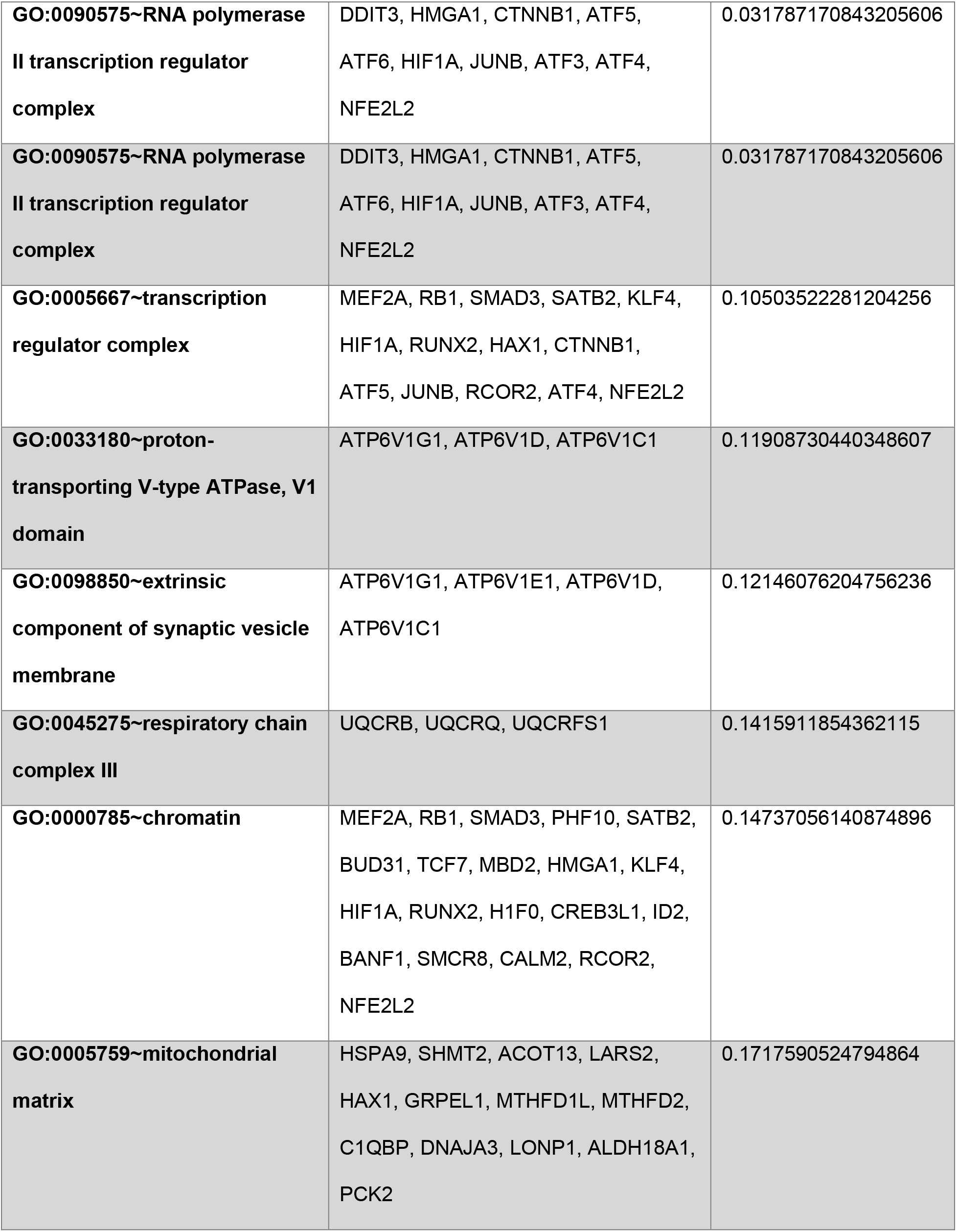
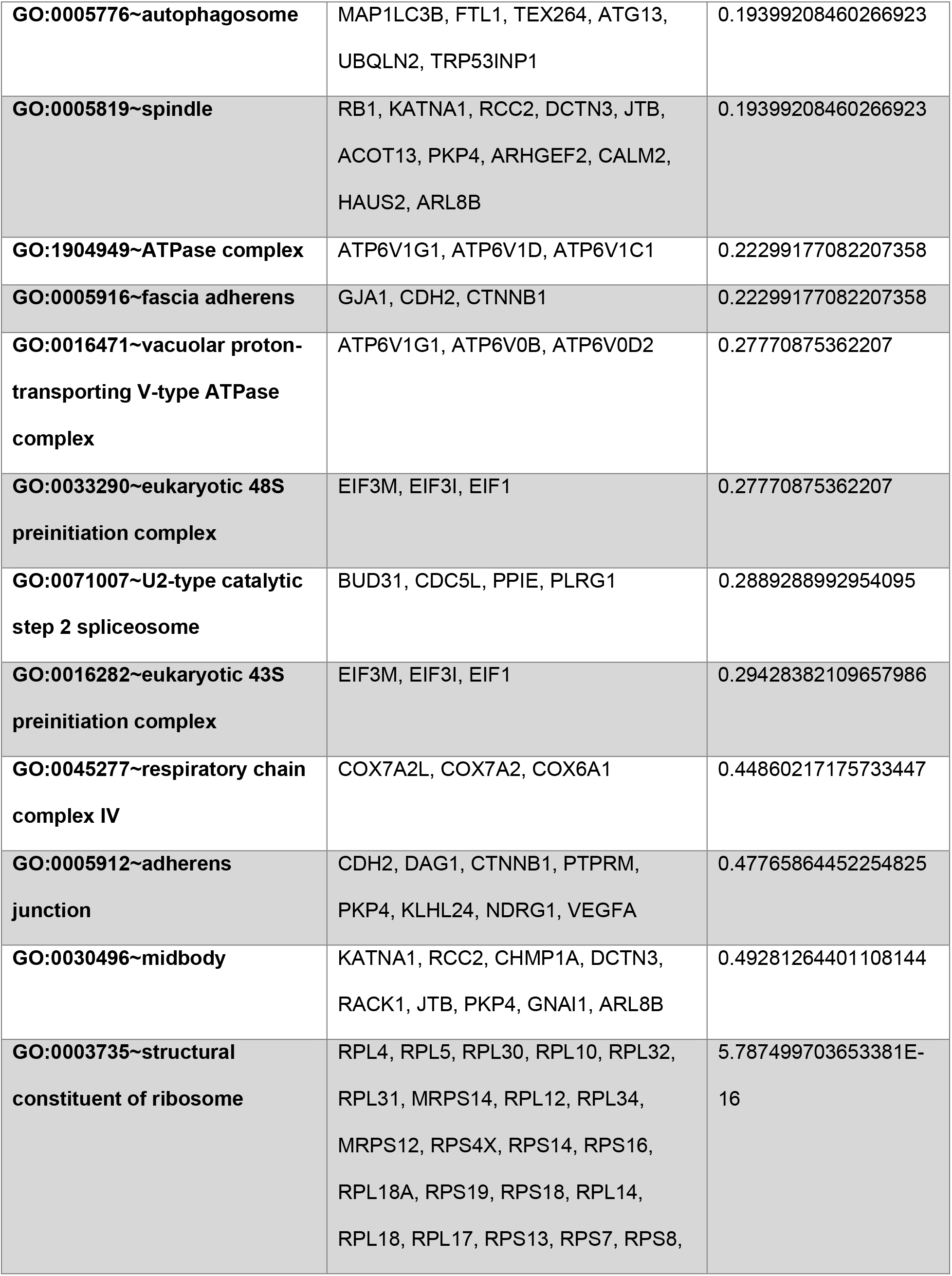
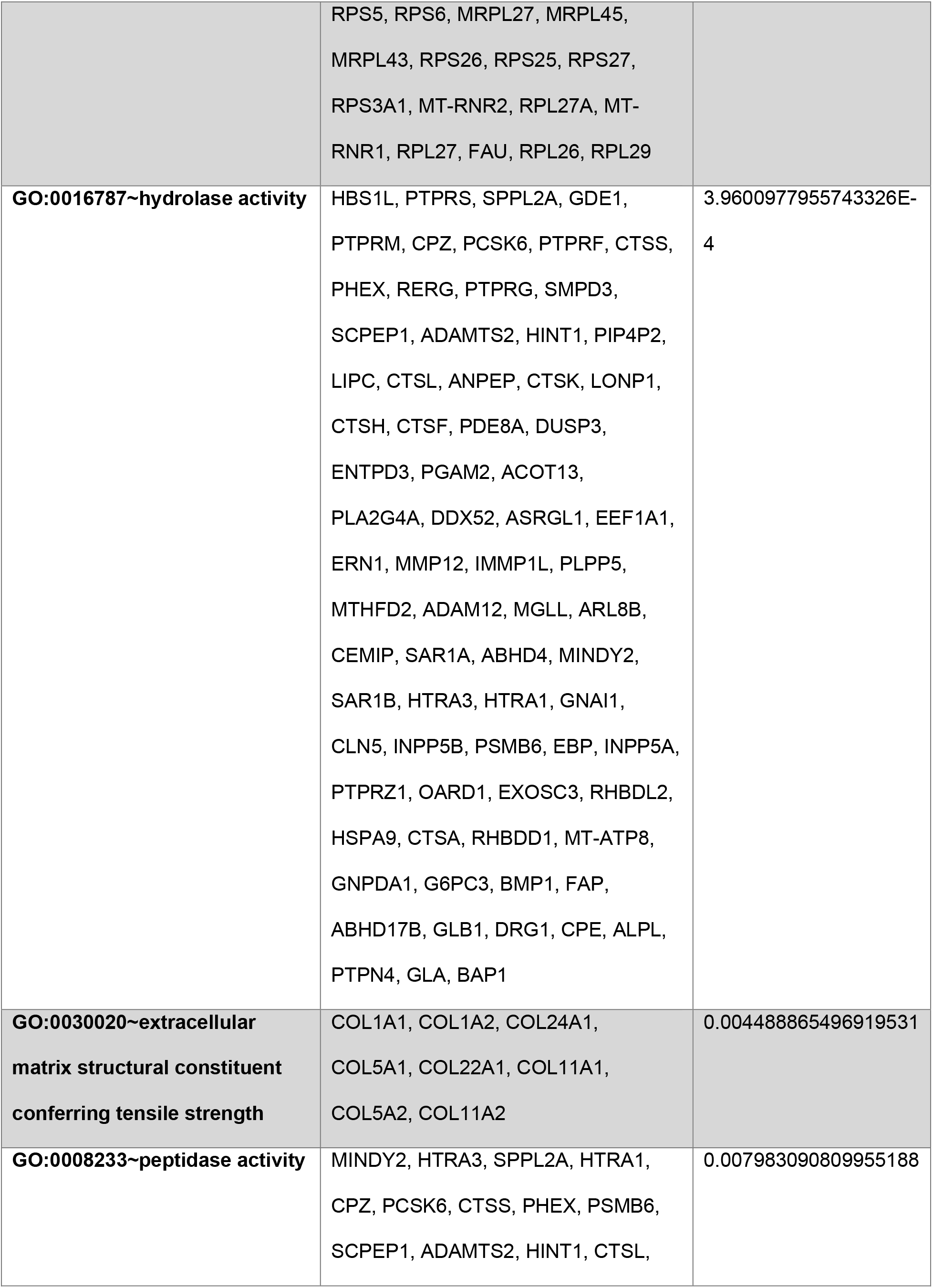
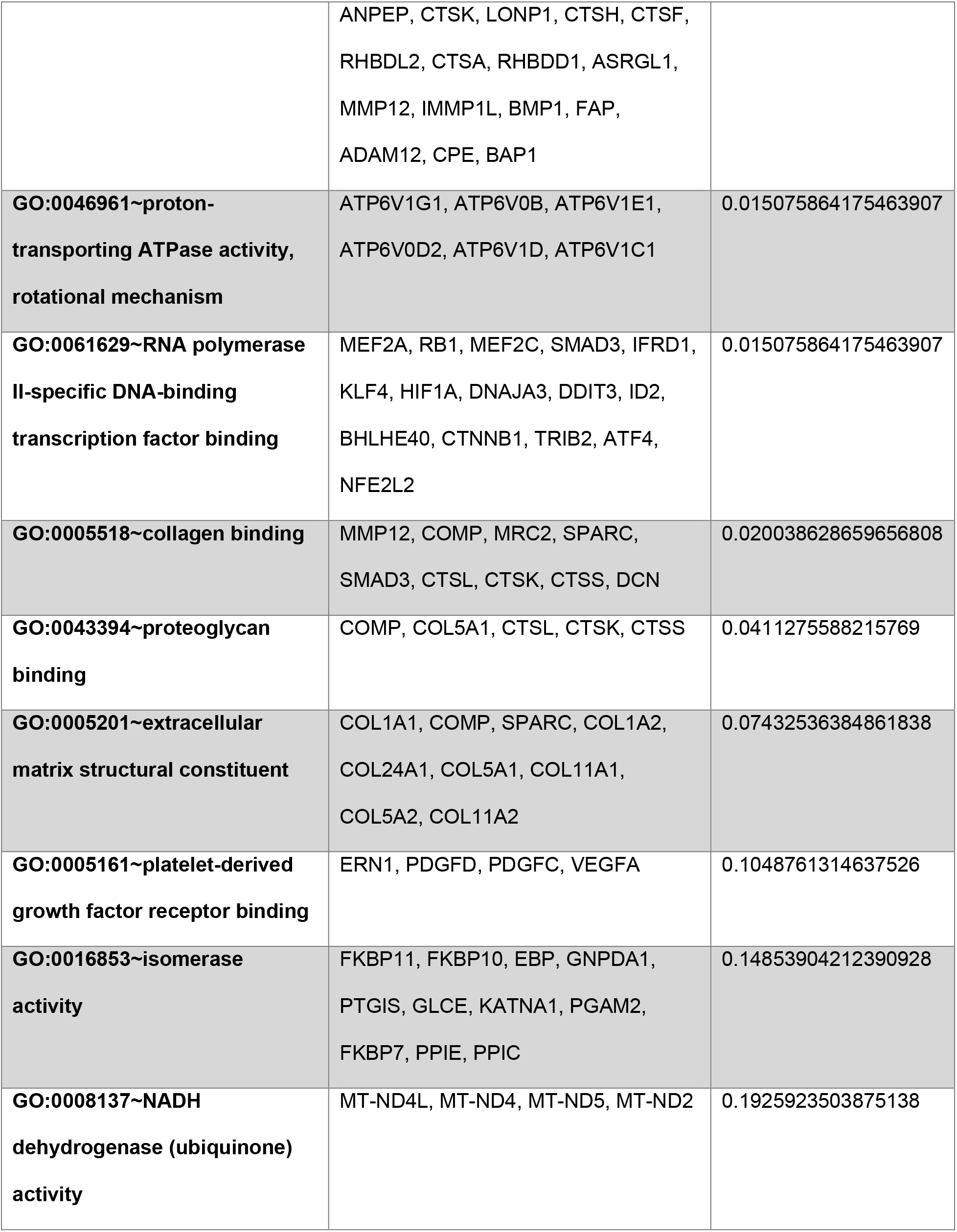
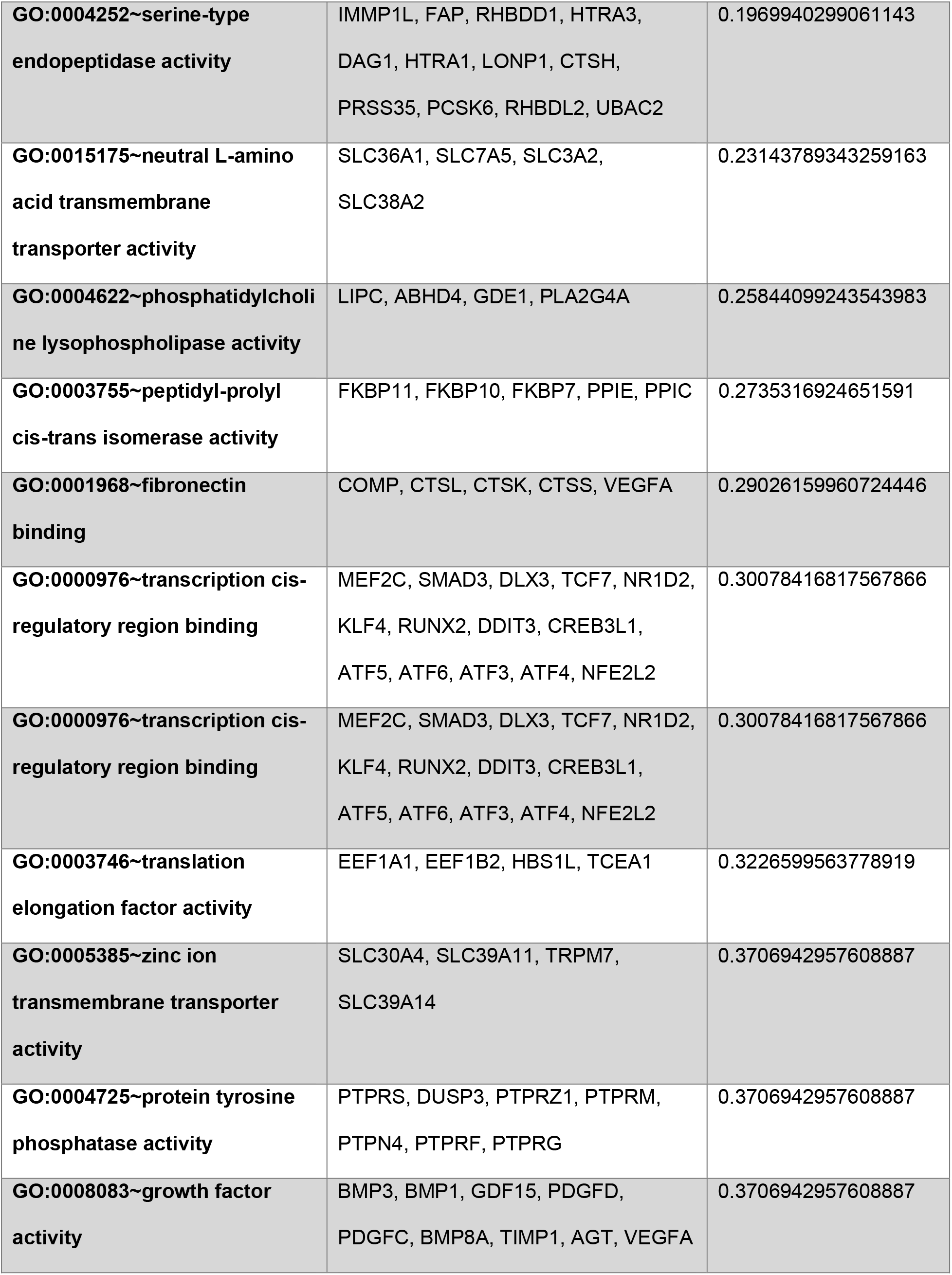
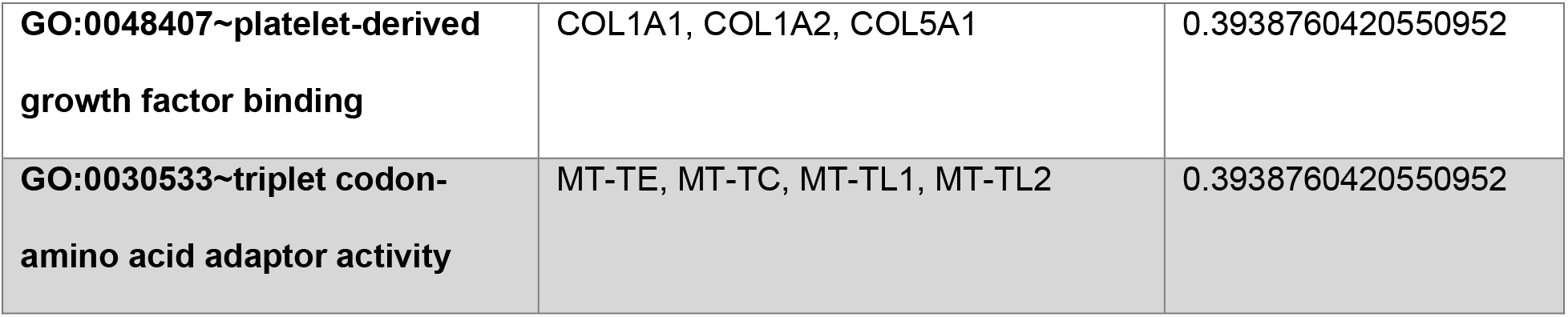
DAVID Gene Ontology (GO) enrichment analysis of genes upregulated in adipogenic medium following LIV treatment compared with untreated controls. Genes included in the analysis exhibited at least a 25% increase in expression in the +LIV group relative to the −LIV group.

